# NSD3 stabilizes nuclear compartmentalization and promotes megabase-scale chromatin interactions

**DOI:** 10.64898/2026.02.07.704091

**Authors:** Yi-Hung Chen, John R. Collette, Krupa Sampat, Ivaylo D. Yonchev, Catherine E. Hawkins, Christopher Ponne, Celeste D. Rosencrance, Kyle P. Eagen

## Abstract

Identifying biomolecules that shape nuclear organization is essential for understanding gene regulation in health and disease. Oncogenic fusion proteins rewire chromosome folding and generate biomolecular condensates, but the cofactors of oncoprotein-driven chromatin regulation remain poorly defined, and whether such factors have analogous functions in fusion-naïve cells is unknown. We find that NSD3 mediates chromosome folding in fusion-positive and fusion-negative cells. NSD3 stabilizes the BRD4-NUT fusion oncoprotein on chromatin, promotes histone H3K36me2, and supports oncogene expression while maintaining BRD4-NUT nuclear condensates. NSD3 loss attenuates distant chromatin interactions between BRD4-NUT megadomains both within and between chromosomes. In cells lacking BRD4-NUT, the short, catalytically inactive isoform of NSD3, NSD3short, promotes chromatin contacts separated by multiple megabases. The ability of NSD3short to promote long-range chromatin contacts requires its PWWP domain. By combining chromatin structural analyses in fusion-positive and fusion-negative cells, we show that interrogating fusion oncoprotein-driven chromosome misfolding reveals the multicomponent basis of nuclear compartmentalization and uncovers an adaptor protein that promotes chromatin contacts independent of enzymatic activity.

## INTRODUCTION

Chromosomes are the molecules that carry genes. Active and inactive genes are often positioned in different regions of the nucleus and are differentially folded^1^. Altered chromatin 3D structure in inherited and acquired disease misregulates gene expression, establishing a structure-function relationship between chromosome conformation and gene activity^2–6^. However, the molecules that fold chromosomes are yet to be fully defined.

Chromosome folding patterns vary across length scales. At the most fundamental level, DNA wraps around histones to form nucleosomes^7^. At a scale greater than nucleosomes, molecular motors, such as cohesin, extrude DNA loops^8–13^. Cohesin-mediated loop extrusion, which is stalled by CCCTC-binding factor (CTCF)^14–19^, has been visualized^20–22^ and engineered^23^ in living cells. At scales of entire chromosomes and whole nuclei, active and inactive loci preferentially associate within euchromatic and heterochromatic compartments^24^. Chromosome compartmentalization also organizes *cis*-regulatory elements into microcompartments^25^, suggestive of how chromosome folding influences gene activity^26,27^. Chromosome compartmentalization is independent of cohesin and CTCF^25,28–32^, therefore, factors that mediate compartmentalization remain mysterious.

Rare human diseases have been informative in implicating molecules that form megabase-scale chromatin interactions to compartmentalize chromosomes. BRD4-NUT is a fusion oncoprotein that drives NUT carcinoma, a poorly differentiated squamous malignancy and one of the most aggressive solid tumors^33^. BRD4-NUT promotes intrachromosomal interactions separated by dozens of megabases as well as interactions between different chromosomes^34^. These interactions form a nuclear subcompartment^34^, which appears in healthy cells as microcompartments^1,25^. BRD4-NUT also forms nuclear condensates^35–41^. Direct visualization of BRD4-NUT condensates by microscopy provides a physical representation of compartmentalization detected by chromosome conformation capture methods^34^, which infer chromosome folding by chemical cross-linking and proximity ligation^24^. Whether additional proteins function alongside BRD4-NUT to promote chromatin interactions remains unknown.

We leveraged BRD4-NUT-driven nuclear organization to identify factors that stabilize compartmentalization by promoting long-range chromatin contacts. We identify a protein that not only stabilizes fusion oncoprotein-driven nuclear compartmentalization but also promotes megabase-scale chromatin contacts in fusion-naïve cells. By identifying a previously unrecognized mediator of chromosome folding, we show that compartmentalization arises from multiple molecular components that function independent of enzymatic activity.

## RESULTS

### NSD3 is enriched in BRD4-NUT chromatin domains and localizes to BRD4-NUT nuclear condensates

To identify protein cofactors of BRD4-NUT-driven chromatin regulation, we reanalyzed mass spectrometry proteomics data of proteins that co-immunoprecipitated with BRD4-NUT after having been crosslinked to chromatin^42^. We quantified the enrichment of proteins that immunoprecipitated with BRD4-NUT from patient-derived 797TRex cells and from non-NUT carcinoma 293TRex cells induced to express epitope-tagged BRD4-NUT relative to input. NSD3 was enriched as a BRD4-NUT-interacting-chromatin-bound protein in both cellular contexts (**Figure 1A**). That NSD3 binds to the extraterminal domains of wild-type BRD4 and of BRD4-NUT substantiates this interaction^43–47^. Numerous additional proteins interact with BRD4-NUT. We prioritized NSD3 because NSD3 is fused to NUT (NSD3-NUT) in a subset of NUT carcinoma tumors^46,48,49^, providing genetic evidence from human patients that supports a biochemical role of NSD3 in BRD4-NUT-driven chromatin regulation.

**Figure 1.**
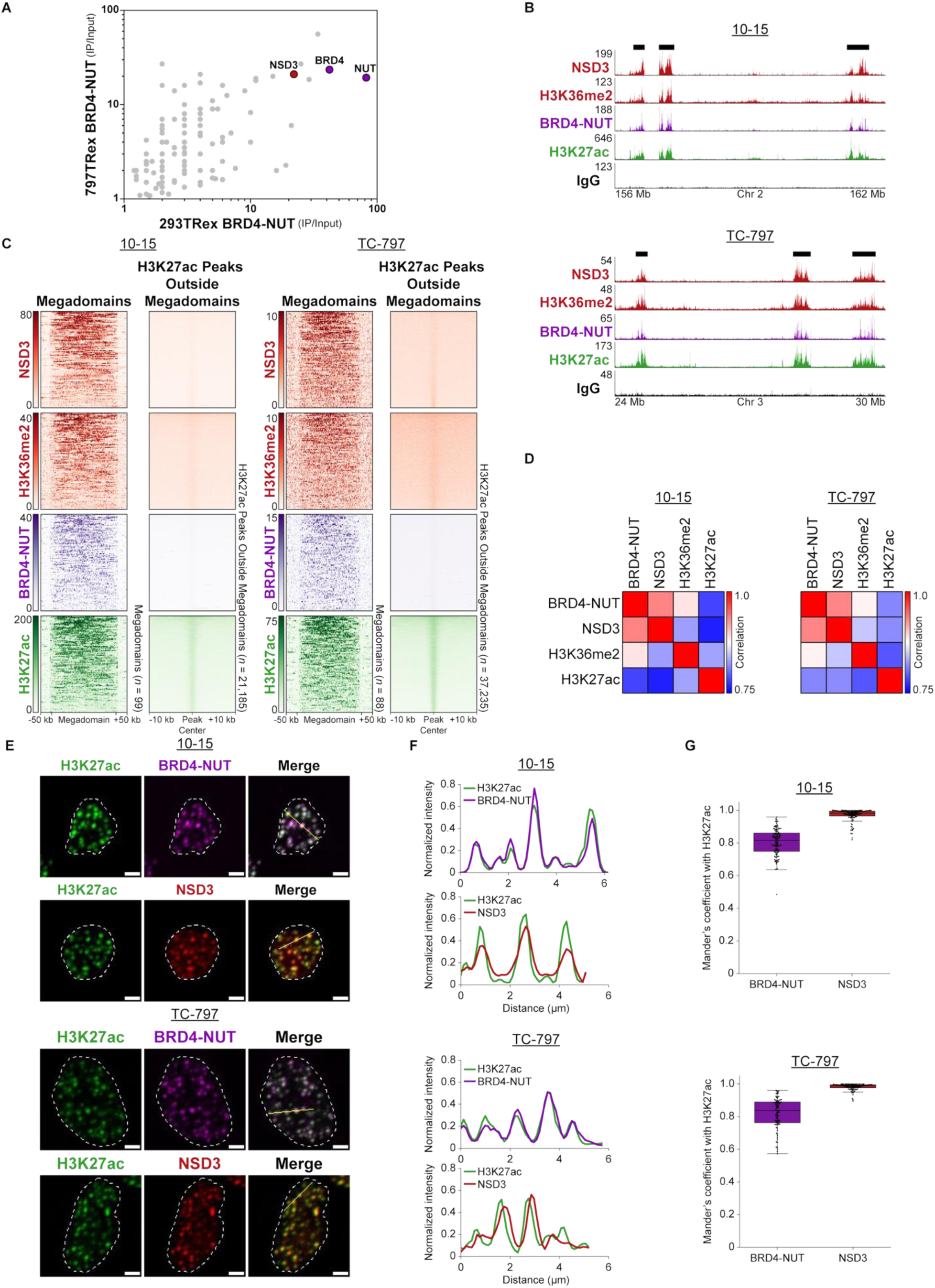
NSD3 and histone H3K36me2 are enriched in BRD4-NUT megadomains. (**A**) Scatterplot of protein interactions for BRD4-NUT determined by mass spectrometry proteomics. Enrichment for each protein in the immunoprecipitate relative to the input is plotted for 293TRex cells (x-axis) and for 797TRex cells (y-axis) induced to express epitope-tagged BRD4-NUT. NSD3 interacts with BRD4-NUT in both naïve (293TRex) and patient-derived (797TRex) cells. (**B**) CUT&RUN profiles for NSD3 (red), histone H3K36me2 (red), BRD4-NUT (purple), histone H3K27ac (green), and IgG (black) from patient-derived 10-15 and TC-797 cells endogenously expressing BRD4-NUT. Black bars indicate BRD4-NUT megadomains. (**C**) Heatmaps of the CUT&RUN signal for NSD3, histone H3K36me2, BRD4-NUT, and histone H3K27ac within each megadomain and at histone H3K27ac peaks outside megadomains in 10-15 and TC-797 cells indicate that NSD3 and histone H3K36me2 are strongly enriched in megadomains. Megadomains were normalized to the same length and 50 kb of flanking DNA is shown next to each normalized megadomain. H3K27ac peak centers were aligned and 10 kb of DNA flanking each H3K27ac peak is shown. IgG signal has been subtracted from each heatmap. (**D**) Pairwise Pearson correlations of CUT&RUN enrichment relative to IgG show that NSD3 and histone H3K36me2 enrichment are more strongly correlated with BRD4-NUT enrichment than histone H3K27ac enrichment in megadomains. (**E**) Immunofluorescence micrographs of histone H3K27ac (green), BRD4-NUT (purple), and NSD3 (red) show that, like BRD4-NUT, NSD3 overlaps with histone H3K27ac within BRD4-NUT condensates. Scale bars, 5 µm. (**F**) Profiles quantifying normalized fluorescence intensity along the yellow lines in the merged images from (E). Fluorescence intensity was normalized by dividing by the total fluorescence intensity across each yellow line. (**G**) Box plots (horizontal lines indicate medians; boxes indicate interquartile range; whiskers extend 1.5 times the interquartile range) of Mander’s coefficients of BRD4-NUT with histone H3K27ac and NSD3 with histone H3K27ac. Data points represent individual nuclei. 110 (10-15 cells) and 89 (TC-797 cells) nuclei were used to analyze the overlap of BRD4-NUT with H3K27ac. 106 (10-15 cells) and 95 (TC-797 cells) nuclei were used to analyze the overlap of NSD3 with H3K27ac.

NSD3 is a histone lysine methyltransferase that catalyzes di-methylation of histone H3 at lysine 36 (H3K36me2)^50–52^. We therefore evaluated the occupancy of NSD3 and H3K36me2 on chromatin relative to BRD4-NUT. BRD4-NUT forms exceptionally broad (100 kb to 2 Mb) hyperacetylated domains of chromatin, defined as megadomains^53^. We performed Cleavage Under Targets and Release Using Nuclease (CUT&RUN)^54^ with quantitative normalization^55^ to determine the localization patterns of NSD3, histone H3K36me2, BRD4-NUT, and histone H3K27ac across the genome in two NUT carcinoma patient-derived cell lines, 10-15^56^ and TC-797^57^, both of which endogenously express BRD4-NUT. The pattern of both NSD3 and H3K36me2 localization on chromatin mirrored that of BRD4-NUT (**Figure 1B**). NSD3 and H3K36me2 both showed robust enrichment within megadomains (**Figure 1C**). To assess whether NSD3 and histone H3K36me2 enrichment is specific to megadomains or extends to other genomic regions we analyzed the IgG-subtracted CUT&RUN enrichment at H3K27ac peaks outside megadomains. Compared to the strong enrichment observed within megadomains, NSD3, histone H3K36me2, and BRD4-NUT showed only weak enrichment at these regions genome-wide (**Figure 1C**). Additionally, pairwise Pearson correlation analysis of CUT&RUN tracks suggests that NSD3 and histone H3K36me2 exhibit stronger correlation with BRD4-NUT than histone H3K27ac (**Figure 1D**). NSD3 and histone H3K36me2 are not only integral to megadomains, but also more precisely define megadomains than H3K27ac.

BRD4-NUT and H3K27ac localize to discrete nuclear foci visible by immunofluorescence microscopy in 10-15 and TC-797 cells (**Figure 1E**). The overlapping intensity profiles of the fluorescence signal for BRD4-NUT and for H3K27ac indicate localization of chromatin modified with histone H3K27ac within BRD4-NUT condensates (**Figure 1F**). To assess whether NSD3 colocalizes with BRD4-NUT condensates, we also stained 10-15 and TC-797 cells for NSD3. Due to antibody incompatibility, we were unable to co-stain for both BRD4-NUT and NSD3 in the same cells and therefore used H3K27ac as a surrogate for BRD4-NUT condensates, as done previously^34,37–41,53^. NSD3 also localizes to distinct nuclear foci that are similar to BRD4-NUT condensates and that colocalize with H3K27ac, indicating that NSD3 is a component of BRD4-NUT condensates (**Figures 1E and F**). To quantitatively assess the colocalization of NSD3 with BRD4-NUT across the population of cells we determined the Mander’s coefficient^58^, which quantifies the degree of colocalization between two fluorescence signals by calculating the fraction of one signal overlapping with the other (**Table S1**). For 10-15 and TC-797 cells, the median Mander’s coefficients of NSD3 with H3K27ac were 0.983 and 0.988, respectively, and the median Mander’s coefficients of BRD4-NUT with H3K27ac were 0.815 and 0.838, respectively (**Figure 1G**). Mander’s coefficient analyses indicate that 98% of NSD3 and greater than 80% of BRD4-NUT overlap with H3K27ac within the nucleus in both cell lines. Because the Mander’s coefficient is directional we also assessed colocalization by reversing the order in which we analyzed the data. The median Mander’s coefficients of H3K27ac with BRD4-NUT were 0.871 and 0.793 in 10-15 and TC-797 cells, respectively, suggesting that H3K27ac nearly completely overlaps with BRD4-NUT (**Figure S1**). The median Mander’s coefficients of H3K27ac with NSD3 were 0.616 and 0.426 in 10-15 and TC-797 cells, respectively, indicating that despite NSD3 almost entirely overlapping with H3K27ac, not all H3K27ac colocalizes with NSD3 (**Figure S1**). Our immunofluorescence analysis demonstrating that NSD3 localizes to BRD4-NUT condensates indicates that BRD4-NUT condensates are multicomponent. Together with the CUT&RUN analysis, these findings implicate NSD3 as a cofactor with BRD4-NUT to mediate BRD4-NUT-driven nuclear organization.

### NSD3 regulates the composition of megadomains

To test a role for NSD3 in BRD4-NUT-driven nuclear organization, we depleted NSD3 using CRISPR interference (CRISPRi)^59^. We stably expressed the Zim3-dCas9 CRISPRi transcriptional repressor^60^ in two NUT carcinoma cell lines, 10-15, and TC-797. We validated our CRISPRi system by depleting the non-essential cell surface protein CD81. Quantifying CD81 levels by flow cytometry indicated robust CD81 depletion (99.7% and 99.8% knockdown efficiency for 10-15 and TC-797, respectively) in the vast majority (97.6% and 99.2% of 10-15 and TC-797 cells depleted, respectively) of both cell populations (**Figure S2**). This confirmed the robust efficacy of Zim3-dCas9 and the feasibility of applying CRISPRi to uniformly deplete NSD3 from 10-15 and TC-797 cells. To deplete NSD3, we transduced Zim3-dCas9-expressing cells with a dual-sgRNA lentiviral construct^61^ targeting the transcription start site of NSD3 (sgNSD3). Immunoblotting showed that NSD3 was depleted to 13.1% of endogenous levels in 10-15 cells (**Figures 2A and B**) and to 28.3% of endogenous levels in TC-797 cells (**Figures S4A and S4B**). We evaluated how depletion of NSD3 affects NUT carcinoma cell viability. Depletion of NSD3 moderately reduced the proliferation of 10-15 and TC-797 cells (**Figure S3A**) and mitigated the blockage of cell differentiation (**Figures S3B and S3C**), consistent with a previous report^46^. Enlarged and flattened cell morphologies, along with elevated levels of the epithelial-specific differentiation marker keratin 7 following NSD3 depletion confirmed cellular differentiation (**Figures S3B and S3C**).

**Figure 2.**
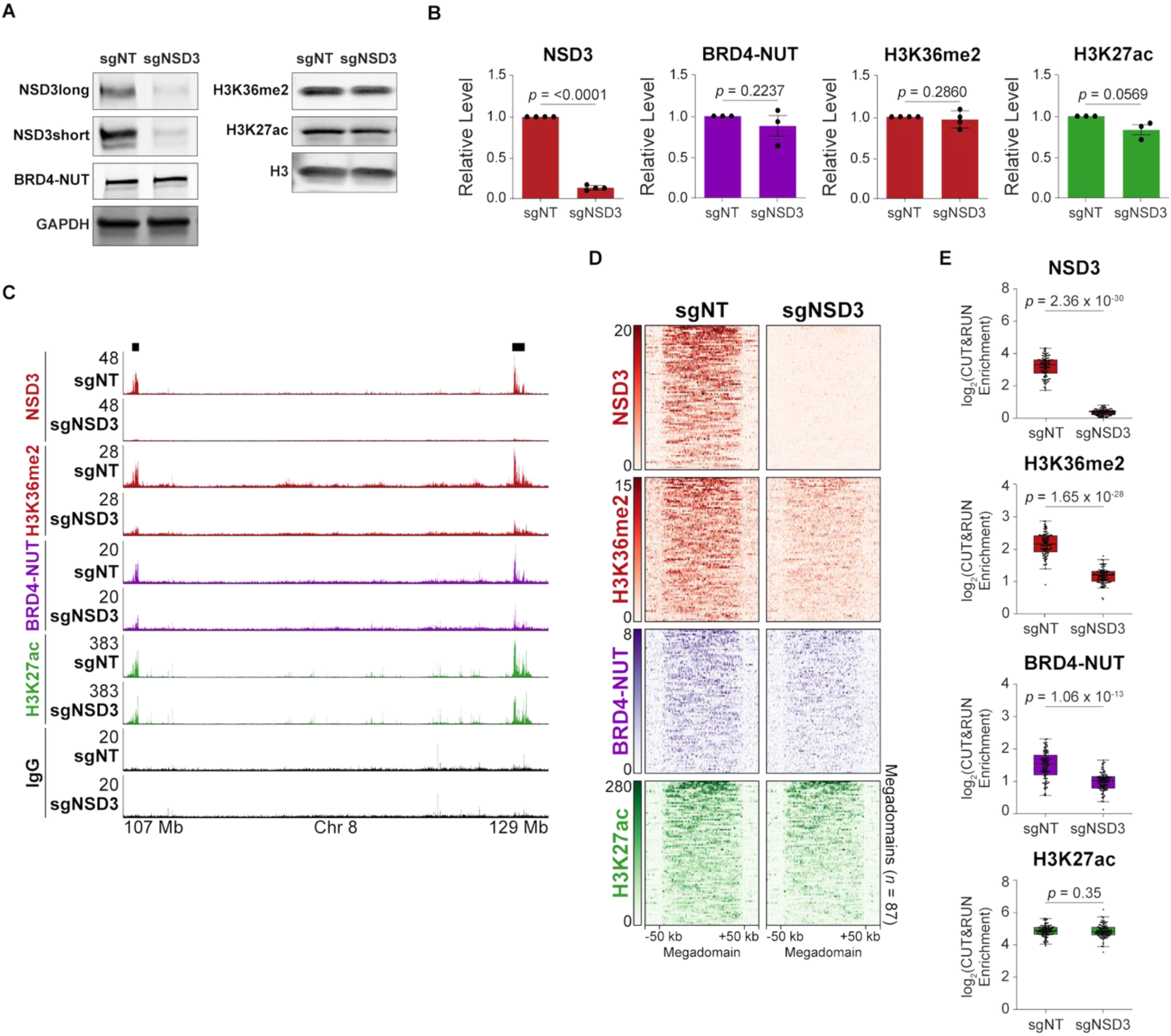
NSD3 deposits H3K36me2 in megadomains and stabilizes BRD4-NUT on chromatin. (**A**) Immunoblots indicate that NSD3 (long and short isoforms) levels are reduced, whereas BRD4-NUT, histone H3K36me2, and histone H3K27ac levels in 10-15 cells remain unchanged upon CRISPRi-mediated depletion of NSD3 (sgNSD3). A non-target sgRNA (sgNT) was used as a control. GAPDH and histone H3 were used as internal controls for non-histone proteins and histone modifications, respectively. (**B**) Quantification of the immunoblots from (A). NSD3short and NSD3long were combined to obtain the level of NSD3. Data points represent biological replicates. Bar and error bars represent the mean and standard error of the mean, respectively. p values determined using paired t-tests. (**C**) CUT&RUN profiles for NSD3 (red), histone H3K36me2 (red), BRD4-NUT (purple), histone H3K27ac (green), and IgG (black) from control (sgNT) and NSD3-depleted (sgNSD3) 10-15 cells. Chromatin occupancy of histone H3K36me2 and BRD4-NUT within megadomains (black bars) is reduced upon depletion of NSD3. (**D**) Heatmaps of the CUT&RUN signal for NSD3, histone H3K36me2, BRD4-NUT, and histone H3K27ac within megadomains in control (sgNT) and NSD3-depleted (sgNSD3) 10-15 cells. Megadomains were normalized to the same length and 50 kb of flanking DNA is shown next to each normalized megadomain. IgG signal has been subtracted from each heatmap. (**E**) Box plots (horizontal lines indicate medians; boxes indicate interquartile range; whiskers extend 1.5 times the interquartile range) indicate that NSD3, histone H3K36me2, and BRD4-NUT, but not histone H3K27ac are reduced within megadomains in NSD3-depleted (sgNSD3) compared to control (sgNT) 10-15 cells. Data points represent the average IgG-normalized CUT&RUN signals within megadomains from two biological replicates. p values determined using one-sided Mann-Whitney *U* tests.

Depletion of NSD3 did not significantly change global levels of H3K36me2 and BRD4-NUT and only slightly diminished global H3K27ac levels in 10-15 (**Figures 2A and 2B**) and TC-797 cells (**Figures S4A and S4B**). Next, we performed CUT&RUN to determine if depletion of NSD3 changed the occupancy of NSD3, H3K36me2, BRD4-NUT, and H3K27ac on chromatin. Depletion of NSD3 significantly decreased the enrichment of H3K36me2 in megadomains by 1.78-fold and 2.18-fold in 10-15 and TC-797 cells, respectively (**Figures 2C, 2D, 2E, S4C, S4D, and S4E**). Depletion of NSD3 significantly reduced the enrichment of BRD4-NUT in megadomains by 1.50-fold but did not significantly alter levels of histone H3K27ac in megadomains in 10-15 cells (**Figures 2C, 2D, and 2E**). Similar results were observed in TC-797 cells, where BRD4-NUT enrichment in megadomains was significantly reduced following NSD3 depletion. Although we detected a 1.19-fold decrease in H3K27ac enrichment within megadomains in TC-797 cells (**Figures S4C, S4D, and S4E**), the reduction was notably less pronounced than the 2.25-fold decrease in BRD4-NUT enrichment (**Figure S4E**). Taken together, these findings suggest NSD3 plays a critical and specific role in controlling the composition of megadomains.

To determine whether NSD3 selectively stabilizes BRD4-NUT chromatin binding within megadomains or across the genome, we examined how depletion of NSD3 affects regions outside megadomains. Specifically, we focused on H3K27ac peaks overlapping BRD4-NUT peaks, so-called H3K27ac-BRD4-NUT peaks, located outside megadomains; loci that reflect BRD4-NUT binding at defined locations rather than spreading across chromatin. NSD3 is enriched at H3K27ac-BRD4-NUT peaks outside megadomains in both cell lines (**Figure S5**). In both cell lines, NSD3 depletion slightly reduced the levels of BRD4-NUT at H3K27ac-BRD4-NUT peaks outside megadomains. The depletion of NSD3 had no impact on the levels of H3K27ac at H3K27ac-BRD4-NUT peaks outside megadomains. H3K36me2 is enriched at H3K27ac-BRD4-NUT peaks outside megadomains in 10-15 cells but is less enriched in TC-797 cells. NSD3 depletion led to a widespread reduction of H3K36me2 levels at H3K27ac-BRD4-NUT peaks outside megadomains in 10-15 cells (**Figure S5A**), suggesting that NSD3-mediated deposition of H3K36me2 is not exclusive to megadomains. Given that megadomains and H3K27ac-BRD4-NUT peaks outside megadomains occupy only 1.10% and 0.60% of the genome in 10-15 and TC-797 cells, respectively, and no changes were observed in global levels of H3K36me2 upon NSD3 depletion (**Figures 2B and S4B**), the reduction of H3K36me2 within megadomains and at H3K27ac-BRD4-NUT peaks outside megadomains suggests that NSD3 selectively catalyzes H3K36me2 at these loci.

### NSD3 stabilizes BRD4-NUT condensates

Pharmacologic inhibition^62^ and degradation^34^ of BRD4-NUT cause complete loss of BRD4-NUT condensates. To dissect whether depletion of NSD3 affects BRD4-NUT condensate stability, we performed immunofluorescence staining, confocal imaging, and quantitative image analysis^63^ following NSD3 depletion in 10-15 and TC-797 cells (**Table S2 and S3**). NSD3 depletion removed nearly all NSD3 fluorescence signal within the nucleus, further confirming robust CRISPRi depletion at the single-cell level (**Figures 3A and S6A**). However, we observed the fluorescence signal from BRD4-NUT condensates to be less intense along with a greater number of BRD4-NUT condensates in NSD3-depleted cells compared to control cells (**Figure 3**). NSD3 depletion had only a minor effect on the total nuclear fluorescence intensity of BRD4-NUT. Mean nuclear BRD4-NUT fluorescence intensity in NSD3-depleted cells was 91.5% and 121.0% of the intensity in control cells for 10-15 and TC-797 cells, respectively (**Figure S6B**), consistent with immunoblotting that showed no change in BRD4-NUT protein levels following NSD3 depletion (**Figures 2B and S4B**). We therefore comprehensively and quantitatively analyzed BRD4-NUT nuclear condensates in NSD3-depleted and control cells. NSD3 depletion significantly reduced the average brightness (mean intensity) and overall brightness (total intensity) of BRD4-NUT within condensates in both cell types (**Figures 3B and S6C**). The mean BRD4-NUT condensate fluorescence intensity decreased 1.91-fold and 1.27-fold in 10-15 and TC-797 cells, respectively, following NSD3 depletion. The total BRD4-NUT fluorescence intensity in condensates decreased 2.02-fold and 1.28-fold in NSD3-depleted 10-15 and TC-797 cells, respectively. Given the mean nuclear BRD4-NUT fluorescence intensity in NSD3-depleted cells, the mean enrichment of BRD4-NUT in condensates decreased 1.75-fold and 1.54-fold in 10-15 and TC-797 cells, respectively. Because BRD4-NUT was less enriched in condensates while total nuclear BRD4-NUT fluorescence intensity did not substantially change following NSD3 depletion, this indicates that a larger fraction of BRD4-NUT resides outside condensates in NSD3-depleted cells. We also observed that the number of the BRD4-NUT condensates per nucleus significantly increased 1.33-fold and 1.31-fold in 10-15 and TC-797 cells, respectively, following NSD3 depletion (**Figure 3C**).

**Figure 3.**
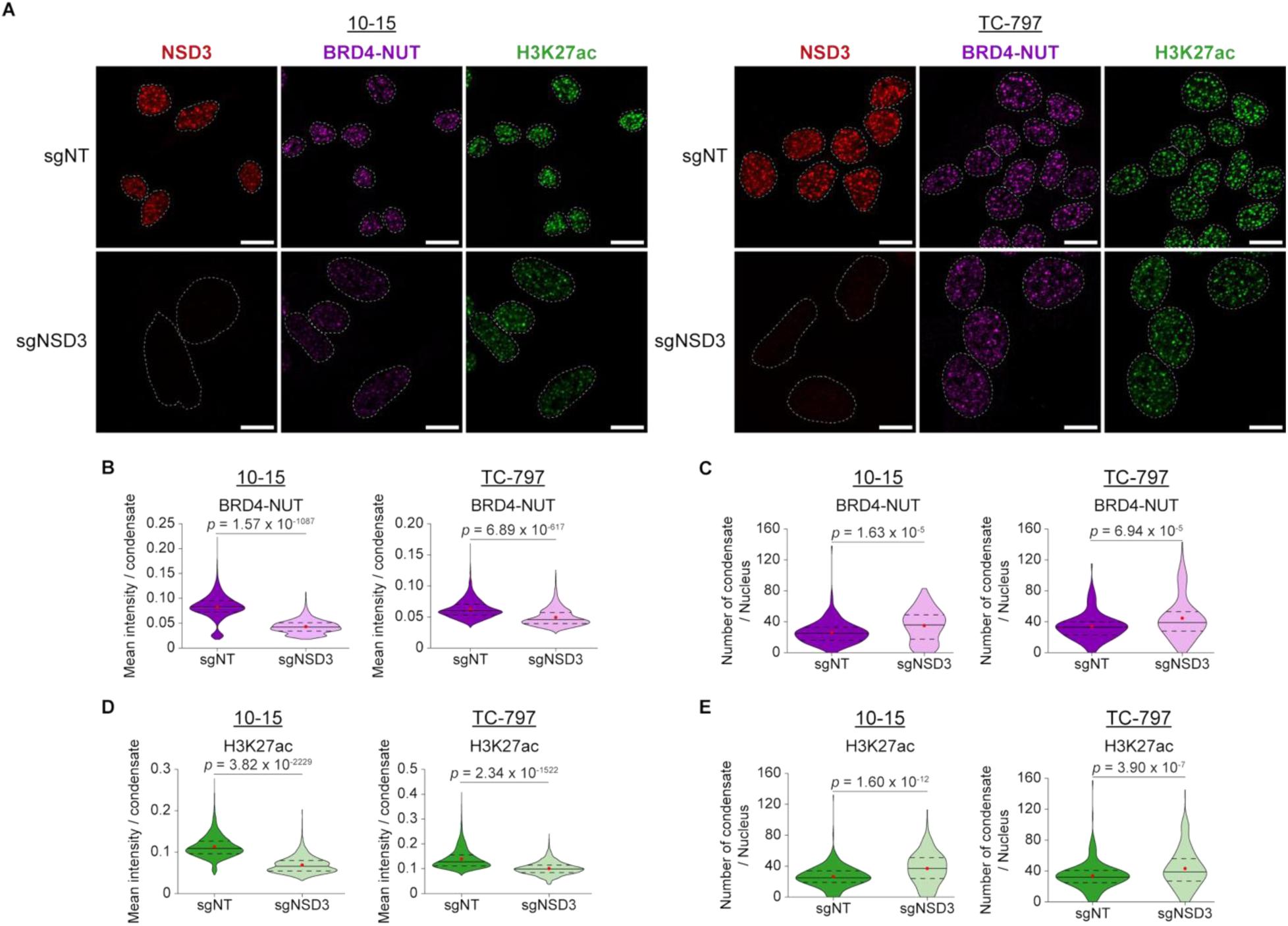
NSD3 stabilizes BRD4-NUT nuclear condensates. (**A**) Immunofluorescence for NSD3 (red), BRD4-NUT (purple), and histone H3K27ac (green) in control (sgNT) and NSD3-depleted (sgNSD3) 10-15 and TC-797 cells, respectively, show that the depletion of NSD3 destabilizes BRD4-NUT condensates containing histone H3K27ac. White dashed lines outline nuclei identified by DAPI staining. Scale bars, 10 µm. (**B**) Violin plots (solid lines indicate medians; red dots indicate means; dashed lines indicate 25th and 75th percentiles) indicate that depletion of NSD3 significantly reduces the mean fluorescence intensity of BRD4-NUT in BRD4-NUT condensates. 5,710 (sgNT, purple) and 4,318 (sgNSD3, pink) condensates were analyzed from 10-15 cell nuclei; 5,594 (sgNT, purple) and 7,511 (sgNSD3, pink) condensates were analyzed from TC-797 cell nuclei. p values determined using one-sided Mann-Whitney *U* tests. (**C**) Violin plots (solid lines indicate medians; red dots indicate means; dashed lines indicate 25th and 75th percentiles) indicate that depletion of NSD3 significantly increases the number of BRD4-NUT condensates per nucleus. 216 (sgNT, purple) and 123 (sgNSD3, pink) 10-15 cell nuclei were analyzed; 164 (sgNT, purple) and 168 (sgNSD3, pink) TC-797 cell nuclei were analyzed. p values determined using one-sided Mann-Whitney *U* tests. (**D**) Violin plots (solid lines indicate medians; red dots indicate means; dashed lines indicate 25th and 75th percentiles) indicate that depletion of NSD3 significantly reduces the mean fluorescence intensity of histone H3K27ac in condensates. 10,064 (sgNT, dark green) and 9,151 (sgNSD3, light green) condensates were analyzed from 10-15 cell nuclei; 10,691 (sgNT, dark green) and 13,408 (sgNSD3, light green) condensates were analyzed from TC-797 cell nuclei. p values determined using one-sided Mann-Whitney *U* tests. (**E**) Violin plots (solid lines indicate medians; red dots indicate means; dashed lines indicate 25th and 75th percentiles) indicate that depletion of NSD3 significantly increases the number of condensates per nucleus. 377 (sgNT, dark green) and 247 (sgNSD3, light green) 10-15 cell nuclei were analyzed; 314 (sgNT, dark green) and 309 (sgNSD3, light green) TC-797 cell nuclei were analyzed. p values determined using one-sided Mann-Whitney *U* tests.

We quantified the histone H3K27ac fluorescence signal as an additional means to analyze BRD4-NUT condensates in NSD3-depleted cells. NSD3 depletion resulted in a slight increase of 1.14-fold and 1.08-fold of total nuclear histone H3K27ac signal intensity that was statistically significant for 10-15 cells but insignificant for TC-797 cells, respectively (**Figure S6D**). Consistent with our observations for BRD4-NUT, NSD3 depletion significantly reduced the average brightness and overall brightness of H3K27ac in condensates in both cells (**Figures 3D and S6E**). The mean H3K27ac fluorescence intensity in condensates decreased 1.63-fold and 1.37-fold in NSD3-depleted 10-15 and TC-797 cells, respectively. The total H3K27ac fluorescence intensity decreased 1.69-fold and 1.35-fold following NSD3 depletion from 10-15 cells and TC-797 cells, respectively. Like the number of BRD4-NUT condensates, the number of H3K27ac foci significantly increased 1.39-fold and 1.27-fold in 10-15 and TC-797 cells, respectively, following NSD3 depletion (**Figure 3E**). Taken together, loss of NSD3 destabilizes BRD4-NUT nuclear condensates.

NSD3 depletion induces morphological changes consistent with cell differentiation (**Figure S3B**). To rule out the possibility that any changes in BRD4-NUT condensates were due to alterations in cell morphology, we analyzed nuclear volume following NSD3 depletion. Although NSD3 depletion led to a significant increase in nuclear volume (1.50-fold relative to control) in 10-15 cells, the nuclear volume of TC-797 cells remained unchanged (1.02-fold relative to control) (**Figure S6F**), suggesting that any changes in nuclear volume are cell type-specific and unlikely to universally contribute to changes in BRD4-NUT condensates. Collectively, our findings demonstrate that NSD3 maintains the stability of BRD4-NUT nuclear condensates.

### NSD3 maintains chromatin 3D contacts between megadomains

Previously, when we induced degradation of BRD4-NUT we observed that loss of BRD4-NUT condensates coincided with loss of intrachromosomal (*cis*) and interchromosomal (*trans*) chromatin interactions between megadomains within a nuclear subcompartment^34^, a structure that is referred to as microcompartments in healthy cells^1,25^. Because depletion of NSD3 destabilizes BRD4-NUT condensates, we asked whether loss of NSD3 similarly weakens chromatin 3D contacts between megadomains. We therefore performed *in situ* high-throughput chromosome conformation capture (Hi-C)^64^ to identify 3,768,750,831 and 2,038,321,587 chromosomal contacts (read pairs that remain after removal of duplicates, poor alignment to the reference genome, and unligated fragments) in NSD3-depleted and control 10-15 cells, respectively; and 2,454,474,310 and 2,038,261,595 chromosomal contacts in NSD3-depleted and control TC-797 cells, respectively (**Table S4**). We first looked for any global changes in chromosome architecture by analyzing the random polymer conformations for all chromosomes by plotting the average intrachromosomal contact probability (*P*) between loci as a function of their linear genomic separation (*s*) to create *P*(*s*) curves^24^. The *P*(*s*) curves of NSD3-depleted and control cells were identical for both 10-15 cells and TC-797 cells (**Figure S7A**), indicating that depletion of NSD3 does not alter global chromosome conformation. Moreover, to exclude any technical biases during the Hi-C procedure, we spiked in *Drosophila melanogaster* cells to human cells prior to cross-linking (Methods). The *P*(*s*) curves for read pairs that aligned to the *Drosophila melanogaster* genome were identical for control and NSD3-depleted samples indicating an absence of technical artifacts in both 10-15 and TC-797 cells (**Figure S7B**).

Unlike the complete loss of BRD4-NUT condensates following degradation of BRD4-NUT^34^, depletion of NSD3 destabilizes condensates. This suggests that, rather than acting as the primary driver of BRD4-NUT-driven chromatin contacts, NSD3 may function as a cofactor that modulates megadomain-megadomain interactions. To elucidate a role for NSD3 in BRD4-NUT-driven compartmentalization, we assessed whether loss of NSD3 altered megadomain-megadomain contact frequency. Visual inspection of the Hi-C contact maps revealed that depletion of NSD3 reduced the contact frequency between megadomains both in *cis* (**Figures 4A and S8A**) and in *trans* (**Figures 4B and S8B**).

**Figure 4.**
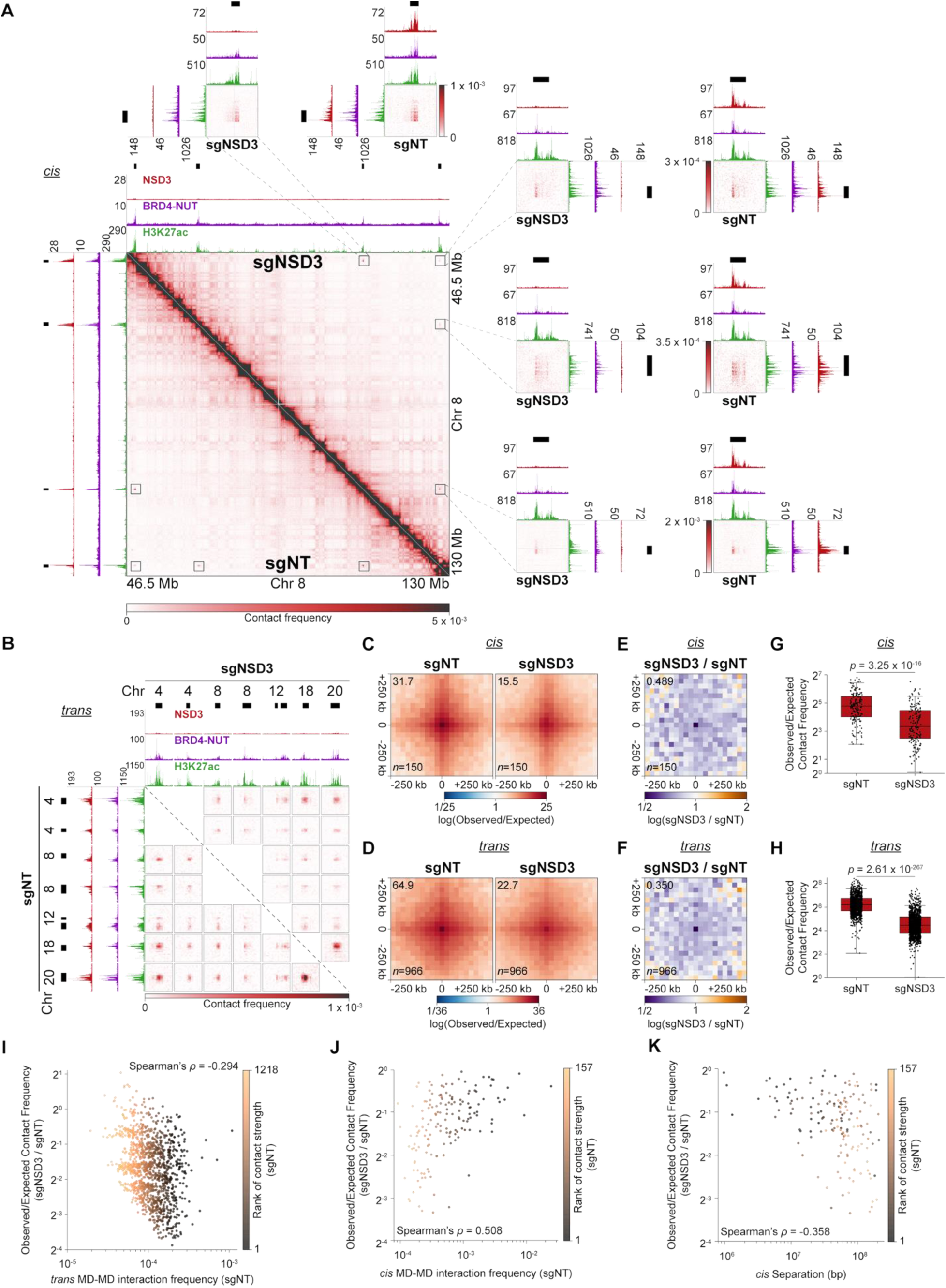
NSD3 stabilizes chromatin interactions between megadomains. (**A**) Intrachromosomal (*cis*) Hi-C contact maps at 100 kb resolution (left bottom) from control (sgNT; below diagonal) and NSD3-depleted 10-15 cells (sgNSD3; above diagonal) show that depletion of NSD3 reduces *cis* megadomain-megadomain (MD-MD) contacts. Boxed regions displaying individual MD-MD contacts are enlarged to the right and above the contact map at 10 kb resolution. CUT&RUN profiles for NSD3, BRD4-NUT, and histone H3K27ac are aligned with the contact maps (sgNT: vertically; sgNSD3: horizontally for the 100 kb resolution map). Black bars indicate megadomains. (**B**) Interchromosomal (*trans*) Hi-C contact maps at 100 kb resolution from control (sgNT; below diagonal) and NSD3-depleted (sgNSD3; above diagonal) 10-15 cells show that the depletion of NSD3 reduces *trans* MD-MD contacts. CUT&RUN profiles for NSD3, BRD4-NUT, and histone H3K27ac are aligned with the contact maps (sgNT: vertically; sgNSD3: horizontally for the 100 kb resolution map). Black bars indicate megadomains. (**C**) Genome-wide pileups of mean observed/expected *cis* Hi-C contact frequencies at MD-MD contacts for control (sgNT) and NSD3-depleted (sgNSD3) 10-15 cells. Values of the central pixel are given at top left. Windows extending beyond chromosome ends and contacts with zero or undefined observed/expected values in any dataset were excluded from pileups. (**D**) Genome-wide pileups of mean observed/expected *trans* Hi-C contact frequencies at MD-MD contacts for control (sgNT) and NSD3-depleted 10-15 cells (sgNSD3). Values of the central pixel are given at top left. Windows extending beyond chromosome ends and contacts with zero or undefined observed/expected values in any dataset were excluded from pileups. (**E**) Genome-wide pileups of the fold-change in mean observed/expected *cis* Hi-C contact frequencies at MD-MD contacts for NSD3-depleted (sgNSD3) versus control (sgNT) 10-15 cells. The value of the central pixel is given at top left. Windows extending beyond chromosome ends and contacts with zero or undefined observed/expected values in any dataset were excluded from pileups. (**F**) Genome-wide pileups of the fold-change in mean observed/expected *trans* Hi-C contact frequencies at MD-MD contacts for NSD3-depleted (sgNSD3) versus control (sgNT) 10-15 cells. The value of the central pixel is given at top left. Windows extending beyond chromosome ends and contacts with zero or undefined observed/expected values in any dataset were excluded from pileups. (**G**) Box plots (horizontal lines show medians; boxes indicate interquartile range; whiskers extend 1.5 times the interquartile range) of mean observed/expected *cis* Hi-C contact frequencies for MD-MD contacts for control (sgNT) and NSD3-depleted (sgNSD3) 10-15 cells show a significant reduction of *cis* MD-MD contacts upon NSD3 depletion. Contacts with zero or undefined observed/expected values in any dataset were removed for a total of 157 *cis* MD-MD contacts. *p* value determined using a one-sided Mann-Whitney *U* test. (**H**) Box plots (horizontal lines show medians; boxes indicate interquartile range; whiskers extend 1.5 times the interquartile range) of mean observed/expected *trans* Hi-C contact frequencies for MD-MD contacts for control (sgNT) and NSD3-depleted (sgNSD3) 10-15 cells show a significant reduction of *trans* MD-MD contacts upon NSD3 depletion. Contacts with zero or undefined observed/expected values in any dataset were removed for a total of 1,218 *trans* MD-MD contacts. *p* values determined using a one-sided Mann-Whitney *U* test. (**I**) For each *trans* MD-MD contact (*n* = 1,218), the fold-change in mean observed/expected Hi-C contact frequency versus the observed MD-MD contact frequency in control (sgNT) 10-15 cells. The color of data points reflects the rank of the observed *trans* MD-MD contact frequency from least to greatest in control (sgNT) 10-15 cells. (**J**) For each *cis* MD-MD contact (*n* = 157), the fold-change in mean observed/expected Hi-C contact frequency versus the observed MD-MD contact frequency in control (sgNT) 10-15 cells. The color of data points reflects the rank of the observed *cis* MD-MD contact frequency from least to greatest in control (sgNT) 10-15 cells. (**K**) For each *cis* MD-MD contact (*n* = 157), the fold-change in mean observed/expected Hi-C contact frequency versus the genomic distance separating the interacting megadomains. The color of data points reflects the rank of the observed *cis* MD-MD contact frequency from least to greatest in control (sgNT) 10-15 cells.

We examined megadomain-megadomain contacts genome-wide. We identified megadomain-megadomain interactions using our previously defined method^34^, and refined this approach from 100 kb to 25 kb resolution by identifying the peak pixel for each interaction (**Figure S9**). We compared megadomain-megadomain peak pixel interaction frequency in NSD3-depleted versus control cells using pileups. To create pileups, we extracted each peak pixel plus flanking DNA from the Hi-C contact map and then averaged the Hi-C contact frequency normalized for expected random polymer conformations of chromosomes (for *cis* interactions) or normalized for pairwise interchromosomal average interaction frequency (for *trans* interactions) for all megadomain-megadomain interactions across all chromosomes in NSD3-depleted and control cells. Pileup analysis demonstrated a reduction in megadomain-megadomain peak pixel interaction strength in both *cis* (**Figures 4C and S8C**) and *trans* (**Figures 4D and S8D**) following NSD3 depletion. *Cis* contact frequency between megadomains in NSD3-depleted cells was 48.9% and 48.7% of the contact frequency in control cells for 10-15 and TC-797 cells, respectively (**Figures 4E and S8E**). *Trans* contact frequency between megadomains in NSD3-depleted cells was 35.0% and 34.4% of the contact frequency in control cells for 10-15 and TC-797 cells, respectively (**Figures 4F and S8F**). In addition to pileups, which display the genome-wide average contact frequency, we also plotted the normalized Hi-C contact frequency for each megadomain-megadomain interaction in control and NSD3-depleted cells, which confirmed a significant reduction in peak pixel contact frequency upon depletion of NSD3 in both *cis* (**Figures 4G and S8G**) and *trans* (**Figures 4H and S8H**) in both cell lines. Our Hi-C analysis indicates a role for NSD3 in stabilizing megadomain-megadomain interactions.

Pileup analysis indicated that *trans* megadomain-megadomain interactions were more sensitive to NSD3 depletion than *cis* megadomain-megadomain interactions (**Figures 4E and 4F**). We therefore asked why *trans* interactions may be more sensitive than *cis* interactions. Because depletion of NSD3 destabilizes, but does not totally abrogate, BRD4-NUT condensates (**Figure 3**), we considered that NSD3 may have a more prominent role in stabilizing stronger rather than weaker megadomain-megadomain interactions. To assess whether interactions with higher contact frequencies are more affected by NSD3 depletion, we examined the relationship between the megadomain-megadomain interaction peak pixel contact frequency observed in control cells and the fold-change in expected-normalized peak pixel contact frequency in NSD3-depleted versus control cells for each megadomain-megadomain interaction. For *trans* megadomain-megadomain interactions, we observed a negative correlation between the contact frequency in control cells and the fold-change in contact frequency following NSD3 depletion in both 10-15 cells (**Figure 4I**, Spearman’s ρ = -0.294, *p*-value = 1.10 x 10^-25^) and TC-797 cells (**Figure S8I**, Spearman’s ρ = -0.353, *p*-value = 8.12 x 10^-21^). We used pileups to substantiate this trend. We assigned *trans* megadomain-megadomain interactions to quintiles based upon the peak pixel contact frequency in control cells and then calculated pileups in each quintile for control and NSD3-depleted cells. Consistent with a negative correlation between the contact frequency in control cells and the fold-change in contact frequency following NSD3 depletion, pileups showed the greatest reduction in contact strength following NSD3 depletion for the strongest interactions (**Figure S10A**), indicating that NSD3 preferentially stabilizes stronger *trans* interactions between megadomains.

In contrast, *cis* megadomain-megadomain interactions exhibited the opposite trend. We observed a positive correlation between contact strength in control samples and the fold-change in contact frequency following NSD3 depletion in both 10-15 cells (**Figure 4J**, Spearman’s ρ = 0.508, *p*-value = 1.12 x 10^-11^) and TC-797 cells (**Figure S8J**, Spearman’s ρ = 0.232, *p*-value = 4.09 x 10^-2^). Like *trans* contacts, we assigned *cis* interactions to quintiles based upon contact frequency in control cells and then calculated pileups in each quintile for control and NSD3-depleted cells. Consistent with the interaction-by-interaction analysis, contacts with weaker *cis* contact strength were more strongly affected after depletion of NSD3 (**Figure S10B**), indicating that weaker intrachromosomal interactions are more susceptible to NSD3 loss. *Cis* contact frequency is influenced by genomic distance, with proximal loci exhibiting higher contact frequencies than distal loci^24^. This pattern is evident in Hi-C contact maps, where short-range interactions closer to the diagonal exhibit stronger contact frequency than interactions farther from the diagonal. To determine whether the positive correlation between contact strength in control cells and the fold-change in contact frequency following NSD3 depletion is due to the distance dependence of *cis* contacts, we examined the relationship between the genomic separation of interacting megadomains and the fold-change in peak pixel expected-normalized contact frequency after depletion of NSD3. We observed a negative correlation between genomic separation and the fold-change in interaction strength following NSD3 depletion in 10-15 cells (**Figure 4K**, Spearman’s ρ = -0.358, *p*-value = 4.22 x 10^-6^) and TC-797 cells (**Figure S8K**, Spearman’s ρ = -0.349, *p*-value = 1.75 x 10^-3^). To investigate this trend further, we assigned interactions into quintiles based on genomic separation and then calculated pileups in each quintile for control and NSD3-depleted cells. Pileups showed that interactions spanning longer genomic distances were the most affected by NSD3 depletion (**Figure S10C**). These findings suggest that NSD3 more strongly contributes to distal *cis* megadomain-megadomain interactions than to more proximal interactions.

### NSD3 supports BRD4-NUT-driven oncogene activation

Genes located in megadomains play a crucial role in NUT carcinoma tumorigenesis^53,56^. We previously showed that megadomain-megadomain interactions promote oncogene expression^34^. Because NSD3 stabilizes megadomain-megadomain interactions, we next investigated a role for NSD3 in megadomain gene expression. We therefore performed RNA-seq on NSD3-depleted and control cells. Of the 241 differentially expressed genes (≥1.5-fold change in expression; Benjamini-Hochberg adjusted *p*-value < 0.05) following NSD3 depletion, downregulated megadomain genes (25 of 68 all downregulated genes) were enriched relative to upregulated megadomain genes (0 of 173 all upregulated genes) in 10-15 cells (**Figure S11A**).

Of the 369 differentially expressed genes following NSD3 depletion in TC-797 cells (**Figure S11A**), downregulated megadomain genes (10 of 67 all downregulated genes) were 9.01-fold enriched relative to upregulated megadomain genes (5 of 302 all upregulated genes). NSD3 depletion affects a substantially smaller subset of genes compared to the 7,610 differentially expressed genes following induced degradation of BRD4-NUT in TC-797 cells in our previous study^34^. This suggests that NSD3 likely fine-tunes gene activation. Additionally, Gene Set Enrichment Analysis (GSEA) demonstrated that megadomain genes were significantly enriched as being downregulated in both cell lines (**Figure S11B**), further supporting the role of NSD3 in sustaining transcriptional activity due to BRD4-NUT-driven nuclear compartmentalization.

*MYC* is a direct target of BRD4-NUT^56^; the *MYC* promoter is located within a megadomain in 10-15 cells and is downstream of a megadomain in TC-797 cells^53^. NSD3 depletion reduced *MYC* expression by 34.1% (*p*-adjust = 1.86 x 10^-9^) and 29.3% (*p*-adjust = 7 x 10^-4^) in 10-15 and TC-797 cells, respectively (**Table S5**). GSEA revealed that MYC target genes were significantly enriched as being downregulated following NSD3 depletion (**Figures S12A and S12B**), indicating that NSD3 plays a role in supporting MYC-driven transcriptional programs in NUT carcinoma.

Consistent with NUT carcinoma cellular differentiation upon NSD3 depletion (**Figure S3**), upregulated genes following NSD3 loss were significantly enriched in multiple differentiation and developmental pathways. In 10-15 cells, these include mesenchymal cell differentiation, skin development, and kidney development, while in TC-797 cells they include keratinocyte differentiation, keratinization, and epidermis development (**Figure S12C**). Together, these findings suggest that NSD3 maintains NUT carcinoma cells in a poorly differentiated state by functioning as a cofactor with BRD4-NUT to drive megadomain-megadomain interactions and condensate formation that influence gene expression.

### The short, catalytically inactive isoform of NSD3 promotes chromatin contacts

As a further test of a role for NSD3 in mediating chromosome folding, we asked whether gain of NSD3 in cells would promote chromatin contacts. NSD3 is amplified and oncogenic in many cancers^51,52,65–68^. Expression of the catalytically inactive short isoform of NSD3 (NSD3short) transforms MCF10A breast epithelial cells^65^ and induces non-tumorigenic MCF10A cells to form breast tumors at a comparable rate and size of tumors driven by the potent oncogene *HRAS*^69^. We therefore expressed NSD3short fused to a HiBiT epitope tag in MCF10A cells under the control of a doxycycline-inducible promoter (**Figure 5A**). Uninduced MCF10A cells transitioned from an epithelial morphology to a more mesenchymal phenotype upon induced expression of NSD3short (**Figure S13A**), consistent with a previous report^69^. RNA-seq analysis indicated that genes that promote an epithelial to mesenchymal transition were upregulated in cells induced to express NSD3short (**Figures S13B and S13C**), consistent with changes in cell morphology. MCF10A cells expressing NSD3short grew at a slower rate than uninduced cells (**Figure S13D**). Importantly, MCF10A cells do not express BRD4-NUT. Therefore, our NSD3short gain-of-function system in fusion-naïve cells complements our loss-of-function experiments in cells that express BRD4-NUT.

**Figure 5.**
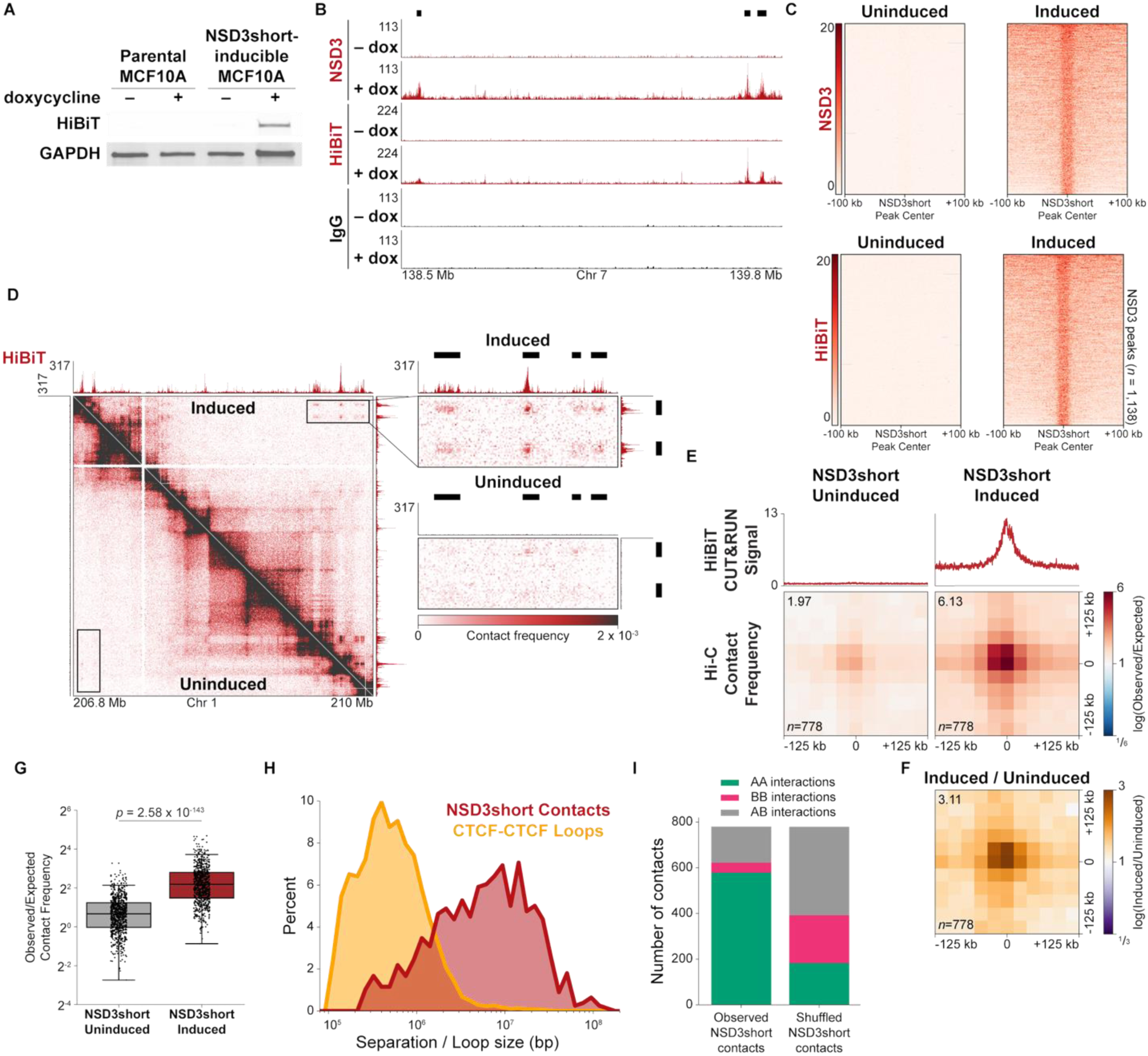
NSD3short promotes chromatin contacts separated by multiple megabases. (**A**) Immunoblots indicate induced expression of HiBiT-tagged NSD3short in MCF10A cells after treatment with 30 ng/mL doxycycline. Water and parental MCF10A cells were used as controls. GAPDH was used as an internal control. (**B**) CUT&RUN profiles for NSD3 (red), HiBiT epitope tag (red), and IgG (black) show NSD3short peaks (black bars) in MCF10A cells induced to express NSD3short versus uninduced MCF10A cells. (**C**) Heatmaps of the CUT&RUN signal for NSD3 and HiBiT epitope tag within each NSD3short peak in uninduced MCF10A cells and cells induced to express NSD3short. NSD3short peak centers were aligned and 100 kb of DNA flanking each NSD3short peak is shown. IgG signal has been subtracted from each heatmap. (**D**) Hi-C contact maps at 5 kb resolution from uninduced MCF10A cells (below diagonal) and cells induced to express NSD3short (above diagonal) show NSD3short-dependent chromatin contacts. Boxed regions displaying interactions separated by >2 Mb are enlarged to the right of the contact map. CUT&RUN profiles for the HiBiT epitope tag are aligned with the contact maps. Black bars indicate NSD3short peaks. (**E**) Genome-wide pileups of mean observed/expected Hi-C contact frequencies at NSD3short-dependent chromatin contacts in uninduced MCF10A cells and cells induced to express NSD3short show that NSD3short promotes chromatin contacts. The mean HiBiT CUT&RUN signal at NSD3short contact sites is shown above the pileups. Values of the central pixel are given at top left. (**F**) Genome-wide pileups of the fold-change in mean observed/expected Hi-C contact frequencies at NSD3short-dependent chromatin contacts in MCF10A cells induced to express NSD3short versus uninduced cells. The value of the central pixel is given at top left. (**G**) Box plots (horizontal lines indicate medians; boxes indicate interquartile range; whiskers extend 1.5 times the interquartile range) of observed/expected Hi-C contact frequencies for NSD3short-dependent chromatin contacts in uninduced MCF10A cells and cells induced to express NSD3short. Data points represent individual chromatin contacts. Contacts with zero or undefined observed/expected values in any dataset were removed for a total of 771 analyzed contacts. *p* value determined using a one-sided Mann-Whitney *U* test. (**H**) Percent of NSD3short-dependent chromatin contacts (red) separated by the distance given on the x-axis and percent of convergent CTCF-CTCF loops (orange) of a given size. The separation between NSD3short contact sites and loop sizes is logarithmically binned. (**I**) Number of NSD3short-dependent chromatin contacts overlapping with AA, BB, or AB Hi-C compartment interactions in uninduced MCF10A cells compared to a random shuffled control set of contacts shows that NSD3short promotes chromatin contacts within the A compartment.

Next, we performed CUT&RUN for both NSD3 and for the HiBiT epitope tag in uninduced MCF10A cells and cells induced to express NSD3short. NSD3short localized on chromatin as evident by peaks of NSD3 and HiBiT CUT&RUN signal (**Figure 5B**). We used the signal from the HiBiT tag to specifically identify highly enriched peaks relative to IgG control (**Figure 5C**; 1,138 peaks). We also performed *in situ* Hi-C in uninduced MCF10A cells and cells induced to express NSD3short identifying 3,143,125,703 and 2,905,877,244 chromatin contacts, respectively (**Table S4**). Global chromosome conformation was comparable between uninduced MCF10A cells and cells induced to express NSD3short (**Figures S14A and S14B**). Visual inspection of the Hi-C contact maps revealed focal contacts between NSD3short peaks (**Figure 5D**). In a manner similar to identifying megadomain-megadomain contacts, we enumerated all possible pairwise interactions between NSD3short peaks and identified 778 contacts between NSD3short peaks enriched relative to local background that were specific to cells expressing NSD3short. We substantiated contacts between NSD3short peaks across all chromosomes using pileups. To create pileups, we extracted each contact between NSD3short peaks plus flanking DNA from the Hi-C contact map, re-scaled the extracted regions to the same dimensions, and then averaged the Hi-C contact frequency normalized for expected random polymer conformations of chromosomes, pixel-by-pixel, across all such regions in uninduced MCF10A cells and cells induced to express NSD3short. We also identified the NSD3short peaks making contacts, so-called NSD3short contact sites, averaged the HiBiT CUT&RUN signal from both experimental conditions at NSD3short contact sites, and aligned the mean HiBiT CUT&RUN signal with the pileups. Pileup analysis demonstrated that NSD3short promotes chromatin contacts (**Figure 5E**). In addition to normalizing for expected random polymer conformations of chromosomes, we also used randomly shifted regions as a control, which also demonstrated that NSD3short promotes chromatin contacts (**Figure S14C**). Contact frequency between NSD3short contact sites was 3.11-fold greater in cells expressing NSD3short compared to uninduced control cells (**Figures 5F and S14D**). We also plotted the Hi-C contact frequency normalized for the expected random polymer conformations of chromosomes for individual NSD3short contacts in uninduced MCF10A cells and cells induced to express NSD3short (**Figure 5G**), which confirmed that NSD3short promotes chromatin contacts and provided statistical significance at the level of individual NSD3short contacts.

Cohesin-mediated loop extrusion stalled by CTCF is apparent in Hi-C contact maps as focal contact enrichment, referred to as DNA loops, with convergently oriented CTCF motifs at the loop anchors^64^. We annotated 15,920 DNA loops with convergent CTCF motifs in cells expressing NSD3short and compared the distribution of distances separating NSD3short contact sites with the distribution of CTCF-CTCF loop sizes. This revealed two distributions with centers that did not overlap: the distance between NSD3short contact sites was centered around 10 Mb, whereas CTCF-CTCF loop sizes were centered around 400 kb (**Figure 5H**).

At the scale of multiple megabase pairs, Hi-C contact maps have revealed that groups of loci preferentially associate into a euchromatic A compartment and a heterochromatic B compartment^24^. We identified A and B compartments in uninduced MCF10A cells and asked how NSD3short contacts overlapped with chromatin interactions within and between compartments. 74.4% of NSD3short contacts coincided with AA compartment interactions in uninduced MCF10A cells (**Figure 5I**), significantly greater than that expected by chance (**Figure S14E**), indicating that NSD3short contacts arise from the A compartment.

### NSD3short’s ability to promote chromatin contacts depends on its PWWP domain

We more deeply probed how NSD3short mediates chromosome folding. NSD3short contains a single known, structurally ordered chromatin reader module: a Pro-Trp-Trp-Pro (PWWP) domain^51,52^. NSD3short’s PWWP domain binds to histone H3 when it is methylated at lysine 36^70,71^. A tryptophan-to-alanine mutation at amino acid residue 284 (W284A) inactivates the PWWP domain^45^ and slows NSD3 recruitment to chromatin in living cells^72^ (**Figure 6A**). We therefore expressed NSD3short^W284A^ at comparable levels to wild-type NSD3short in MCF10A cells (**Figure 6B and C**). MCF10A cells expressing NSD3short^W284A^ have an intermediate morphology between epithelial, uninduced MCF10A cells and mesenchymal-like cells that express wild-type NSD3short (**Figure S15A**). RNA-seq analysis supported that cells induced to express NSD3short^W284A^ upregulated genes that promoted an epithelial to mesenchymal transition (**Figures S15B and S15C**). MCF10A cells expressing NSD3short^W284A^ grew slower than uninduced MCF10A cells, but faster than cells expressing wild-type NSD3short (**Figure S15D**).

**Figure 6.**
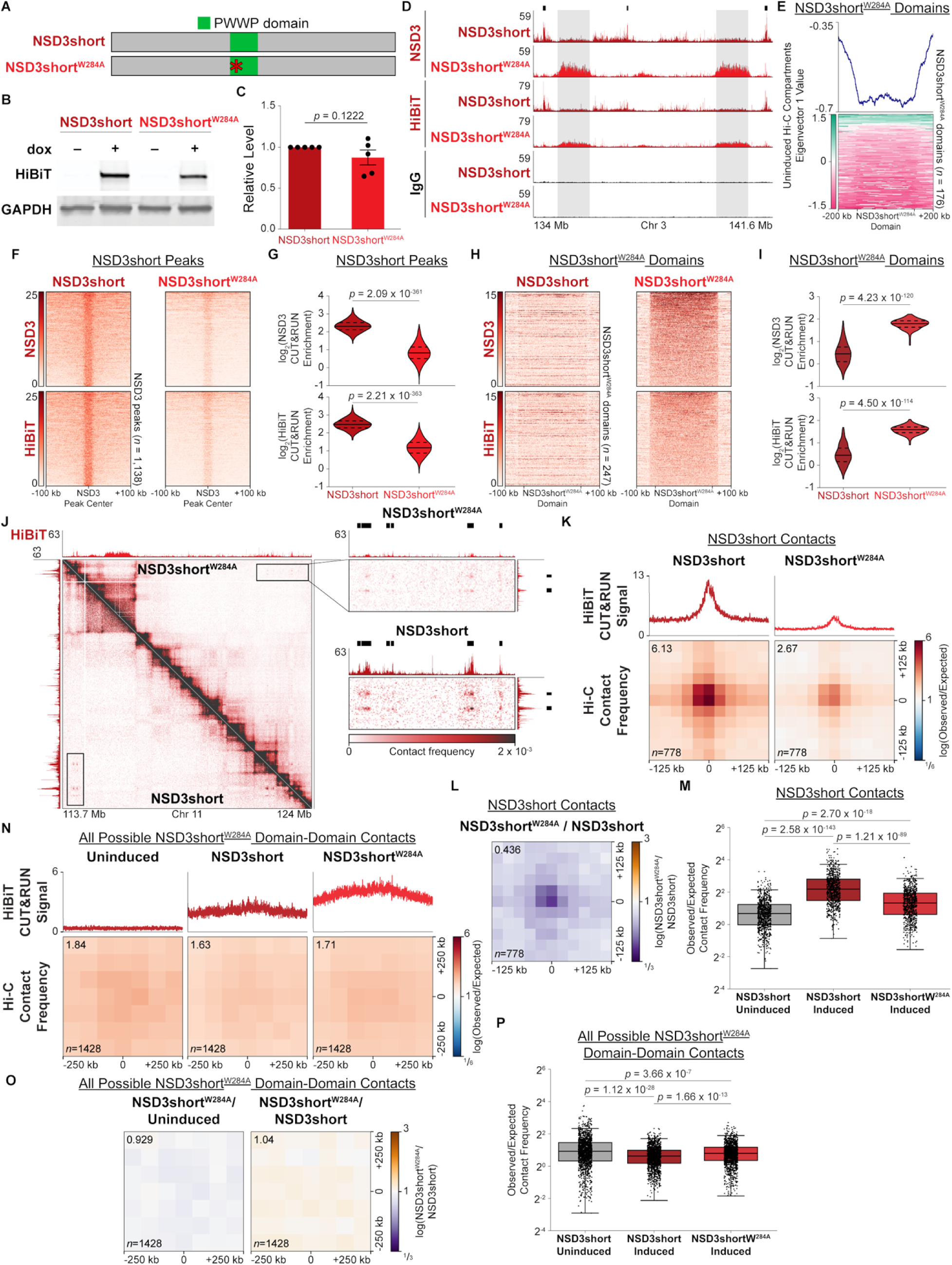
NSD3short requires the PWWP domain to promote chromatin contacts. (**A**) Domain organization of NSD3short. A Trp284Ala mutation inactivates the PWWP domain of NSD3short. (**B**) Immunoblots indicate comparable expression levels of HiBiT-tagged NSD3short and NSD3short^W284A^. MCF10A cells were treated with 30 ng/mL or 25 ng/mL doxycycline to induce expression of NSD3short or NSD3short^W284A^, respectively. GAPDH was used as an internal control. (**C**) Quantification of immunoblots related to (B). Bar and error bars represent the mean and standard error of the mean, respectively, of five biological replicates. p value determined using paired t-test. (**D**) CUT&RUN profiles for NSD3, HiBiT epitope tag, and IgG from MCF10A cells expressing NSD3short (dark red) and NSD3short^W284A^ (light red). NSD3short^W284A^ chromatin occupancy is reduced within NSD3short peaks (black bars) and redistributes to broad domains (gray shading) with low NSD3short wild-type occupancy. (**E**) Profile (top) and heatmap (bottom) of the Hi-C compartment eigenvector 1 values from uninduced MCF10A cells within NSD3short^W284A^ domains show that NSD3short^W284A^ domains form in regions of negative eigenvector 1 values corresponding to B compartment intervals in uninduced cells. Profile represents the mean of the Hi-C compartment eigenvector 1 value at NSD3short^W284A^ domains. NSD3short^W284A^ domains were normalized to the same length and 200 kb of flanking DNA is shown next to each normalized NSD3short^W284A^ domain. (**F**) Heatmaps of the CUT&RUN signal for NSD3 and HiBiT epitope tag centered at NSD3short peaks in MCF10A cells expressing NSD3short or NSD3short^W284A^ show reduced genome-wide occupancy of NSD3short^W284A^ in NSD3short peaks. NSD3short peak centers were aligned and 100 kb of DNA flanking each NSD3short peak is shown. IgG signal has been subtracted from each heatmap. (**G**) Violin plots (solid lines indicate medians; dashed lines indicate 25th and 75th percentiles) indicate that NSD3 and HiBiT epitope tag CUT&RUN enrichment relative to IgG within NSD3short peaks is decreased for NSD3short^W284A^ compared to NSD3short. *p* values determined using one-sided Mann-Whitney *U* tests. (**H**) Heatmaps of the CUT&RUN signal for NSD3 and HiBiT epitope tag within each NSD3short^W284A^ domain in MCF10A cells expressing NSD3short or NSD3short^W284A^ show increased genome-wide occupancy of NSD3short^W284A^ in NSD3short^W284A^ domains. NSD3short^W284A^ domains were normalized to the same length and 100 kb of flanking DNA is shown next to each normalized NSD3short^W284A^ domain. IgG signal has been subtracted from each heatmap. (**I**) Violin plots (solid lines indicate medians; dashed lines indicate 25th and 75th percentiles) indicate that NSD3 and HiBiT epitope tag CUT&RUN enrichment relative to IgG within NSD3short^W284A^ domains is increased for NSD3short^W284A^ compared to NSD3short. *p* values determined using one-sided Mann-Whitney *U* tests. (**J**) Hi-C contact maps at 10 kb resolution from MCF10A cells expressing NSD3short (below diagonal) and cells expressing NSD3short^W284A^ (above diagonal) show that NSD3short^W284A^ has a reduced capacity to promote chromatin contacts. Boxed regions displaying interactions separated by >8 Mb are enlarged to the right of the contact map. CUT&RUN profiles for the HiBiT epitope tag are aligned with the contact maps. Black bars indicate NSD3short peaks. (**K**) Genome-wide pileups of mean observed/expected Hi-C contact frequencies at NSD3short-dependent chromatin contacts in MCF10A cells expressing NSD3short and cells expressing NSD3short^W284A^. The mean HiBiT CUT&RUN signal at NSD3short contact sites is shown above the pileups. Values of the central pixel are given at top left. (**L**) Genome-wide pileups of the fold-change in mean observed/expected Hi-C contact frequencies at NSD3short-dependent chromatin interactions in MCF10A cells expressing NSD3short and cells expressing NSD3short^W284A^. The value of the central pixel is given at top left. (**M**) Box plots (horizontal lines indicate medians; boxes indicate interquartile range; whiskers extend 1.5 times the interquartile range) of observed/expected Hi-C contact frequencies for NSD3short-dependent chromatin contacts in uninduced MCF10A cells, cells expressing NSD3short, and cells expressing NSD3short^W284A^. Data points represent individual chromatin contacts. Contacts with zero or undefined observed/expected values in any dataset were removed for a total of 771 analyzed contacts. p values determined using one-sided Mann-Whitney *U* tests. Data for uninduced MCF10A cells and cells expressing NSD3short are reproduced from Figure 5G. (**N**) Genome-wide pileups of mean observed/expected Hi-C contact frequencies at all possible NSD3short^W284A^ domain-domain contacts in uninduced MCF10A cells, cells expressing NSD3short, and cells expressing NSD3short^W284A^ show that NSD3short^W284A^ domains do not interact. The mean HiBiT CUT&RUN signal at NSD3short^W284A^ domains is shown above the pileups. Values of the central pixel are given at top left. (**O**) Genome-wide pileups of the fold-change in mean observed/expected Hi-C contact frequencies at all possible NSD3short^W284A^ domain-domain contacts in uninduced MCF10A cells, cells expressing NSD3short, and cells expressing NSD3short^W284A^. The value of the central pixel is given at top left. (**P**) Box plots (horizontal lines indicate medians; boxes indicate interquartile range; whiskers extend 1.5 times the interquartile range) of observed/expected Hi-C contact frequencies for all possible NSD3short^W284A^ domain-domain contacts in uninduced MCF10A cells, cells expressing NSD3short, and cells expressing NSD3short^W284A^. Data points represent individual chromatin contacts. Contacts with zero or undefined observed/expected values in any dataset were removed for a total of 1,443 analyzed contacts. *p* values determined using two-sided Mann-Whitney *U* tests.

We performed CUT&RUN for both NSD3 and the HiBiT tag in cells induced to express NSD3short^W284A^. NSD3short^W284A^ did not form well-defined peaks at the same locations as wild-type NSD3short (**Figure 6D**) consistent with the W284A mutation inactivating the PWWP domain and thus NSD3short’s ability to properly localize on chromatin. Strikingly, NSD3short^W284A^ redistributed on chromatin and formed broad domains that spanned multiple megabases (**Figure 6D**). Due to their comparable sizes, we annotated NSD3short^W284A^ domains in a manner analogous to annotating megadomains (Methods). To determine where NSD3short^W284A^ was redistributing to, we examined the location of NSD3short^W284A^ domains with respect to A and B compartments in uninduced MCF10A cells. The first eigenvector (EV) of Hi-C contact maps is used to classify loci into A and B compartments, with positive EV1 values assigned to the A compartment and negative EV1 values assigned to the B compartment. EV1 values within NSD3short^W284A^ domains were predominantly negative (**Figure 6E**), and 88.6% of NSD3short domains overlapped B compartment intervals in uninduced MCF10A cells (**Figure S16A**), greater than that expected by chance (**Figure S16B**). This indicates that NSD3short^W284A^ domains were arising from the B compartment in uninduced cells. The redistribution of NSD3short on chromatin resulted in a decreased occupancy of NSD3short^W284A^ at NSD3short peaks (**Figures 6F and G**) and an increased occupancy within NSD3short^W284A^ domains (**Figures 6H and I**) compared to wild-type NSD3short.

To evaluate if the altered occupancy and localization of NSD3short^W284A^ on chromatin compared to wild-type NSD3short might impact chromosome folding we performed *in situ* Hi-C in cells induced to express NSD3short^W284A^ and identified 5,240,513,652 contacts (**Table S4**). Eigenvector analysis and compartment annotation indicated that EV1 values within NSD3short^W284A^ domains remained negative, with 93% of NSD3short domains overlapping B compartment intervals in MCF10A cells expressing NSD3short^W284A^ (**Figures S16C, S16D, and S16E**). Therefore, NSD3short^W284A^ domains not only arise from the B compartment but themselves fall within the B compartment. Global chromosome conformation was comparable between uninduced MCF10A cells, cells induced to express NSD3short, and cells expressing NSD3short^W284A^ (**Figures S17A and S17B**).

Visualizing chromatin interactions as Hi-C contact maps revealed that NSD3short^W284A^ did not promote chromatin interactions to the same extent as wild-type NSD3short (**Figure 6J**). Extending this analysis to all chromosomes using pileup analysis demonstrated an impaired ability of NSD3short^W284A^ to promote chromatin contacts genome-wide (**Figures 6K and S17C**). Contact frequency between NSD3short contact sites was decreased 2.29-fold in cells induced to express NSD3short^W284A^ compared to cells expressing wild-type NSD3short (**Figure 6L and S17D**). Though reduced compared to cells expressing wild-type NSD3short, contact frequency between NSD3short contact sites in cells expressing NSD3short^W284A^ was greater than that in uninduced MCF10A cells (**Figure 6M**). We detected chromatin contacts in cells expressing NSD3short^W284A^ between wild-type NSD3short peaks, but in aggregate, these contacts were weaker than in cells expressing wild-type NSD3short (**Figure S17E**). Furthermore, chromatin contacts common to cells expressing both variants of NSD3short, and contacts unique to cells expressing wild-type NSD3short were comparably weaker in cells expressing NSD3short^W284A^ (**Figure S17E**). The strength of contacts unique to cells expressing NSD3short^W284A^ was comparable between cells expressing wild-type NSD3short and cells expressing NSD3short^W284A^ (**Figure S17E**). Our detection of contacts unique to cells expressing NSD3short^W284A^, and the comparable frequencies for these contacts between cells expressing either variant of NSD3short, is likely due to denser Hi-C contact maps for cells induced to express NSD3short^W284A^ (**Table S4**).

We also inspected Hi-C contact maps to determine if NSD3short^W284A^ domains interacted with one another. However, we were unable to find interactions between NSD3short^W284A^ domains that were stronger in cells expressing NSD3short^W284A^ compared to uninduced MCF10A cells. To quantitatively assess if NSD3short^W284A^ domains contacted each other, we enumerated all possible NSD3short^W284A^ domain-domain contacts and then used pileups to plot the Hi-C contact frequency normalized for random polymer conformations of chromosomes in uninduced MCF10A cells, cells induced to express NSD3short, and cells expressing NSD3short^W284A^. Such pileup analysis did not detect any differences in normalized Hi-C contact frequency across conditions (**Figures 6N-O, S17F, and S17G**), indicating that NSD3short^W284A^ does not promote interactions between NSD3short^W284A^ domains. Though slight, normalized Hi-C contact frequency between all possible NSD3short^W284A^ domains was significantly reduced in cells expressing NSD3short^W284A^ compared to uninduced MCF10A cells (**Figure 6P**). We therefore find no evidence that NSD3short^W284A^ promotes chromatin contacts.

## DISCUSSION

Identification of NSD3 as a previously unrecognized mediator of chromosome folding is notable for several reasons: (1) cofactor stabilization of fusion oncoprotein-driven chromosome 3D organization and condensates, (2) multicomponent basis of chromosome compartmentalization, and (3) adaptor proteins that organize chromosomes into compartments independent of enzymatic activity. We identified a role for NSD3 in chromosome folding by investigating how the fusion oncoprotein BRD4-NUT, which causes a rare cancer, drives long-range chromatin interactions. Confirmation that NSD3short promotes megabase-scale chromatin interactions in fusion naïve cells highlights how mechanistic analysis of chromosome misfolding in a rare disease informs upon chromosome folding more broadly.

NSD3short^W284A^ occupying broad chromatin domains was unexpected but revealing. Inactivating NSD3short’s PWWP domain might have been expected to simply reduce NSD3short chromatin occupancy. Instead, the W284A mutation dramatically redistributes NSD3short across the genome, generating megabase-scale domains. However, we find no evidence of chromatin interactions between NSD3short^W284A^ domains. Thus, NSD3short^W284A^ identifies a separation-of-function mutant that uncouples chromatin occupancy from chromatin interactions. Wild-type NSD3short promotes interactions within the A compartment; NSD3short^W284A^ domains localize to the B compartment and do not interact. The capacity of a protein to promote chromatin contacts depends on the nuclear compartment in which it resides. Therefore, 1D localization along the chromatin fiber is not sufficient to predict 3D chromatin interactions.

Preferential interactions among *cis*-regulatory elements form microcompartments, which are remarkably resilient to acute transcriptional inhibition or cohesin loss^25^. Microcompartments, like A/B compartments, therefore, do not form by loop extrusion^25,28–32^. BRD4-NUT-driven megadomain-megadomain interactions are also inconsistent with loop extrusion and persist in the absence of transcription^34^; notably, the nuclear subcompartment formed by BRD4-NUT appears in healthy cells as a microcompartment^1,25^. The cofactor role of NSD3 in maintaining BRD4-NUT-driven compartmentalization further indicates that microcompartments are multicomponent. Multiple cofactors contribute distinct biochemical and structural functions, including BRD4-NUT, the histone acetyltransferase p300 recruited by BRD4-NUT to deposit H3K27ac^37,40,53,73^, NSD3, and H3K36me2 marked chromatin. Despite each factor providing unique capabilities, they collectively ensure the stability and resilience of nuclear organization.

BRD4-NUT and NSD3short both lack catalytic domains, yet both promote chromatin contacts and contribute to compartmentalization. Both function as adaptor proteins: BRD4-NUT recognizes acetylated histones through the bromodomains of BRD4 and stimulates p300 via NUT^37,38,40,41,53,73,74^; NSD3short binds methylated histone H3K36 through the PWWP domain^70,71^ and binds the extraterminal domain of BRD4^43,45^. We have demonstrated that BRD4-NUT drives *de novo* compartment formation^34^, and here show that NSD3 stabilizes compartmentalization. We propose that compartments, subcompartments, and microcompartments are assembled by adaptor proteins. Although adaptor proteins may lack enzymatic activity, they can provide the multivalent interactions needed to support long-range chromatin contacts. The identification of these two adaptor proteins that compartmentalize chromosomes strongly supports this model. LDB1, which establishes multi-enhancer networks but lacks enzymatic activity^75,76^, is another adaptor protein consistent with this framework.

Although NSD3short^W284A^ illustrates a separation of function between chromatin binding and chromatin interactions, we did not observe such an uncoupling between BRD4-NUT condensates and megadomain-megadomain interactions. In previous work, pharmacologic degradation of BRD4-NUT induced complete loss of condensates and megadomain-megadomain interactions^34^. Here, depletion of NSD3 from BRD4-NUT+ cells did not abolish condensates but instead resulted in condensate destabilization coincident with weakening, but not complete loss, of megadomain-megadomain interactions. The fold-change in BRD4-NUT enrichment in condensates was comparable to the decrease in contact frequency between megadomains following NSD3 depletion. These observations suggest that NSD3 is not a core driver but rather stabilizes condensate integrity and long-range chromatin contacts, consistent with an adaptor role that stabilizes both through multivalent interactions. Megadomain-megadomain interactions and condensates are therefore, at least at present, inseparable.

NSD3 depletion also downregulates megadomain genes, including *MYC*, which is important for NUT carcinoma oncogenesis^56^. Fewer genes are downregulated after loss of NSD3 than after loss of BRD4-NUT^34^. Nonetheless, slowed cell proliferation and cell differentiation accompany oncogene downregulation, weakening of megadomain-megadomain interactions, and condensate destabilization. NSD3 seems to add an additional, functional layer of regulation by ensuring appropriate valency between interacting megadomains within condensates.

Following NSD3 depletion in 10-15 cells, BRD4-NUT and NSD3 are less enriched in megadomains, whereas H3K27ac levels remain unchanged. Given that condensates are less stable and that megadomain-megadomain interactions decrease under these conditions, histone H3K27ac is unlikely to be the primary determinant of BRD4-NUT-driven condensate integrity and chromatin interactions. Instead, non-histone proteins, such as BRD4-NUT, NSD3, and perhaps other adaptor proteins, play prominent roles.

Amplification of NSD3 has been reported as an oncogenic driver not only in breast cancer^51,52,65^, but also in lung cancer^66,67^ and pancreatic ductal adenocarcinoma^68^. In addition to copy number gains, rearrangements of NSD3 generate oncogenic fusions, including NUP98-NSD3 in acute myeloid leukemia^77,78^ and NSD3-NUT in NUT carcinoma^46,48,49^. NSD3 functions oncogenically through both catalytic and non-catalytic mechanisms. The catalytic SET domain of NSD3 deposits H3K36me2 to drive lung squamous cell carcinoma^79^ and head and neck squamous cell carcinoma^80^. On the other hand, NSD3short has been shown to act within multiprotein complexes to mediate oncogenic transcription^45–47,81,82^. Here, we identify a role for NSD3 in chromosome folding in two cancer contexts. In NUT carcinoma, NSD3 deposits H3K36me2 in megadomains and stabilizes BRD4-NUT on chromatin, supporting oncogene expression by maintaining BRD4-NUT condensate integrity and megadomain-megadomain interactions. In MCF10A cells, catalytically inactive NSD3short promotes megabase-scale chromatin interactions coincident with a transition from an epithelial morphology to a more mesenchymal phenotype, which has been shown to drive tumor formation^69^. Our work expands our understanding of how NSD3 contributes to oncogenesis by regulating 3D genome organization.

Multiple strategies have been developed to therapeutically target NSD3^72,83–85^. An NSD3 inhibitor selectively targets the PWWP domain present in both NSD3long and NSD3short^72^. Because NSD3short^W284A^ redistributes on chromatin, it will be important to determine if PWWP-directed NSD3 inhibitors act, in part, by redistributing NSD3 across the genome. Therapeutic strategies to target BRD4-NUT using bromodomain and extraterminal (BET) protein inhibitors have shown limited clinical success because of severe adverse effects that result in a narrow therapeutic window^86,87^. Given a cooperative role of NSD3 in BRD4-NUT-driven nuclear organization, a strategy that combines BET inhibitors with NSD3-targeting agents may be beneficial in NUT carcinoma. Lastly, our results suggest that small molecules that target NSD3 may act, in part, by altering chromosome folding.

## ACKNOWLEDGMENTS

We thank Jingyi Yan for cloning dual-sgRNAs; Christopher A. French for providing the 10-15 cell line; Jeffrey A. Toretsky for providing the TC-797 cell line; Daniel R. Foltz for providing pA-MNase; and Hafiza Sara Akram, Joshua A. Riback, and Jingyi Yan for critical reading of the manuscript. This work was supported by grants from the National Cancer Institute (U01CA294062), NIH Office of the Director (DP5OD024587), the Cancer Prevention & Research Institute of Texas (RR210082), Alex’s Lemonade Stand Foundation for Childhood Cancer (22-25794), and The Welch Foundation (Q-2232-20250403) to K.P.E. Data analysis was performed on the HPC cluster that is managed by the Biostatistics and Informatics Shared Resource (BISR) at Baylor College of Medicine and supported by an NIH S10 Shared Instrument Grant S10-OD032185, NCI P30-CA125123 and institutional funds from the Dan L Duncan Comprehensive Cancer Center and Baylor College of Medicine. This project was supported by the Cytometry and Cell Sorting Core at Baylor College of Medicine with funding from the CPRIT Core Facility Support Award (CPRIT-RP240432) and the NIH (CA125123, OD036336, and OD038251), and the assistance of Joel M. Sederstrom. Y.C. was supported by a Bert W. O’Malley, MD Student Scholar in Medical Research Award sponsored by the Diana Helis Henry Medical Research Foundation and the Adrienne Helis Mavin Medical Research Foundation. C.D.R. was supported by a National Science Foundation Graduate Research Fellowship (DGE-2234667). K.P.E. is a CPRIT Scholar in Cancer Research. The content is solely the responsibility of the authors and does not necessarily represent the official views of the National Institutes of Health. This manuscript is the result of funding in whole or in part by the National Institutes of Health (NIH). It is subject to the NIH Public Access Policy. Through acceptance of this federal funding, NIH has been given a right to make this manuscript publicly available in PubMed Central upon the Official Date of Publication, as defined by NIH.

## AUTHOR CONTRIBUTIONS

Y.C., C.D.R., and K.P.E. conceived the study. Y.C., J.R.C., and K.P.E. designed experiments. Y.C., J.R.C., C.E.H., and C.P. developed novel cell lines. Y.C. performed confocal microscopy experiments. Y.C. and J.R.C. performed Hi-C experiments. Y.C., J.R.C., and I.D.Y. performed CUT&RUN experiments. Y.C. and J.R.C. performed RNA-seq experiments. Y.C., K.S., and K.P.E. designed data analysis. Y.C. performed confocal microscopy analysis. Y.C., K.S., and K.P.E. performed Hi-C data analysis. Y.C., K.S., and K.P.E. performed CUT&RUN data analysis. K.S. performed RNA-seq data analysis. Y.C., J.R.C., K.S., and K.P.E. discussed and interpreted the results. Y.C., J.R.C., K.S., and K.P.E. wrote the manuscript.

## DECLARATION OF INTERESTS

The authors declare no competing interests.

## METHODS

### Cell culture

TC-797 cells were a kind gift from Jeffrey A. Toretsky (Georgetown University, Washington, DC). 10-15 cells were a kind gift from Christopher A. French (Brigham and Women’s Hospital, Boston, MA). TC-797 and 10-15 cells were grown in DMEM with 4.5 g/L glucose and sodium pyruvate without L-glutamine (Corning #15013CV), 10% Fetal Bovine Serum (GenDEPOT #F0900-050), 1% GlutaMAX (Life Technologies #35050-061), 100 units/mL penicillin, 100 μg/mL streptomycin (ThermoFisher #15070-063), and 1.25 μg/mL plasmocin prophylactic (Invivogen #ant-mpp) at 37° C with 5% CO_2_. Lenti-X 293T cells (Takara Bio #632180) were cultured in DMEM supplemented with 10% fetal bovine serum, 1% GlutaMAX, 100 units/mL penicillin, and 100 μg/mL streptomycin at 37° C with 5% CO_2_. MCF10A cells were cultured in DMEM/F12 (Corning #10-092-CV) supplemented with 5% donor equine serum (HyClone #SH30074.03), 20 ng/mL epidermal growth factor (EGF) (Peprotech #AF-100-15), 0.5 mg/mL hydrocortisone (Sigma #H0888), 100 ng/mL cholera toxin (Sigma #C8052), 10 mg/mL insulin (Sigma #I1882), 100 units/mL penicillin, 100 mg/mL streptomycin, and 1.25 μg/mL plasmocin prophylactic (Invivogen #ant-mpp) at 37° C with 5% CO_2_. Cell lines regularly tested negative for mycoplasma with the Mycoplasma PCR Detection Kit (ABM #G238). *Drosophila* Kc167 cells (Drosophila Genomics Research Center #1) were cultured in CCM3 media (Cytiva #SH30065.02) at 25° C.

### Lentivirus transfer plasmids

Zim3-dCas9 plasmid (pHR-UCOE-SFFV-Zim3-dCas9-P2A-Hygro) was purchased from Addgene (#188768). Dual-sgRNA design and cloning were performed according to the protocol described by Replogle et al. (2022)^61^. The dual sgRNA targeting NSD3 plasmid (dual-sgNSD3) contains mU6-GGGCCGAGGGGGCTGTGCAC-sgRNA_scaffold-hU6-GGCCTGGGTAGCACGGTCCT-sgRNA_scaffold cloned in a hU6-sgRNA EF1Alpha-puro-T2A-BFP plasmid (Addgene # 60955). The dual sgRNA non-target control plasmid (dual-sgNT) contains mU6-GAGGTGGCGGACGCCCGCAT-sgRNA_scaffold-hU6-GCTCTCTGTATCATCTTGCC-sgRNA_scaffold cloned in a pU6-sgRNA EF1Alpha-puro-T2A-BFP plasmid (Addgene # 60955). The protein coding sequence for the short isoform of NSD3 (Ensembl transcript ID ENST00000316985) was codon optimized using the GeneArt GeneOptimizer software (Thermo Fisher) and then an 11 amino acid HiBiT epitope tag plus a Gly-Ser-Gly linker was added to the N-terminus. HiBiT-Gly-Ser-Gly-NSD3short was placed downstream of a TRE3GS promoter and, along with the Tet-On 3G transcriptional transactivator under the control of a hPGK promoter, synthesized (Genewiz) in a Sleeping Beauty vector. The synthesized construct was then subcloned into the pCDH-EF1α-MCS-(PGK-Puro) vector (System Biosciences #CD810A-1) by replacing the EF1α-MCS-(PGK-Puro) sequence to generate plasmid pCDH-TetOne-BSD-NSD3short. The Trp284Ala mutation was generated in the NSD3short coding sequence by mutagenesis of the Trp-encoding codon (TGG) to an Ala-encoding codon (GCC) to generate plasmid pCDH-TetOne-BSD-NSD3shortW284A.

### Lentivirus packaging and concentration

7,500,000 Lenti-X 293 cells were seeded in a 10 cm cell culture dish 24 hours before packaging. The next day, cells were transfected with 5 µg lentiviral transfer plasmid along with 4 µg packaging plasmid psPAX2 (Adgene #12260) and 1 µg pMD2.G (Addgene #12259) using the TransIT-Lenti reagent (MirusBio #MIR 6604) according to the TransIT-Lenti manufacturer’s instructions. Cells were incubated at 37° C with 5% CO₂ for 48 hours. The lentivirus-containing supernatant was collected by filtration through a 0.45 µm PVDF membrane (VWR #76478-988) and was concentrated by adding one volume of 4x Lentivirus Concentrator Solution (40% (w/v) PEG-8000 and 1.2 M NaCl in PBS pH 7.4) to three volumes of supernatant. After overnight incubation at 4° C with gentle rotation, the concentrated lentivirus-containing supernatant was centrifuged at 1,600 x g for 60 minutes at 4°C, and the resulting pellet was resuspended in Lentivirus Stabilizer solution (Origene # TR30039) at 1/10 the original culture supernatant volume. Lentivirus titer was estimated using Lenti-X GoStix Plus (Takara Bio #631280). Aliquots were prepared and stored at −80° C.

### CRISPRi cell line engineering and validation

#### Lentivirus transduction of Zim3-dCas9

50,000 TC-797 cells or 10-15 cells were transduced with pHR-UCOE-SFFV-Zim3-dCas9-P2A-Hygro (Addgene #188768)^61^ lentivirus particles corresponding to 6.25 ng viral p24 or 1.5625 ng viral p24, respectively. Before transduction, complete growth medium lacking antibiotics was supplemented with polybrene at a final concentration of 10 μg/mL. Lentiviral aliquots were thawed on ice and used immediately to avoid multiple freeze-thaw cycles. For transduction, 50,000 cells were first seeded per well in a 6-well plate in 1 mL of complete media containing polybrene. The thawed lentivirus was drop-added to the 6-well plate and mixed by gently swirling. After 48 hours of incubation with the lentivirus, the media was replaced with fresh complete growth media containing 200 μg/mL hygromycin (Corning #31282-04-9) for antibiotic selection. The hygromycin treatment was refreshed every 48-72 hours. TC-797 Zim3-dCas9 cells and 10-15 Zim3-dCas9 cells were cryopreserved 21 days after lentivirus transduction. Estimated Multiplicity of Infection (MOI) for TC-797 Zim3-dCas9 cells and 10-15 Zim3-dCas9 cells was 0.00082 and 0.00702, respectively. TC-797 Zim3-dCas9 and 10-15 Zim3-dCas9 cells were further maintained with complete growth media containing 100 μg/mL hygromycin (Corning #31282-04-9). For simplicity, when used for CRISPRi experiments, these cell lines are referred to as TC-797 or 10-15 throughout the manuscript.

#### Activity-based sorting of Zim-dCas9 cells

TC-797 and 10-15 cells stably expressing the Zim3-dCas9 CRISPRi effector were transiently transfected with 50 nM sgRNA duplex targeting non-essential cell surface protein CD81 (sgRNA-CD81) (mixture of Alt-R CRISPR-Cas9 crRNA-CD81 and Alt-R CRISPR-Cas9 tracrRNA ATTO 488 (IDT #10007810)) using Lipofectamine RNAiMAX Reagent (Invitrogen #13778-100) 24 hours after seeding, following the manufacturer’s instructions. The sgRNA duplex with a non-targeting sequence (sgRNA-NT) was used as a control. Cells were detached and collected after 72 hours (TC-797) and 96 hours (10-15) of incubation with sgRNA duplex. Cells were then stained with APC-CD81 antibody (BioLegend #349509) at a 1:500 dilution for 30 minutes at room temperature on a nutator, protected from light. After staining, cells were washed once with FACS buffer and centrifuged at 300 x g for 5 min. Cells were resuspended in FACS buffer with 55 ng/mL DAPI (Thermo Fisher #D3571). TC-797 cells were analyzed and sorted by BD FACSDiscover S8 Cell Sorter. 10-15 cells were analyzed and sorted by BD FACSAria III Cell Sorter. APC signal (emission wavelength = 660 nm) was used to measure the CD81 expression. We used the APC signal intensity from sgRNA-NT to define the threshold of the gate of CD81-positive and CD81-negative cells. We then analyzed the APC signal of cells transfected with sgRNA-CD81 duplex and sorted and collected the cells within the CD81 negative gate. The sorted cells were further expanded and depletion efficiency was validated by lentiviral transduction of sgRNA-CD81 (see below). After validation, the sorted Zim-dCas9 cells were cryopreserved in liquid nitrogen and maintained in the complete medium containing 100 μg/mL hygromycin.

#### Cell cytometry to validate Zim-dCas9 cells

Mammalian cells were detached from cell culture dishes and harvested at room temperature. 1,400,000 cells were used for each sample and resuspended with 500 μL FACS buffer (1% fetal bovine serum (FBS), 10 mM HEPES-NaOH pH 7.5, and 1 mM EDTA in 1X PBS, sterile-filtered). 1 μL primary antibody was added for a 1:500 dilution. Primary antibodies were APC-CD81 antibody (BioLegend #349509), and APC Mouse IgG1 κ Isotype Ctrl (FC) Antibody (BioLegend # 400122). Cells were incubated for 30 min at room temperature on a nutator, protected from light. After staining, cells were washed once with 500 μL FACS buffer and centrifuged at 300 x g for 5 min. Cells were resuspended in 250 μL FACS buffer with 55 ng/mL DAPI (Thermo Fisher #D3571). Flow cytometry was performed using a BD FACSCanto II System, and data were analyzed by the BD FACSCanto Clinical Software and FlowJo software (version: 10.0.0).

### Lentivirus transduction of sgRNA

500,000 TC-797 cells or 400,000 10-15 cells, respectively, were transduced with 25 μL of sgRNA lentivirus particles in a 60 mm plate. Before transduction, complete growth medium lacking antibiotics was supplemented with polybrene at a final concentration of 10 μg/mL. Lentiviral aliquots were thawed on ice and used immediately to avoid multiple freeze-thaw cycles. For transduction, cells were first seeded in a 60 mm plate in 4.25 mL of complete growth media containing polybrene. The thawed lentivirus was diluted with 500 μL complete growth medium containing polybrene, drop-added to the 60 mm plate, and mixed by gently swirling. After 48 hours of incubation with the lentivirus, the media was replaced with fresh complete media containing 400 ng/mL puromycin (InvivoGen #ant-pr-1) for antibiotic selection. The puromycin treatment was refreshed every 48-72 hours.

### Lentivirus transduction of NSD3short variants

Lentivirus particles corresponding to 150 ng viral p24 were diluted in 500 μL complete growth media lacking antibiotics supplemented with 10 μg/mL polybrene and added to individual wells of a 6-well plate. 50,000 MCF10A cells in 1 mL polybrene-containing media were added dropwise to lentivirus-containing wells. After 72 hours, the media was replaced with fresh complete growth media containing 5 μg/mL blasticidin. Blasticidin treatment was refreshed every 48-72 hours for 14 days in order to maintain antibiotic selection. MCF10A(NSD3short) and MCF10A(NSD3short^W284A^) cell lines were expanded and cryopreserved. Estimated MOI for NSD3short and NSD3short^W284A^ lentiviruses was 0.002.

### NSD3short variant induction

MCF10A cells were treated with either 30 ng/mL doxycycline (Fisher Scientific #AC446060050) to induce NSD3short expression or 25 ng/mL doxycycline to induce NSD3short^W284A^ expression for a total of 5 days. On day 3, treated cells were given fresh MCF10A complete growth media containing 30 ng/mL or 25 ng/mL doxycycline for NSD3short or NSD3short^W284A^, respectively. Doxycycline was solubilized in sterile water.

### Wright-Giemsa staining

150,000 TC-797 cells or 300,000 10-15 cells 11 days after lentiviral transduction of sgRNA were seeded onto a KOH-treated and 0.01% poly-lysine (Sigma #P4707-50ML) coated cover glass (Fisherfinest #12-548-A 18X18-1) in wells of a 12-well plate 24 hours before the staining. The Camco Stain Pak (Cambridge Diagnostic Products #702) was used for staining. For TC-797 cells, the cover glass was stained by dipping 4 times into Camco Stain Pak Fixative Solution (1 sec for each dip), followed by dipping twice into Camco Stain Pak Solution I (1 sec for each dip) and then dipping once into Camco Stain Pak Solution II. For 10-15 Zim3-dCas9 cells, the cover glass was stained by dipping 4 times into Camco Stain Pak Fixative solution (1 sec for each dip), followed by dipping three times into Camco Stain Pak Solution I (1 sec for each dip) and then dipping three times into Camco Stain Pak Solution II.

15,000 MCF10A cells were seeded onto a KOH-treated and 0.01% poly-lysine coated cover glass in wells of a 12-well plate containing 1 mL fresh MCF10A complete growth media. Cells were either mock-treated with sterile water or treated with doxycycline for 5 days to induce NSD3short or NSD3short^W284A^ expression as described above. The cover glass was stained by dipping five times into Camco Stain Pak Fixative solution (1 sec for each dip), followed by dipping once into Camco Stain Pak Solution I (1 sec for each dip) and then dipping once into Camco Stain Pak Solution II.

For all cell types, cover glass was rinsed by dipping into distilled water for a few seconds. After the cover glass dried, cover glass was mounted onto glass slides using Permount Mounting Medium (Fisher Scientific #SP15-100). The cover glass was imaged by a Leica MC170HD camera and processed by Fiji ImageJ^88^.

### CellTiter Glo assay

Cell viability of NSD3-depleted cells was assessed 8 days after lentiviral transduction with the sgRNA using the CellTiter-Glo Luminescent Cell Viability Assay (Promega, #G7572). On day 5 of sgRNA transduction, 900 TC-797 cells or 5,000 10-15 cells were placed in one well of a 96-well plate. The cell viability was collected starting from 16 hours after the cells were placed and defined as the 0-hour time point. Viability was then measured every 24 hours. Plates containing cells were equilibrated to room temperature for 1 hour. The culture medium was removed and replaced with 100 µL of fresh DMEM (room temperature) in each well. Subsequently, 100 µL of CellTiter-Glo reagent was added to each well. Blank wells contained 100 µL DMEM and 100 µL CellTiter-Glo reagent as a background control. Plates were mixed on an orbital shaker at 450 rpm for 2 minutes at room temperature. After mixing, 190 µL of the lysate was transferred to a Costar opaque-walled black 96-well plate (Corning #3631). The plate was incubated at room temperature for 10 minutes to stabilize the luminescent signal. Luminescence was measured using a Tecan Infinite M200 plate reader with the following parameters: integration time of 1,000 ms and automatic attenuation. For each sample, four technical replicates were collected. Data was analyzed by GraphPad Prism software (version 10). Three biological replicates from each condition were used for statistical analysis.

### Growth curves

100,000 MCF10A cells were seeded into wells of multiple 6-well plates containing 2 mL of MCF10A complete growth media and incubated overnight at 37° C. The following day (Day 0), cells were mock treated with sterile water or treated with doxycycline to induce NSD3short or NSD3short^W284A^ expression as described above. To obtain cell counts, cells from each experimental condition were detached using 250 μL TrypLE (Life Technologies #12605-028), mixed with 250 μL PBS, and then 200 μL of each sample was counted on a Vi-Cell BLU (Beckman Coulter) machine. For each experimental condition, cell counts were determined daily for up to 5 days of doxycycline treatment. Three replicates from each condition were used for statistical analysis.

### Immunofluorescence

Immunofluorescence was performed with DAPI counterstaining. Primary antibodies used were NUT rabbit monoclonal antibody (Cell Signaling Technologies #3625, lot 8, RRID: AB_2066833), H3K27ac mouse monoclonal antibody (Active Motif #39685, RRID: AB_2722569), WHSC1L1 rabbit monoclonal antibody (Cell Signaling Technologies #92056, lot 1, RRID: AB_2800178. Secondary antibodies included Goat anti-Rabbit IgG (H+L) Highly Cross-Adsorbed Secondary Antibody, Alexa Fluor 488 (Thermo Fisher #A-11034, RRID: AB_2576217) and Goat anti-Mouse IgG (H+L) Highly Cross-Adsorbed Secondary Antibody, Alexa Fluor 647 (Thermo Fisher #A-21236, RRID: AB_2535805).

High-precision cover glass (Bioscience Tools #CSHP-No1.5-18) was cleaned by submerging the cover glass in 1 M KOH and sonicating for 15 minutes in an ultrasonic cleaner. Cover glasses were washed 5 times with ultrapure water, once with 70% ethanol, and then air-dried in a tissue culture hood. One cover glass was placed in each well of a 12-well culture dish. Cover glasses were treated for 5 min with 0.01% poly-lysine (Sigma #P4707-50ML) while at room temperature. Cover glasses were washed three times with sterile water, air-dried, and sterilized by UV light in the tissue culture hood.

Cells were seeded in wells 2-24 hours before fixation, adjusting for optimal density. Cells with 1 mL complete media were fixed by adding 1 mL of 8% EM-grade paraformaldehyde (Electron Microscopy Sciences #15714) in PBS to give a final concentration of 4% paraformaldehyde and then incubated for 20 min at room temperature while rocking. Cells were washed three times with 0.5 mL of PBS and then permeabilized with 1 mL of 1% Triton X-100 in PBS while rocking for 20 min at room temperature. Cells were washed three times with 1 mL of PBS, and then blocked for 1 hour at room temperature with blocking buffer (5% non-fat milk with 0.02% sodium azide in PBS) while rocking. Primary antibodies were diluted 1:1,000 using blocking buffer, and 0.5 mL of diluted primary antibody was added to each well and then incubated with the fixed, permeabilized cells with rocking overnight at 4° C. Cells were washed three times with 1 mL of PBS. Secondary antibodies, diluted 1:1,000 with blocking buffer, were added after the last wash and incubated with rocking for 1 hour at room temperature, protected from light. While maintaining protection from light, cells were washed three times with 1 mL of PBS, followed by counterstaining with 300 nM DAPI dilactate (Fisher #D3571) in PBS for 10 min at room temperature with rocking. Two final rounds of washing were done with 1 mL of PBS, and the cover glass was mounted on microscope slides using ProLong Diamond (Thermo Fisher #P36965). Slides were cured overnight, protected from light.

Slides were imaged on a CSU-W1 Yokogawa Spinning Disk Field Scanning Confocal System using a Nikon Plan APO Lambda 100x/1.45 Oil OFN25 DIC N2 objective. Images were acquired with Nikon NIS-Elements Advanced Research software at an image bit depth of 16 bits. Images were collected as Z stacks on a Photometrics Prime 95B Scientific CMOS Camera, one for each imaging channel, with a 110 nm pixel size. Appropriate filters for fluorochromes Alexa Fluor 488, Alexa Fluor 640, and 405-DAPI were used. Images were projected along the Z axis using Nikon’s Extended Depth of Focus (EDF) algorithm, the BRD4-NUT channel was pseudocolored magenta, the histone H3K27ac channel was pseudocolored green, the NSD3 channel was pseudocolored red, and the DAPI channel was pseudocolored blue each using a linear LUT that covered the full range of the data, the channels merged, and the image converted to TIFF format using Nikon NIS-Elements Advanced Research software. The converted TIFF images served as the raw data for subsequent analyses. For visualization purposes, images of the same channel were linearly rescaled to the same minimum and maximum across experimental conditions using Fiji ImageJ^88^.

### Immunoblotting

Primary antibodies were NUT (C52B1) Rabbit mAb (Cell Signaling Technologies #3625, lot 8, RRID: AB_2066833) diluted 1:5,000; Acetyl-Histone H3 (Lys27) (D5E4) XP Rabbit mAb (Cell Signaling Technology #8173S, lot 9, RRID: AB_10949503) diluted 1:1,000; WHSC1L1 rabbit monoclonal antibody (Cell Signaling Technologies #92056, lot 1, RRID: AB_2800178) diluted 1:10,000; Di-Methyl-Histone H3 (Lys36) (C75H12) rabbit monoclonal antibody (Cell Signaling Technologies #2901, lot 5, RRID: AB_1030983) diluted 1:1,000; and anti-HiBiT mouse monoclonal antibody (Promega #N7200, RRID: AB_3665694) diluted 1:10,000. Loading control primary antibodies were human anti-GAPDH antibody, clone AbD22549_hIgG1 (Bio-Rad #HCA272) diluted 1:5,000; and histone H3 (96C10) mouse monoclonal antibody (Cell Signaling Technologies #3638, RRID: AB_1642229) diluted 1:5000. Secondary antibodies were goat anti-rabbit IgG AzureSpectra 800 (Azure Biosystems # AC2134) diluted 1:5,000; goat anti-mouse IgG AzureSpectra 700 (Azure Biosystems # AC2129) diluted 1:5,000; and goat anti-human IgG AzureSpectra 490 (Azure Biosystems # AC2208) diluted 1:5,000.

1,000,000 uncross-linked cells, collected simultaneously with those for CUT&RUN or cross-linked for Hi-C, were collected in a 1.5 mL tube, centrifuged at 300 x g for 5 min at 4° C, the supernatant discarded, and the cell pellet resuspended in 1 mL of ice-cold PBS. Cells were then centrifuged at 300 x g for 5 min at 4° C, the supernatant discarded, and the cell pellet resuspended in 50 μL of RIPA lysis buffer (50 mM Tris-HCl pH 8.0, 150 mM NaCl, 0.1% Triton X-100, 0.05% Sodium deoxycholate, 1 mM EDTA, 0.01% SDS). Cells were lysed at 95° C for 5 min, cooled to room temperature, and whole-cell extracts were stored at -80° C. Cell extracts were thawed on ice. Nucleic acid was removed by treating samples with 250 U of Recombinant Dr. Nuclease (Syd Labs #BP4200) and incubating at 37° C for 10 min. Protein concentration was quantified using a Bio-Rad DC Protein Assay (Bio-Rad#5000111). 10 μg of protein in 1X Laemmli Sample Buffer (Bio-Rad #161-0737 or Bio-Rad #161-0747) with 2.5% 2-mercaptoethanol (Sigma #M6250) was incubated at 95° C for 2 min, and then run on a 4%–20% Mini-PROTEAN TGX gel (Bio-Rad #4561093 or Bio-Rad #4561096) at 130 V. Samples were transferred to a 0.45 μm LF PVDF membrane using a Bio-Rad Trans-Blot Turbo Transfer System and Trans-Blot Turbo TransferPack (Bio-Rad # 1704274). For the blotting of BRD4-NUT, proteins were transferred from gels to LF PVDF membranes using a wet transfer system in 1x transfer buffer (25 mM Tris, 192 mM glycine, 20% methanol, pH 8.3) at 0.09 A for 18 hours at 4°

C. The TotalStain Q fluorescent total protein stain kit (Azure Biosystems #AC2225) was used to ensure successful transfer according to the manufacturer’s instructions. The membrane was blocked with blocking buffer (TBST + 5% non-fat dry milk) for 30 min at room temperature while rocking. The blocking buffer was discarded, and the primary antibody (diluted in blocking buffer) was added and incubated overnight at 4 °C while rocking. Buffer was discarded, and the membrane was washed three times for 10 min at room temperature with TBST (20 mM Tris-HCl pH 7.5, 150 mM NaCl, 0.1% Tween 20) while rocking. Secondary antibody (diluted in blocking buffer) was added, incubated for 1 hour while rocking at room temperature, and then washed three times for 10 min at room temperature with TBST while rocking. After removing the TBST, the membrane was washed with PBS for 5 min and immersed in 100% methanol for 30 seconds. After drying, the blot was imaged with an Azure Biosystems Sapphire FL Biomolecular Imager (Azure Biosystems, Dublin, CA) and quantified with AzureSpot Pro software (Azure Biosystems, Dublin, CA).

### CUT&RUN library preparation

Primary antibodies were NUT (C52B1) rabbit monoclonal antibody (Cell Signaling Technologies #3625, lot 8, RRID: AB_2066833), Acetyl-Histone H3 (Lys27) (D5E4) XP rabbit monoclonal antibody (Cell Signaling Technology #8173S, lot 9, RRID: AB_10949503), WHSC1L1 rabbit monoclonal antibody (Cell Signaling Technologies #92056, lot 1, RRID: AB_2800178), H3K36me2 rabbit monoclonal antibody (T.571.7) (Invitrogen # MA5-14867), and anti-HiBiT mouse monoclonal antibody (Promega #N7200, RRID: AB_3665694). Negative control primary antibody was rabbit anti-mouse IgG H&L (Abcam #ab46540). NUT expression is restricted to testis^89^, therefore we employed the above anti-NUT antibody for BRD4-NUT CUT&RUN, as done previously for ChIP-seq^34,53^.

Mammalian cells were detached from cell culture plates using TrypLE Express (Thermo Fisher #12605028) and harvested at room temperature. Cells were centrifuged at 300 x g for 5 min at room temperature and washed twice with Wash buffer (20 mM HEPES-NaOH pH 7.5, 150 mM NaCl, 0.5 mM spermidine (Sigma #S0266-1G), 1x cOmplete EDTA-free Protease Inhibitor Cocktail (Sigma-Aldrich #11873580001)). Concanavalin A-coated magnetic beads (Bangs Laboratories #BP531) were washed twice with 10 times the volume of Binding buffer (20 mM HEPES-KOH pH 7.9, 10 mM KCl, 1 mM CaCl_2_, 1 mM MnCl_2_) and the beads then resuspended with the same volume of Binding buffer after the last wash. For each sample, 400,000 mammalian cells were bound to 10 μL concanavalin A-coated magnetic beads by adding the cells in the wash buffer to the beads and mixing on an end-over-end rotator for 5-10 min. Tubes containing bead-bound cells were placed on a magnetic stand, the liquid removed from the bead-bound cells, and then bead-bound cells were resuspended in 50 μL ice-cold Antibody buffer (20 mM HEPES-NaOH pH 7.5, 150 mM NaCl, 0.5 mM spermidine, 0.1% digitonin, 1x protease inhibitor cocktail, 2 mM EDTA, 0.1 mg/mL BSA, 100 nM trichostatin A (Sigma #T8552)).

For H3K36me2 and IgG antibodies:

1 μL of SNAP-CUTANA K-MetStat Panel (Epicypher #19-1002) was added to each sample, followed by 1 μL of primary antibody in sequential order. After mixing by gently pipetting, the sample was nutated overnight at 4° C.

For NUT, Acetyl-Histone H3 (Lys27), WHSC1L1, and HiBiT antibodies: 1 μL of primary antibody was added to each sample. After mixing by gently pipetting, the sample was nutated overnight at 4° C.

For HiBiT antibody:

Following overnight nutation at 4° C, only samples containing the mouse monoclonal HiBiT antibody were removed from 4° C and placed on a magnetic stand. After withdrawing liquid, samples were washed twice with 100 μL antibody buffer. Following removal of all liquid after the second wash, samples were removed from the magnetic stand, resuspended with 50 μL antibody buffer + 1 μL rabbit anti-mouse IgG (Abcam #ab46540), and incubated at 4° C for 1 hour on a nutator.

Proceeded identically for all samples:

Samples were washed twice with ice-cold Dig-wash buffer (20 mM HEPES-NaOH pH 7.5, 150 mM NaCl, 0.5 mM spermidine, 0.1% digitonin, 1x protease inhibitor cocktail) and incubated with pA-MNase (700 ng/mL final concentration; a kind gift from Daniel R. Foltz) diluted in Dig-wash buffer for 1 hour at 4° C. After two additional washes with Dig-wash buffer, samples were resuspended with 50 μL ice-cold Dig-wash buffer. Chromatin digestion was initiated by adding 1 μL of 100 mM CaCl₂, gently mixed, and nutating for 30 min at 4°C. The reaction was stopped by adding 50 μL of 2x STOP buffer (340 mM NaCl, 20 mM EDTA, 4 mM EGTA, 0.1% digitonin, 50 μg/mL glycogen (Thermo Fisher #R0561), 100 μg/mL RNase A (Thermo Fisher #EN0531)). CUT&RUN fragments were released by incubating samples at 37° C for 10 min in a thermocycler with a heated lid (42° C). Supernatants were collected after magnetic separation. DNA was purified using a DNA Clean & Concentrator-5 kit (Zymo Research #D4004) following the manufacturer’s instructions. Purified DNA was quantified using a DeNovix High Sensitivity dsDNA Assay (DeNovix #KIT-DSDNA-HIGH-2) and DeNovix DS-11 FX+ fluorometer.

10 ng of DNA from each sample was prepared for high-throughput sequencing following the directions for ‘‘NEBNext End Prep’’ and “Adaptor Ligation’’ in the NEBNext Ultra II DNA Library Prep Kit for Illumina (New England Biolabs #E7645). After the cleanup of adaptor-ligated DNA without size selection by CleanNGS beads (Bulldog Bio #CNGS050), the sample was eluted and reclaimed using 8.5 μL 0.1x TE buffer (1 mM Tris-HCl pH 8.0, 0.1 mM EDTA). PCR was set up using 7.5 μL adaptor-ligated DNA, 5 µL of premixed Index/Universal Primer from the NEBNext Multiplex Oligos for Illumina (96 Unique Dual Index Primer Pairs) (NEB #E6440), and 12.5 μL NEBNext Ultra II Q5 Master Mix. The sample was run on a thermocycler at 98° C for 45 s followed by 15 cycles of 98° C for 15 s, 60° C for 10 s, followed by 1 cycle at 72° C for 1 min, and finally held at 4 °C. Two cleanups without size selection were applied to remove excess primers, adaptors, and adaptor dimers. For the first cleanup, 25 μL of CleanNGS beads was added to 25 μL of the PCR sample. After mixing and incubation for 5 min, the beads were collected in a magnetic stand for 5 min, and the supernatant was discarded. The beads were washed twice with 80% ethanol for 30 seconds while on the magnet stand. Beads were air-dried on the magnet, resuspended in 25 μL of 0.1x TE buffer for the second round of cleanup. 25 μL of SPRI buffer (10 mM Tris-HCl pH 8.0, 1 mM EDTA, 2.5 M NaCl, 20% PEG 8000, 0.05% Tween-20) was added to the 25 μL resuspended beads in 0.1x TE buffer. After mixing and incubation for 5 min, the beads were collected in a magnetic stand for 5 min, and the supernatant was discarded. The beads were washed twice with 80% ethanol for 30 seconds while on the magnet stand. Beads were air-dried on the magnet, resuspended in 16 μL of 0.1x TE buffer for eluting DNA, and 15 μL of the supernatant containing the eluted DNA transferred to a fresh tube.

Final sample concentration and quality were determined with Agilent 4200 TapeStation D1000 ScreenTape (Agilent, Santa Clara, CA) to determine sample concentration (region analysis from 140-1000bp) and sample quality. DNA was paired-end sequenced, 150 cycles each read, on an Illumina NovaSeq X Plus instrument. CUT&RUN was performed in duplicate, where each biological replicate represents cells from a different passage as well as treated independently.

### Cell crosslinking for Hi-C

MCF10A cells and transduced NUT carcinoma cells were treated with TrypLE Express and incubated for 10-15 min to detach adherent cells. TrypLE Express was quenched by adding media without serum. One day prior to crosslinking, 2.5x10^7^ *Drosophila* Kc167 cells were seeded onto 10 cm dishes. Kc167 cells were detached from the plate by blowing medium at the cells using a serological pipette. Kc167 cells were then centrifuged at 500 x g for 5 min and resuspended in mammalian media lacking serum to reach a density of 1x10^7^ cells/mL. Multiples of 5 million mammalian cells were added to conical tubes pre-treated with 0.05% BSA, and Kc167 cells were added to the tube to achieve a spike-in ratio of 2:1 (mammalian cells:fly cells). Cells were then centrifuged at 500 x g for 5 min and resuspended at a density of 1x10^6^ cells/mL in mammalian media without serum three times to remove traces of serum. Cells were resuspended in PBS + 0.05% BSA at a density of 1x10^6^ cells/mL and crosslinked by adding 32% EM-grade paraformaldehyde (Electron Microscopy Sciences #15714) to a final concentration of 1% and incubated at room temperature for 10 min while constantly rotating end-over-end. Paraformaldehyde was quenched by adding 2.5 M glycine to a final concentration of 0.15 M while constantly rotating end-over-end for 5 min at room temperature. Cells were then centrifuged at 3,000 x g for 5 min and resuspended at a density of 1x10^6^ cells/mL in PBS + 0.05% BSA two times to remove traces of paraformaldehyde. Cells were then centrifuged at 3,000 x g for 5 min and resuspended at a density of 1x10^6^ cells/mL in PBS + 0.05% BSA containing 3 mM disuccinimidyl glutarate (DSG) (Proteochem #c1104) and incubated at room temperature for 45 min while constantly rotating end-over-end. DSG was quenched by adding 2.5 M glycine to a final concentration of 0.15 M while constantly rotating end-over-end for 5 min at room temperature. Cells were then centrifuged at 3,000 x g for 5 min at 4° C, resuspended in 1 mL of ice-cold PBS + 0.05% BSA per 5 million mammalian cells and subsequently pelleted. Cell pellets were flash frozen in dry ice/ethanol and stored at -80° C.

### Hi-C library preparation

Hi-C libraries were prepared using minor modifications to the previously described *in situ* Hi-C protocol^64^. Five million cross-linked mammalian cells (with Kc167 cell spike-in), thawed on ice, were resuspended in 250 μL of Hi-C lysis buffer (10mM Tris-HCl pH 8.0, 10 mM NaCl, 0.2% Igepal CA-630) with 50 μL of 100x Protease Inhibitor Cocktail (Sigma #P8340), and incubated on ice for 15 min. The lysate was then centrifuged at 2,500 x g for 5 min at 4° C and the supernatant discarded. Nuclei were washed with 500 μL of ice-cold Hi-C lysis buffer, centrifuged at 2,500 x g for 5 min at 4° C, and the supernatant discarded. The pellet was gently resuspended in 50 μL of 0.5% SDS, incubated at 62° C for 10 min, and immediately cooled on ice. SDS was neutralized by adding 145 μL of water and then 25 μL of 10% Triton X-100 to the tube, mixed gently by inversion, and incubated for 15 min at 37° C. A 10% aliquot of the sample was taken as an undigested control. 22 μL of 1.1% Triton X-100 was added to the sample to compensate for the loss in volume. 25 μL of 10x NEBuffer 2 (New England Biolabs) and 4 μL of 25 U/mL MboI (New England Biolabs #R0147) were added to the sample and then gently inverted to mix. The sample was incubated overnight at 37° C in a ThermoMixer (Eppendorf) set to 750 rpm, 15 s on, 4 min 45 s off, to digest the chromatin.

To inactivate MboI, samples were incubated at 62° C for 20 min while shaking at 750 rpm, 15 s on, 4 min 45 s off in a ThermoMixer, and immediately cooled on ice. A second 10% aliquot was taken as a digested control. 24.9 μL of 1x NEBuffer 2 was added to the sample to compensate for the loss in volume. Restriction fragment overhangs were filled in and biotin labeled by adding 50 μL of fill-in master mix (24.75 μL water, 16.5 μL 1 mM Biotin-14-dATP (Axxora #JBS-NU-835-BIO14-L), 1.65 μL 10 mM dTTP, 1.65 μL 10 mM dGTP, 1.65 μL 10 mM dCTP, 8.8 μL 5 U/mL DNA Polymerase I, Large (Klenow) Fragment (New England Biolabs #M0210)) and incubated at 37° C for 45 min while shaking at 7 50 rpm, 15 s on, 4 min 45 s off in a ThermoMixer. 900 μL of ligation master mix (702.45 μL water, 126 μL 10x T4 DNA Ligase Buffer (New England Biolabs), 105 μL 10% Triton X-100, 6.3 μL 20 mg/mL BSA, 5.25 μL 400U/mL T4 DNA ligase (New England Biolabs #M0202)) was added and incubated at room temperature while slowly rotating end-over-end for 4 hours.

DNA extraction was performed by centrifuging the nuclei at 2,500 x g for 5 min at 4° C, discarding the supernatant, and resuspending the pellet in 300 μL of extraction buffer (10 mM Tris-HCl pH 8.0, 0.5 M NaCl, 1% SDS). For the undigested and digested control samples, 275 μL of extraction buffer was added to each control sample. 50 μL of 800 U/mL proteinase K was added and samples were incubated at 55° C for 30 min with constant mixing at 750 rpm on a ThermoMixer. The temperature was increased to 68° C and constant mixing was maintained while incubating overnight to reverse the cross-links. Samples were cooled to room temperature and briefly centrifuged. 1 μL of 20 mg/mL glycogen followed by 875 μL of 100% ice-cold ethanol was added to each sample, mixed by inverting the tube, and incubated at 80° C for 20 min. Samples were centrifuged at max speed for 30 min at 4° C. The pellet was washed twice with 800 μL of 70% ethanol and centrifuging at max speed for 10 min at 4° C. Samples were air-dried on ice for < 5 min, and, before completely drying, resuspended in 80 μL of 10 mM Tris-HCl pH 8.0. Samples were incubated at 42° C for 15 min mixing constantly at 750 rpm on a ThermoMixer to completely dissolve the DNA. 1 μL of RNase A (Thermo Scientific #EN0531) was added and incubated at 37° C for 30 min while constantly mixing at 750 rpm on a ThermoMixer. DNA concentration was determined using a a DeNovix High Sensitivity dsDNA Assay (DeNovix #KIT-DSDNA-HIGH-2) and DeNovix DS-11 FX+ fluorometer.

No more than 10 μg of DNA per Covaris microTUBE AFA Fiber (Covaris #520045) was sheared to a target peak of 400 bp using a Covaris M220 focused-ultrasonicator (Covaris, Woburn, MA) at duty factor 10%, peak incident power 50 W, 200 cycles per burst for 70 s. 125 μL of sample was transferred to a fresh 1.5 mL low-bind tube, the microTUBE washed with 75 μL of 10mM Tris-HCl pH 8.0, 1 mM EDTA and added to the sample, and the sample size selected by first adding 130 μL (0.65x) of CleanNGS DNA & RNA Clean-Up Magnetic Beads (Bulldog Bio #CNGS-0050). Sample was vortexed to mix, incubated for 5 min at room temperature, and then the beads were collected in a magnetic stand for 5 min. The supernatant was transferred to a new tube, 30 μL (0.8x final) of SPRIselect beads were added, vortexed to mix, and incubated for 5 min at room temperature. Beads were collected with a magnet and washed twice with 200 μL 85% ethanol for 30 s. Beads were air-dried on the magnet, resuspended in 53 μL of 10 mM Tris-HCl pH 8.0, and then incubated at room temperature for 5 min to elute the DNA. Beads were collected with a magnet and 51 μL of the eluate was transferred to a new tube. DNA concentration was determined using a DeNovix High Sensitivity dsDNA Assay and DeNovix DS-11 FX+ fluorometer.

50 μL of Dynabeads MyOne Streptavidin C1 beads (Life Technologies #65001) were washed twice with 100 μL 1x B&W buffer + 0.05% Tween 20 (2x Bind & Wash (B&W) buffer: 10mM Tris-HCl pH 7.5, 2M NaCl, 1mM EDTA) and then resuspended in 50 μL of 2x B&W buffer. DNA was added to the beads and incubated at room temperature for 30 min while rotating end-over-end. Beads were washed twice with 100 μL 1x B&W buffer + 0.05% Tween 20, washed twice with 100 μL of TE buffer, and resuspended in 25 μL of 10 mM Tris-HCl pH 8.0. DNA was prepared for high-throughput sequencing following the directions for ‘‘NEBNext End Prep’’ and ‘‘Adaptor Ligation’’ in the NEBNext Ultra II DNA Library Prep Kit for Illumina (New England Biolabs #E7645) while still bound to the beads. Beads were then collected with a magnet, the supernatant discarded, the beads washed twice with 100 μL 1x B&W buffer + 0.05% Tween 20, and then twice with 100 μL 10mM Tris-HCl pH 8.0, 1mM EDTA. The sample was eluted by resuspending the beads in 5 μL of 95% freshly deionized formamide, 10 mM EDTA pH 8.0, and incubating at 90° C for 10 min. Beads were collected with a magnet and the supernatant was transferred to a fresh tube containing 16 μL of 10 mM Tris-HCl pH 8.0.

Small-scale qPCR was utilized to determine the optimal number of PCR cycles for library amplification. Reactions were performed in triplicate with 4 μL 100x diluted library DNA, 6 μL primer mix (3.33 mM Universal PCR primer, 3.33 mM Index Primer, 1.66 mM MgCl_2_, 3.33x SYBR Green I), 10 μL NEBNext Ultra II Q5 Master Mix and thermocycled at 98° C for 30 s followed by 35 cycles of 98° C for 10 s, 65° C for 75 s. Linear Rn versus cycle number was plotted to determine the cycle number corresponding to one-third of maximum fluorescent intensity, and, after accounting for 100-fold dilution, the optimal number of PCR cycles needed was determined. Final library amplification was performed in duplicate using 2.5 μL water, 2.5 μL 10 mM MgCl_2_, 10 µL of premixed Index/Universal Primer from the NEBNext Multiplex Oligos for Illumina (96 Unique Dual Index Primer Pairs), 10 μL eluted DNA (undiluted), 25 μL NEBNext Ultra II Q5 Master Mix. The split sample was run on a thermocycler at 98° C for 30 s followed by X cycles (as determined above by qPCR) of 98° C for 10 s, 65° C for 75 s, followed by 1 cycle at 65° C for 5 min, and finally held at 4° C. The amplified library was purified twice as in ‘‘Cleanup of PCR Amplification’’ in the NEBNext Ultra II DNA Library Prep Kit for Illumina, using CleanNGS DNA & RNA Clean-Up Magnetic Beads volumes of 0.7x and 0.8x. The DNA was eluted in 16 μL 10mM Tris-HCl pH 8.0 and 15 μL was collected as the final product.

Final sample concentration and DNA integrity were determined using an Agilent 4200 TapeStation D1000 ScreenTape (#5067-5582). DNA was paired-end sequenced, 150 cycles each read, on a NovaSeq X Plus instrument.

Hi-C was performed in duplicate for uninduced and induced MCF10A cells, where each biological replicate represents cells from a different passage as well as treated and cross-linked independently. Two technical replicates, where each technical replicate represents cross-linked cell pellets from the same biological replicate, were utilized from each biological replicate for each experimental condition. Hi-C was performed in duplicate for non-target control sgRNA (sgNT) and NSD3-targeting guide RNA (sgNSD3) transduced TC-797 and 10-15 cells, where each biological replicate represents cells from a different passage as well as transduced and cross-linked independently.

### RNA-seq library preparation

Total RNA was extracted using the Direct-zol RNA Microprep Kit (Zymo Research #R2060) following the manufacturer’s instructions. Mammalian cells were detached from the tissue culture plates and harvested at room temperature. 500,000 cells were centrifuged at 300 x g for 5 min at room temperature and lysed in 300 μL TRI Reagent (Zymo Research #R2050-1-200) by pipetting. Lysates were mixed with 300 μL of 100% ethanol and loaded onto Zymo-Spin IC Columns (Zymo Research #C1004). Columns were washed with 400 μL of RNA Wash Buffer (Zymo Research #R1003-3-48) and treated with a total volume of 40 μL 30 U DNase I (Zymo Research #E1011-A) in DNA Digestion Buffer (Zymo Research #E1010-1-4) for 15 min at room temperature. After two washes with 400 μL of Direct-zol RNA PreWash (Zymo Research #R2050-2-160) and a final wash with 700 μL of RNA Wash Buffer, RNA was eluted in 45 μL DNase/RNase-Free Water (Zymo Research #W1001-6). RNA was aliquoted into RNase-free PCR tube strips to minimize freeze-thaw cycles and stored at -80°C freezer.

RNA integrity and concentration were assessed using the Agilent 4200 TapeStation with RNA ScreenTape (Agilent #5067-5576), RNA ScreenTape Sample Buffer (Agilent #5067-5577), and RNA ScreenTape Ladder (Agilent #5067-5578) according to the manufacturer’s instructions. Samples were denatured at 72°C for 3 min, cooled on ice, and analyzed immediately. RNA integrity number equivalent (RINe) values were obtained using the TapeStation Analysis software.

2 μL of 1:100 diluted ERCC RNA Spike-In Mix (Invitrogen #4456740) was added to each sample, which contains the cell numbers that contribute to 1 μg of RNA calculated from biological triplicates of the control samples. The RNA from an equal number of cells of each sample was prepared for high-throughput sequencing following the ribosomal RNA-depleted library preparation using KAPA RNA HyperPrep Kit with RiboErase (HMR) (Roche # 08098140702) conducted by Admera Health, LLC (South Plainfield, NJ).

### Mass spectrometry proteomics data processing

Proteomics data of counts of recovered proteins were downloaded from Dataset S1 (pnas.1702086114.sd01.xlsx) of Alekseyenko et al., 2017^42^. A pseudocount of 1 was added to each protein and then the counts from the immunoprecipitation sample were divided by the counts from the respective input sample to give relative enrichment values. Relative enrichment values were plotted as double logarithmic (log-log) plots.

### Immunofluorescence data processing

#### Colocalization analysis

Mander’s coefficients were calculated using CellProfiler^63^ v4.2.8. Two-dimensional Extended Depth of Focus (EDF) images with three channels (H3K27ac/BRD4-NUT/DAPI or H3K27ac/NSD3/DAPI), converted from three-dimensional projection confocal images by Nikon NIS-Elements Advanced Research software, were used for analysis. Illumination was corrected for images from each channel, images aligned, and image segmented to identify nuclei based on DAPI staining. The channels for co-immunofluorescence (histone H3K27ac with BRD4-NUT or histone H3K27ac with BRD4-NUT) were analyzed per identified nuclei. Mander’s coefficients were calculated by analyzing each pixel within a nucleus.

#### Condensate annotation and quantification analysis

Condensates were annotated and quantified using CellProfiler^63^ v4.2.8. Two-dimensional Extended Depth of Focus (EDF) images with three channels (H3K27ac/BRD4-NUT/DAPI or H3K27ac/NSD3/DAPI), converted from three-dimensional projection confocal images by Nikon NIS-Elements Advanced Research software, were used for analysis. Nuclei were identified based on the DAPI staining using the “IdentifyPrimaryObjects” module. The total nuclear fluorescence intensity for each channel was then quantified within the identified nuclei using the “MeasureObjectIntensity” module.

BRD4-NUT or H3K27ac were used as a marker to annotate condensates. To identify condensates, the fluorescence signal of BRD4-NUT or H3K27ac was first enhanced to highlight speckle-like features using the “EnhanceorSurpressFeature” module. Segmentation was then performed with appropriate thresholding and smoothing filters via the “IdentifyPrimaryObjects” module. For each marker, the number of annotated condensates per nucleus was calculated using the “MeasureObjectSizeShape” module. The mean fluorescence intensity of the annotated condensates and the total fluorescence intensity of the annotated condensates were calculated using the “MeasureObjectIntensity” module and extracted from *MeanIntensity* and *IntegratedIntensity* measurements, respectively. Statistical significance was determined using a one-sided Mann-Whitney *U* test.

#### Three-dimensional nuclear volume quantification

Nuclear volume was quantified in three dimensions using CellProfiler^63^ v4.2.8. The three-dimensional projection (z-stack) images with the DAPI channel were used for analysis. All the images were rescaled to a uniform intensity range and downsampled to improve processing speed. The DAPI signal was homogenized within the nuclei, and an appropriate threshold was applied for segmentation, followed by the removal of small holes caused by segmentation artifacts. The Watershed algorithm was then used to separate adjacent nuclei. Three-dimensional nuclear volume was calculated using the “MeasureObjectSizeShape” module. Statistical significance was determined using a one-sided Mann-Whitney *U* test.

### CUT&RUN data processing

#### CUT&RUN profile generation

Reads were trimmed for adaptor sequences and low-quality bases using trimmomatic v0.36. Quality control of the sequencing data was conducted using FastQC v0.11.9, both before and after trimming. FastQ Screen v0.14.0 was used to detect any potential contaminant sequences. Subsequently, reads were aligned to concatenated hg38/Genome Reference Consortium Human reference 38 (GRCh38) and dm6/BDGP Release 6 + ISO1 MT assemblies of the human and *Drosophila melanogaster* genomes, respectively, using bowtie2 v2.3.5 with parameters --local --very-sensitive-local --no-unal --nomixed --no-discordant phred33 -I 10 -X 700. Reads with mapping quality (MAPQ) >= 30 were retained using samtools v1.3.1. Picard MarkDuplicates v2.9.2 was utilized to remove PCR duplicates. The greenlist method was used to quantitatively calibrate CUT&RUN profiles via use of an internal reference^55^. Read counts from the curated set of hg38 greenlist regions^55^ were extracted using deepTools^90^ multiBamSummary v3.4.3 and used to derive a per-sample greenlist scaling factor, gamma, calculated as the ratio of reads counts from the canonical chromosomes (in thousands) to the total greenlist read counts. Normalized coverage tracks were generated using deepTools^90^ bamCoverage v3.4.3 using parameters --binSize 1 --normalizeUsing RPGC --effectiveGenomeSize 2800000000 --extendReads --samFlagInclude 64 --scaleFactor gamma.

#### Megadomain annotation

Broadly enriched regions of H3K27ac and BRD4-NUT were identified using epic2^91^ v0.0.52 on the canonical human chromosomes using the corresponding IgG sample as control with options --bin-size 200 --gaps-allowed 3. To capture the entire domain, regions within 5 kb and 15 kb of one another were combined using bedtools v2.30.0 merge for H3K27ac and BRD4-NUT, respectively. Domains were then filtered based on domain size and signal strength using the ROSE (Ranked Ordering of Super-enhancers) algorithm^92^. Domains that passed both size and signal cutoffs were retained using bedtools intersect. The above was performed on each biological replicate of H3K27ac and BRD4-NUT. To identify H3K27ac domains dependent upon BRD4-NUT, overlapping H3K27ac and BRD4-NUT domains were identified using bedtools intersect. Domains within 100 kb of one another were then combined using bedtools merge. To create a high-confidence domain list, domains identified in both replicates were intersected using bedtools intersect. These resulting domains were considered “megadomains”.

#### H3K27ac-BRD4-NUT peaks outside megadomains annotation

H3K27ac and BRD4-NUT peaks were first merged across biological replicates using bedtools intersect for each feature separately to obtain consensus peaks. Peaks located outside megadomains were then identified using bedtools intersect -v option. Overlapping H3K27ac and BRD4-NUT peaks outside megadomains were identified using bedtools intersect to identify regions co-occupied by both marks.

#### NSD3short peak annotation

Enriched regions of NSD3short were identified based on the HiBiT epitope tag from MCF10A cells induced to express NSD3short (NSD3short induced). These regions were identified using epic2 v0.0.52 on the canonical human chromosomes using the corresponding IgG as control with options --bin-size 200 --gaps-allowed 3. To capture the entire region, peaks within 5 kb distance of one another were stitched together using bedtools v2.30.0 merge. These stitched regions were then filtered based on size and signal strength. Because the Hi-C resolution was 25 kb, a cutoff size of 12.5 kb was used based on Nyquist-Shannon sampling principle; the signal cutoff was determined using the ROSE algorithm^92^. The above was repeated across all 3 replicates. To create a high-confidence peak list, regions identified in at least 2 replicates were selected using bedtools multiinter with -cluster option. To account for slight trimming of domains following intersection, the final consensus regions were again filtered using a cutoff size of 12.5 kb. These resulting domains were considered “NSD3short peaks”.

#### NSD3short^W284A^ domain annotation

Broadly enriched regions of NSD3 and HiBiT epitope tag from MCF10A cells induced to express NSD3short^W284A^ (NSD3short^W284A^ induced) were identified using epic2 v0.0.52 on the canonical human chromosomes using IgG sample as control with options --bin-size 200 and --gaps-allowed 3. To capture the broad domain region, peaks within 30 kb distance of one another were stitched together using bedtools v2.30.0 merge. These domains were then filtered based on domain size and signal strength using the ROSE algorithm^4^. Domains that passed both size and signal cutoffs were retained using bedtools intersect. Common domains identified for NSD3 and HiBiT epitope tag were obtained using bedtools intersect, retaining the original coordinates from NSD3. The domains within 100 kb of one another were combined using bedtools merge. The above was repeated across all 3 replicates. To create a high-confidence domain list, domains identified in at least 2 replicates were selected using bedtools multiinter with default parameters. In cases where this resulted in consecutive domains, such domains were merged using bedtools merge -d 0. These resulting domains were considered “NSD3short^W284A^ domains”.

#### Metaplot and heatmap visualization

To account for background signal, greenlist normalized CUT&RUN profiles were normalized using IgG. This was done using deepTools^90^ bigwigCompare v3.4.3 with parameters --operation subtract --pseudocount 0 0 --binSize 100. Any negative values in the IgG-normalized profiles were clipped to 0, as these reflect the inherent sparsity of CUT&RUN data rather than the biological signal. All metaplots and heatmaps were generated using deepTools^90^ v3.4.3.

#### CUT&RUN enrichment quantification

CUT&RUN enrichment was quantified using deepTools^90^ v3.4.3. Read counts were first obtained with either multiBamSummary (10-15 and TC-797 cells) or multiBigwigSummary (MCF10A cells) functions, using the corresponding greenlist normalized files for each sample. Enrichment values for each sample were divided by the corresponding IgG control enrichment values to normalize for background signal. Mean enrichment values relative to IgG were then computed by averaging the normalized signal across all biological replicates. Statistical significance was determined using a one-sided Mann-Whitney *U* test.

### Hi-C data processing

#### Hi-C contact map generation

Data was processed using the Juicer pipeline^64,93^. Reads were aligned to concatenated hg38/Genome Reference Consortium Human Reference 38 (GRCh38) and dm6/BDGPRelease 6 + ISO1 MT assemblies of the human and *Drosophila melanogaster* genomes, respectively, using BWA-MEM v0.7.18-r1243-dirty with parameter -SP5M. Reads were then assigned to MboI restriction fragments, duplicates removed, reads with a MAPQ >= 30 were retained, and reads mapping to the same restriction fragment removed. The genome was divided into equally spaced, non-overlapping bins and the number of reads counted in each pair of bins using Juicer Tools^64,93^ v1.22.01 with command juicer pre. Hi-C contact maps were then converted from .hic format to .cool format using hic2cool v0.8.3.

Hi-C contact maps were normalized by matrix balancing using the iterative correction (IC) algorithm^94^ using cooler^95^ v0.9.3 with command balance and parameters --max-iters 1000 --convergence-policy store_nan. The GRCh38 and *Drosophila melanogaster* genomes were normalized separately. Replicate reproducibility was evaluated by computing the HiCRep stratum adjusted correlation coefficient (SCC)^96^ using hicreppy^97^ v0.1.0 with parameters --hvalue 20 --maxdist 5000000 at 10 kb resolution. As biological and technical replicates were highly reproducible (Table S4), datasets were merged after filtering, as described above, and the combined contact maps IC normalized. All analysis was performed on and figures represent merged, IC normalized contact maps.

#### Contact probability as a function of genomic distance

Contact probability (*P*) as a function of genomic separation (s) (*P*(*s*) curve) was calculated using cooltools^98^ v0.7.1. Expected *cis* contact probabilities were calculated using the expected_cis function from cooltools on chromosome arms omitting arms chr13_p, chr14_p, chr15_p, chr21_p, chr22_p because of their poor mappability with parameters smooth=True, aggregate_smoothed=True, smooth_sigma=0.1. *P*(*s*) curves were plotted as double logarithmic (log-log) plots.

#### Annotating megadomain-megadomain and NSD3short contacts

CUT&RUN annotated megadomains and NSD3short peaks were used to identify megadomain-megadomain contacts and NSD3short contacts, respectively, based on enrichment relative to local background as previously described^34^. All possible megadomain-megadomain contacts were enumerated genome-wide; all possible interactions between NSD3short peaks were enumerated in *cis*. Next, for each potential interaction, a donut-shaped local background was defined. The width of this donut was 3 pixels, excluding a 1-pixel gap surrounding the interaction. The average number of ICE normalized contacts within this donut was obtained. The donut signal was then adjusted by dividing the average expected contact frequency for the candidate interaction by the donut’s average expected value to normalize for the distance-dependent decay in contact frequency.

*Cis* and *trans* megadomain-megadomain contacts were analyzed and annotated separately. The candidate contacts overlapping the diagonal and regions with enrichment less than 2-fold relative to their respective donut region were removed. Statistical significance was assessed by testing whether the observed raw contact counts within each candidate interaction were significantly higher than the expected; the corresponding donut region was used as the expected. Multiple hypothesis testing was performed using the Benjamini-Hochberg FDR control procedure with an FDR threshold of 0.005 for *cis* interactions and 0.1 for *trans* interactions. This process was performed at 100 kb resolution for non-target sgRNA (sgNT) control cells to identify *cis* and *trans* megadomain-megadomain contacts.

Boundaries for NSD3short peaks were adjusted to the start and end of 25 kb Hi-C bins by rounding the peak start coordinates down and peak end coordinates up. In cases where this resulted in consecutive peak regions, such peaks were merged using bedtools merge -d 0. Interactions between NSD3short peaks were analyzed and annotated in *cis* at 25 kb resolution from NSD3short uninduced and NSD3short induced samples. The candidate contacts overlapping the diagonal and regions with enrichment less than 2-fold relative to their respective donut region were removed. An FDR threshold of 0.05 was applied. To obtain contacts specific to the NSD3short induction, contacts common to both uninduced and NSD3short induced conditions were removed from the induced set using bedtools pairToPair -type notboth. Similarly, *cis* contacts were annotated from NSD3short^W284A^ induced Hi-C contact maps at 25 kb resolution using NSD3short peaks with the same threshold of enrichment and FDR, and contacts shared with the uninduced sample were excluded to identify mutant-specific contacts.

#### Peak pixel identification of megadomain-megadomain contacts

After identifying megadomain-megadomain contacts at 100 kb resolution, we refined the interaction to 25 kb resolution by identifying the peak pixel for each interaction using connected component analysis. For each megadomain-megadomain interaction we identified all pixels for that interaction at 25 kb resolution. We then identified 4-connectivity (edges only) clusters of connected pixels (components) for each megadomain-megadomain interaction. Connected components were required to have a minimum size of 5 pixels. If multiple connected components were identified, the connected component with the greatest number of pixels was used. The pixel with the highest contact frequency was then identified as the peak pixel.

We detected 2,167 (*cis*: 160; *trans*: 2,007) and 1,350 (*cis*: 84; *trans*: 1,266) megadomain-megadomain interactions in 10-15 and TC-797 cells, respectively. 2,074 (*cis*: 160; *trans*: 1,914) and 1,243 (*cis*: 84; *trans*: 1,159) megadomain-megadomain interaction peak pixels were identified from 10-15 and TC-797 cells, respectively, due to the required minimum size of 5 connected component pixels.

#### Pileups

Pileups were computed using coolpuppy^99^ v1.1.0. Expected contact frequencies for *cis* or *trans* chromosomes were calculated using the expected_cis or expected_trans functions from cooltools^98^ v0.7.1. Pileup analysis was performed using the coolpup.pileup function from coolpuppy, using the calculated expected contact frequencies, and with parameters min_diag = 0, and mindist = 0. flank = 250_000 parameter was used for megadomain-megadomain interaction peak pixels, flank = 125_000 was used for NSD3short contacts, and flank = 250_000 was used for all possible *cis* NSD3short^W284A^ domain-domain contacts. Any region plus its flank extending beyond the chromosome boundaries was excluded from the analysis. Additionally, for pileups of megadomain-megadomain interaction peak pixels, contacts with undefined or zero observed/expected contact frequency in any dataset were excluded from pileups. 1,116 (*cis*: 150; *trans*: 966) and 649 (*cis*: 78; *trans*: 571) megadomain-megadomain interaction peak pixels were analyzed by pileups between conditions for 10-15 and TC-797 cells, respectively, due to the requirements that the region plus its flank not extend beyond chromosome boundaries and that contacts be defined and greater than zero in all datasets. These requirements did not exclude any NSD3short contacts from pileups. Pileups were also generated using randomly shifted control regions using the parameter nshifts = 100. Aggregate pileups were visualized using the plotpup.plot function. To compare pileups between conditions, pileups were divided using the divide_pups function. Pileups were calculated at 25 kb resolution for megadomain-megadomain peak pixels, 25 kb resolution for NSD3short contacts, and 100 kb resolution for all possible *cis* NSD3short^W284A^ domain-domain contacts.

#### Quantification and comparison of contacts between conditions

To quantify and compare contacts between conditions the mean observed/expected contact frequency was calculated for each candidate interaction. The expected contact frequency for *cis* and *trans* contacts was determined using the expected_cis or expected_trans functions from cooltools^98^ v0.7.1, respectively. Contacts with undefined or zero observed/expected contact frequency in any dataset were excluded from quantification and comparison. 1,375 (*cis*: 157; *trans*: 1,218) and 737 (*cis*: 78; *trans*: 659) megadomain-megadomain interaction peak pixels were quantified and compared between conditions for 10-15 and TC-797 cells, respectively, due to the requirement that contacts be defined and greater than zero in all datasets. 771 NSD3short contacts were quantified and compared between conditions due to the requirement that contacts be defined and greater than zero in all datasets. Statistical significance was assessed using either a one-sided or two-sided Mann-Whitney *U* test, as appropriate.

#### CTCF-CTCF loop annotation

First, dots in Hi-C contact maps were annotated using the dots algorithm from cooltools^98^ v0.7.1. Second, dots were annotated with the CTCF motif and those dots with convergent CTCF motifs were identified as CTCF-CTCF loops.

#### Compartment annotation

To annotate compartments, eigenvector decomposition was performed using the eigs_cis function from cooltools^98^ v0.7.1 at 100 kb resolution using GC content to determine the orientation of the eigenvectors. The first eigenvector (EV1) was used as the compartment signal. Regions with positive EV1 values were assigned to the A compartment, while those with negative EV1 values were assigned to the B compartment.

#### Compartment analysis for NSD3short contacts

Compartment A and compartment B BED files obtained using NSD3short uninduced samples were converted into a BEDPE format by enumerating all possible intra-chromosomal contacts genome-wide, including self-interacting contacts to capture compartments near the diagonal. Unique NSD3short contacts obtained from NSD3short induced samples were overlapped with AA and BB BEDPE files using bedtools pairToPair -type both. Contacts assigned to AA and BB compartments were concatenated and subsequently overlapped with NSD3short contacts using bedtools pairToPair -type notboth to identify contacts spanning both compartments (AB).

To compute statistical significance, each NSD3short interaction in the BEDPE file was first assigned a unique ID and then split into individual contact sites. Contact sites were shuffled independently using bedtools shuffle with parameters -chrom and -excl, ensuring that the shuffled anchors were retained on their original chromosome and were excluded from the centromeric regions and genome assembly gaps. After shuffling, anchors were re-paired based on their IDs to reconstruct the shuffled BEDPE file, generating randomly shuffled control contacts. Overlap counts for each shuffled interaction in the AA, BB, and AB compartments were calculated. This was repeated 10,000 times to generate a null distribution of the expected overlap counts. A random permutation test was then used to calculate a p value by comparing the observed overlap count to the null distribution of expected counts.

#### Compartment analysis for NSD3short^W284A^ domains

NSD3short^W284A^ domains were intersected with the compartment A and compartment B BED files. Regions corresponding to gaps within the compartment BED files were identified using bedtools complement, and any domain overlapping these gaps by >= 50% of its length was excluded from further analysis. In case the domain overlapped with both compartments, assignment was based on the compartment with >= 50% overlap and a greater overlap fraction compared to the other compartment.

To compute statistical significance, a control set was generated by shuffling the filtered domains using bedtools shuffle with parameters -chrom -noOverlapping -excl. These options ensured that the shuffled domains remained on their original chromosome, did not overlap with each other, and were excluded from compartment gaps, centromeric regions, and genome assembly gaps, respectively. Each shuffled set was then intersected with the compartment BED files using the same criteria as above, and counts were computed. This was repeated 10,000 times to generate a null distribution of expected overlap counts. A random permutation test was then used to calculate a p value by comparing the observed overlap count to the null distribution of expected counts. The above was performed using compartments annotated from both uninduced and NSD3short^W284A^ samples.

### RNA-seq data processing

#### Differential Gene Expression Analysis

Raw reads were assessed for quality using FastQC v0.12.1 and FastQ Screen v0.14.0. Reads were trimmed for adaptor sequences and low-quality bases using trimmomatic v0.39. Cleaned reads were then aligned to the concatenated reference containing the GRCh38 genome assembly with GENCODE v29 gene annotations and ERCC RNA spike-in controls using STAR^100^ v2.7.11a with parameter --outSAMtype BAM SortedByCoordinate. Samtools v1.3.1 was used to index the BAM files with samtools index.

Gene-level counts were obtained using featureCounts^101^ v2.0.6 with options -p --countReadPairs -s 2. Reads were counted against GENCODE v29 gene annotations and ERCC RNA spike-in controls. Count data was then imported into R v4.2.3 for normalization and downstream analysis. To account for unwanted variation, normalization was performed using RUVSeq^102^ v1.36.0 with the RUVg method, which utilizes the ERCC spike-in controls as negative control genes. Genes with fewer than 5 counts in at least 2 samples were excluded from RUVg normalization, and the number of unwanted factors (k) was set as 1.

Differential expression analysis was then performed using DESeq2^103^ v1.42.0. Genes with fewer than 10 counts in all samples for NSD3-depletion (sgNSD3) versus non-targeting sgRNA (sgNT) comparison and genes with fewer than 10 counts in at least 3 samples for MCF10A NSD3short induced vs NSD3short uninduced and NSD3short^W284A^ induced vs NSD3short^W284A^ uninduced comparisons were excluded from the analysis. Differentially expressed genes (DEGs) were identified using an absolute log_2_ fold-change threshold of 0.585 (fold-change of 1.5) and Benjamini-Hochberg adjusted p value of 0.05 for sgNSD3 vs sgNT comparison to capture as many DEGs as possible. An absolute log_2_ fold-change threshold of 1 (fold-change of 2) and Benjamini-Hochberg adjusted p value of 0.01 for NSD3short induced vs NSD3short uninduced and NSD3short^W284A^ induced vs NSD3short^W284A^ uninduced comparisons to more conservatively filter the data. Functional enrichment analysis of DEGs was performed using clusterProfiler^104^ v4.10.0.

#### RNA-seq data visualization

ggplot2 v3.5.2 was used for data visualization. enrichplot (DOI: 10.18129/B9.bioc.enrichplot) v1.22.0 was used for generating the enrichment plot for Gene Set Enrichment Analysis (GSEA).

#### Megadomain gene annotation

Promoter regions were defined using GENCODE v29 gene annotations. For each gene, the promoter was set as 1,000 bp upstream and 100 bp downstream of the transcription start site. Then, megadomains identified from CUT&RUN were intersected with promoters using bedtools intersect to generate a list of genes whose promoters overlapped megadomains. These resulting genes were considered “megadomain genes”.

## SUPPLEMENTARY FIGURES

**Figure S1.**
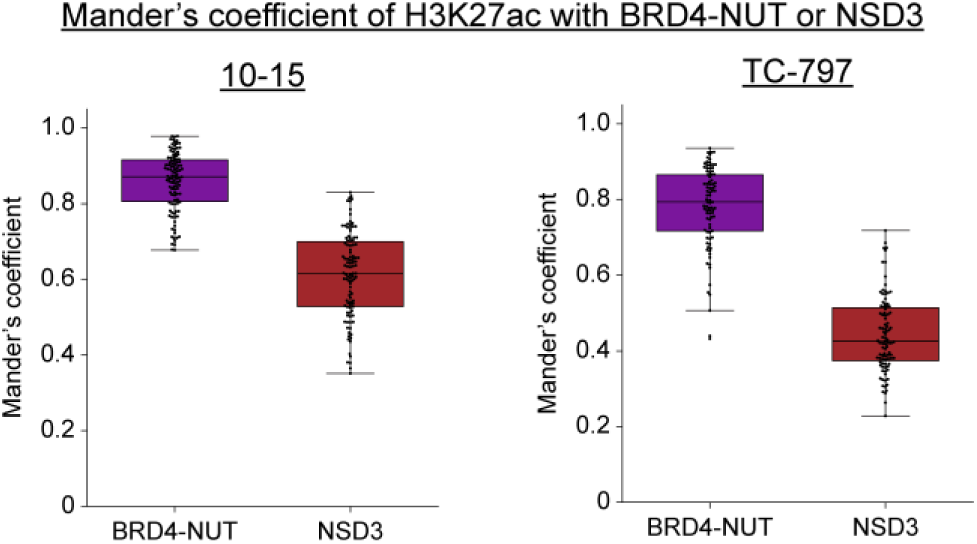
Further analysis of factors overlapping with BRD4-NUT nuclear condensates. Box plots (horizontal lines indicate medians; boxes indicate interquartile range; whiskers extend 1.5 times the interquartile range) of Mander’s coefficients of histone H3K27ac with BRD4-NUT and histone H3K27ac with NSD3. Data points represent individual nuclei. 110 (10-15 cells) and 89 (TC-797 cells) nuclei were used to analyze the overlap of histone H3K27ac with BRD4-NUT. 106 (10-15 cell) and 95 (TC-797 cell) nuclei were used to analyze the overlap of histone H3K27ac with NSD3.

**Figure S2.**
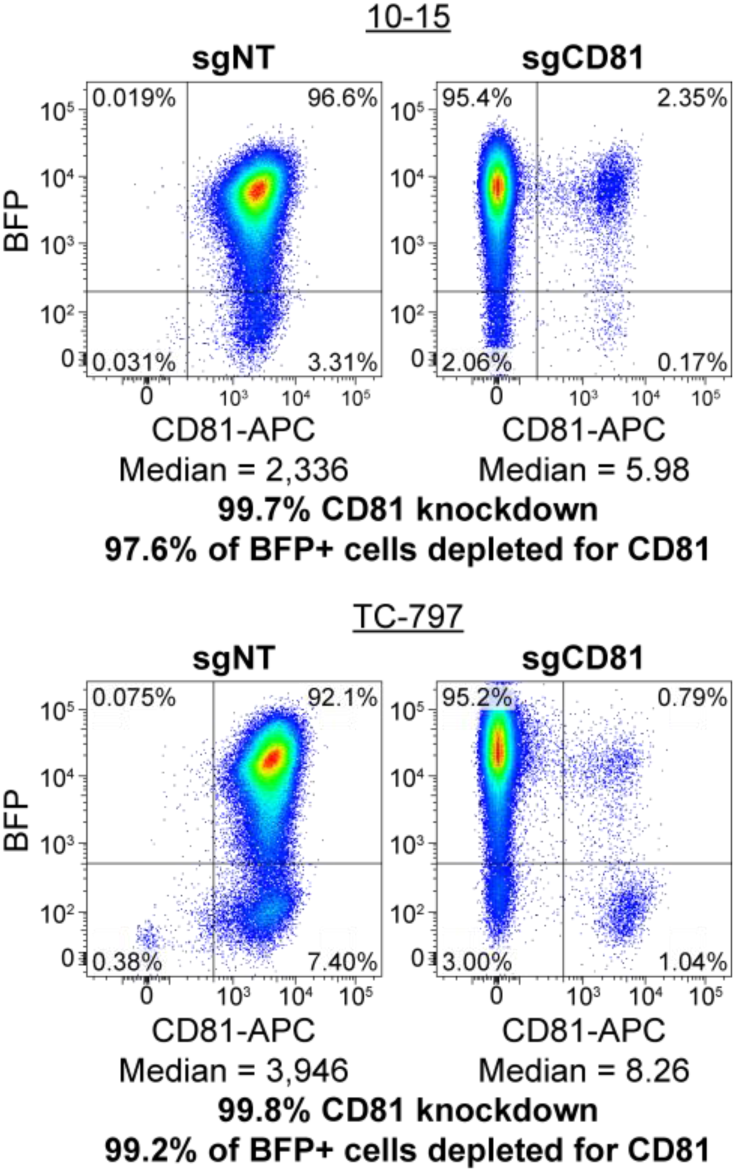
Validation of 10-15 and TC-797 cell lines expressing the Zim3-dCas9 CRISPRi effector. NUT carcinoma cell lines 10-15 and TC-797 stably expressing the CRISPRi Zim3-dCas9 CRISPRi transcriptional repressor were validated by quantifying depletion of the non-essential cell surface protein CD81. 10-15 and TC-797 cells were transduced with a non-targeting sgRNA (sgNT) or a sgRNA targeting CD81 (sgCD81). Knockdown was assessed by flow cytometry. Knockdown efficiency was 99+% with 97+% of the cell population depleted of CD81. sgRNA lentiviral vectors contain a BFP marker.

**Figure S3.**
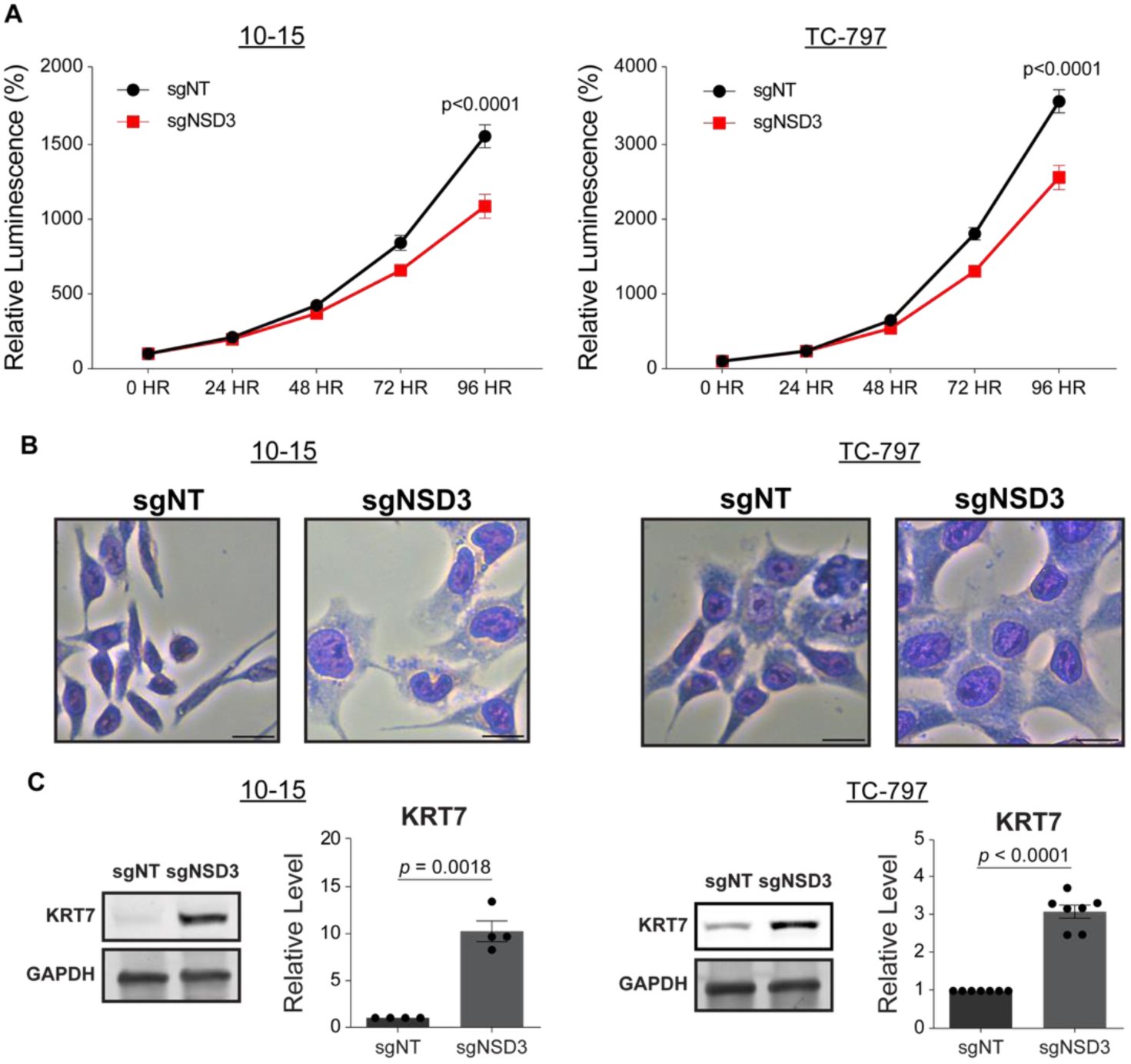
Depletion of NSD3 slows proliferation and promotes differentiation of NUT carcinoma cells. (**A**) Cell proliferation of control (sgNT) and NSD3-depleted (sgNSD3) 10-15 and TC-797 cells on each day given on the x-axis determined by CellTiter-Glo assay. Data points and error bars represent the mean and standard error of the mean (SEM), respectively, of three biological replicates. p value determined using paired t-test. (**B**) Photomicrographs of control (sgNT) and NSD3-depleted (sgNSD3) 10-15 and TC-797 cells show that the depletion of NSD3 induces differentiated cell morphology as evidenced by enlarged cells and nuclei, flattened cells, and a lower nucleus-to-cytoplasm ratio. Cells were transduced with a lentivirus expressing sgNT or sgNSD3 for 11 days, followed by Wright-Giemsa staining. Scale bars, 20 µm. (**C**) Immunoblots indicate that depletion of NSD3 leads to the upregulation of epithelial-specific differentiation marker keratin 7 (KRT7) in 10-15 and TC-797 cells. Cells were transduced with a lentivirus expressing sgNT (control) or sgNSD3 for 5 days (TC-797 cells) or 8 days (10-15 cells). GAPDH was used as an internal control. Quantification is shown to the right of each immunoblot; data points represent biological replicates; bar and error bars represent mean and standard error of the mean, respectively. p value determined using paired t-test.

**Figure S4.**
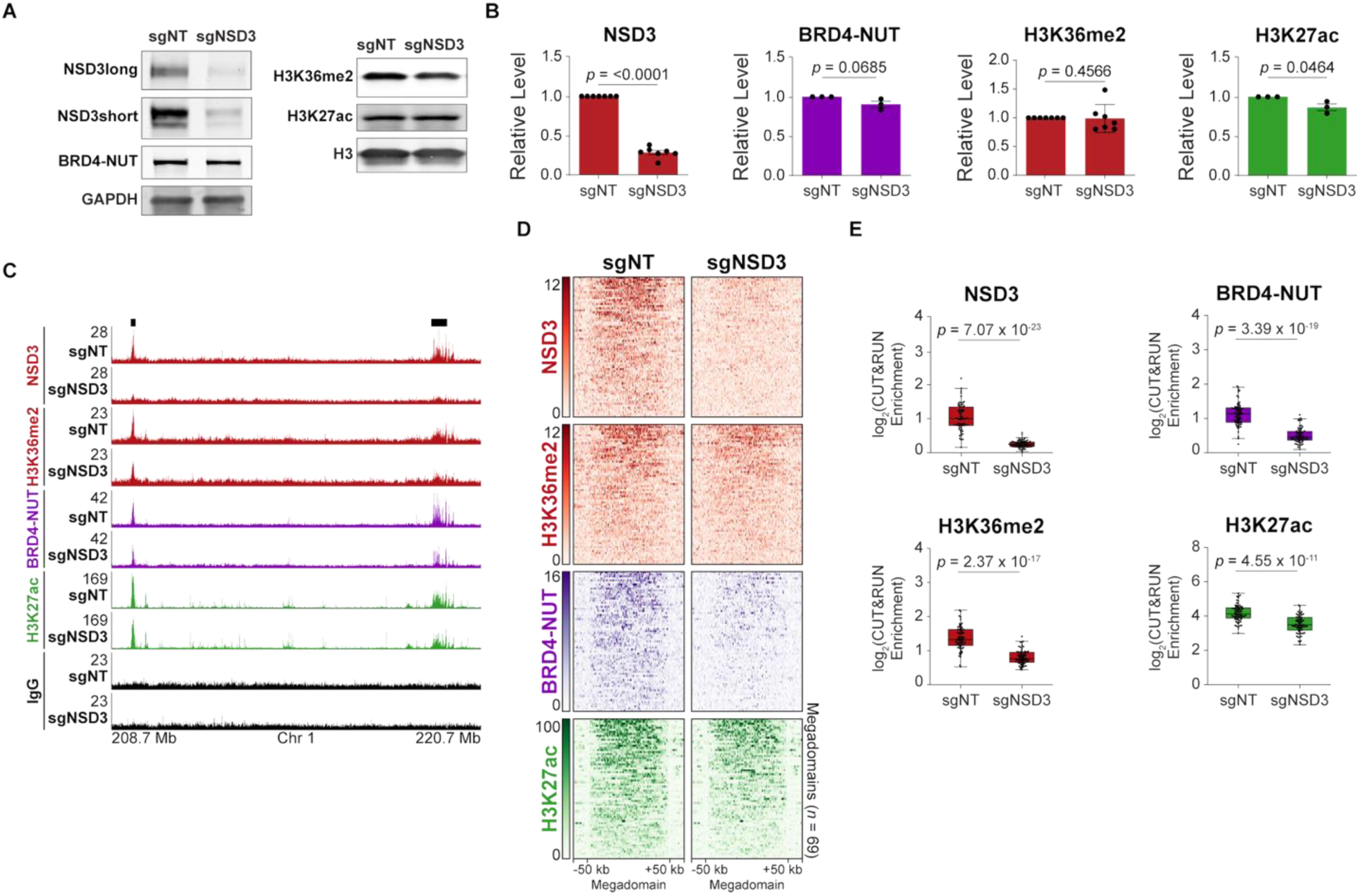
NSD3 deposits H3K36me2 in megadomains and stabilizes BRD4-NUT on chromatin in TC-797 cells. (**A**) Immunoblots indicate NSD3 (long and short isoforms) levels are reduced, but BRD4-NUT, histone H3K36me2, and histone H3K27ac levels in TC-797 cells are unchanged after CRISPRi-mediated depletion of NSD3 (sgNSD3). A non-target sgRNA (sgNT) was used as a control. GAPDH and histone H3 were used as internal controls for non-histone proteins and histone modifications, respectively. (**B**) Quantification of the immunoblots from (A). NSD3short and NSD3long were combined to obtain the level of NSD3. Data points represent biological replicates. Bar and error bars represent the mean and standard error of the mean, respectively, of biological replicates. p value determined using paired t-tests. (**C**) CUT&RUN profiles for NSD3 (red), histone H3K36me2 (red), BRD4-NUT (purple), histone H3K27ac (green), and IgG (black) from control (sgNT) TC-797 cells and NSD3-depleted (sgNSD3) TC-797 cells. Chromatin occupancy of histone H3K36me2 and BRD4-NUT within megadomains (black bars) is reduced upon depletion of NSD3. (**D**) Heatmaps of the CUT&RUN signal for NSD3, histone H3K36me2, BRD4-NUT, and histone H3K27ac within each megadomains in control (sgNT) and NSD3-depleted (sgNSD3) TC-797 cells. Megadomains were normalized to the same length, and 50 kb of flanking DNA is shown next to each normalized megadomain. IgG signal has been subtracted from each heatmap. (**E**) Box plots (horizontal lines indicate medians; boxes indicate interquartile range; whiskers extend 1.5 times the interquartile range) indicate that NSD3, histone H3K36me2, BRD4-NUT, and histone H3K27ac are reduced within megadomains in NSD3-depleted (sgNSD3) compared to control (sgNT) TC-797 cells. Data points represent the average IgG-normalized CUT&RUN signals within megadomains from two biological replicates. p values determined using one-sided Mann-Whitney *U* tests.

**Figure S5.**
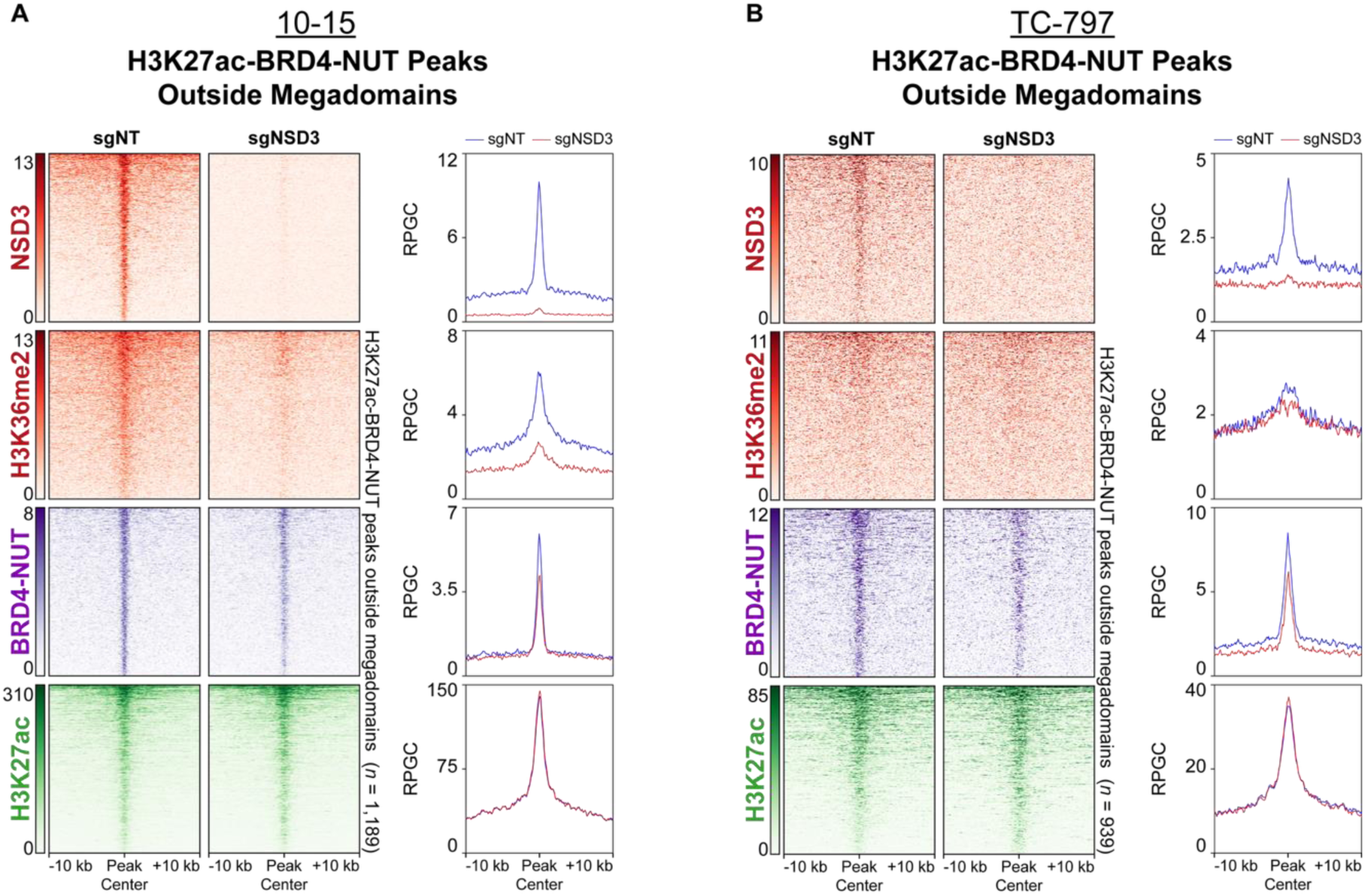
CUT&RUN analysis of histone H3K27ac-BRD4-NUT peaks outside of megadomains upon CRISPRi-mediated depletion of NSD3. (**A**) (Left) Heatmaps of the CUT&RUN signal for NSD3, histone H3K36me2, BRD4-NUT, and histone H3K27ac centered at histone H3K27ac peaks overlapping BRD4-NUT peaks (H3K27ac-BRD4-NUT peaks) outside of megadomains in control (sgNT) and NSD3-depleted (sgNSD3) 10-15 cells. H3K27ac-BRD4-NUT peak centers outside of megadomains were aligned, and 10 kb of DNA flanking each peak is shown. IgG signal has been subtracted from each heatmap. (Right) Profiles of the mean CUT&RUN signal for NSD3, histone H3K36me2, BRD4-NUT, and histone H3K27ac centered at H3K27ac-BRD4-NUT peaks outside of megadomains in control (sgNT, blue) and NSD3-depleted (sgNSD3, red) 10-15 cells with 10 kb flanking DNA. IgG signal has been subtracted from each profile. (**B**) Same as (A) for TC-797 cells.

**Figure S6.**
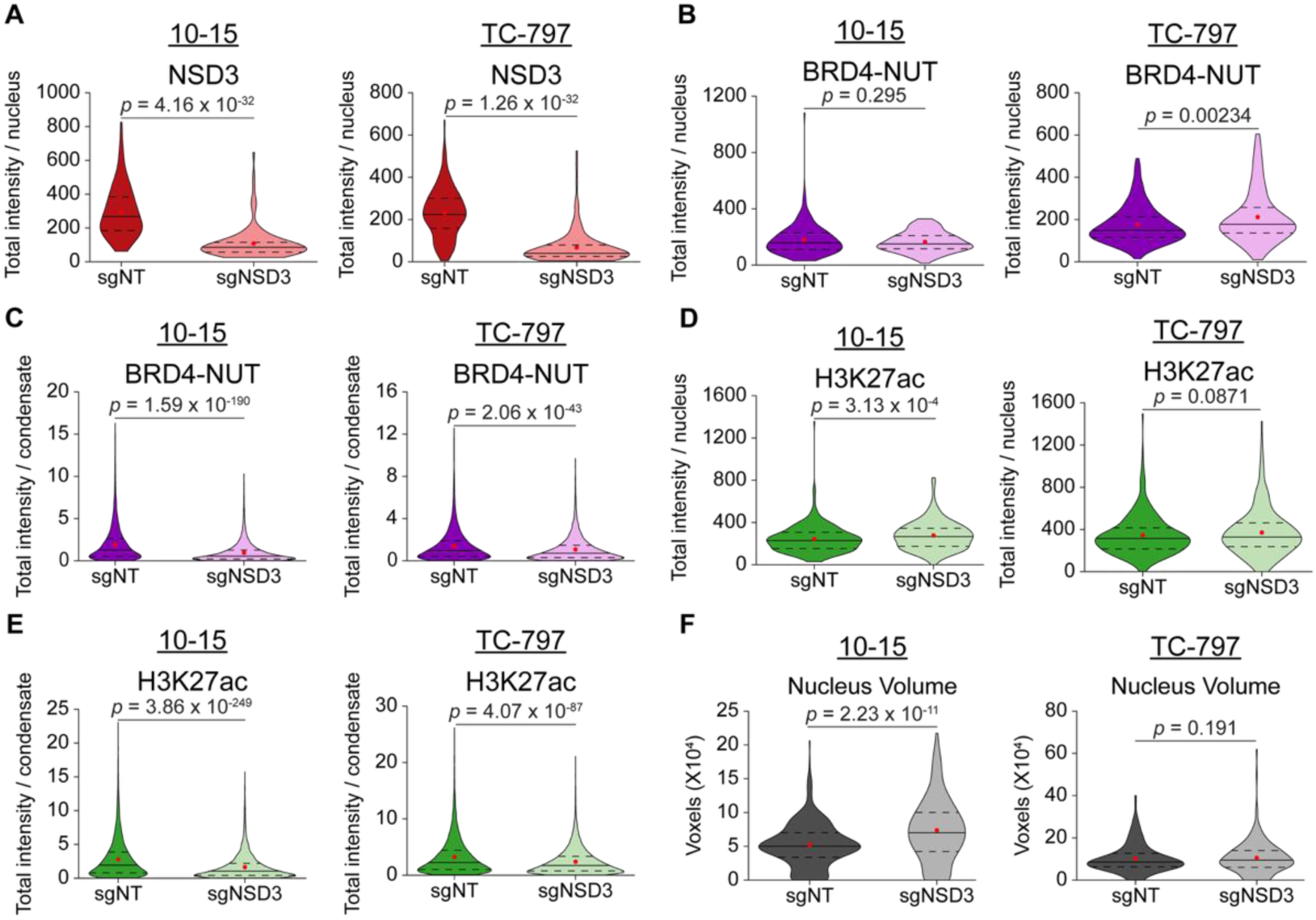
Further analysis of BRD4-NUT condensates upon depletion of NSD3. (**A**) Violin plots (solid lines indicate medians; red dots indicate means; dashed lines indicate 25th and 75th percentiles) indicate that depletion of NSD3 significantly decreases the total nuclear fluorescence intensity of NSD3. 161 (sgNT, dark red) and 124 (sgNSD3, light red) 10-15 cell nuclei were analyzed; 150 (sgNT, dark red) and 141 (sgNSD3, light red) TC-797 cell nuclei were analyzed. p values determined using one-sided Mann-Whitney *U* tests. (**B**) Violin plots (solid lines indicate medians; red dots indicate means; dashed lines indicate 25th and 75th percentiles) of the total nuclear fluorescence intensity of BRD4-NUT. 216 (sgNT, purple) and 123 (sgNSD3, pink) 10-15 cell nuclei were analyzed; 164 (sgNT, purple) and 168 (sgNSD3, pink) TC-797 cell nuclei were analyzed. NSD3 depletion slightly increases the total nuclear fluorescence intensity of BRD4-NUT in TC-797 cells but does not affect the intensity of BRD4-NUT in 10-15 cells. p values determined using one-sided Mann-Whitney *U* tests. (**C**) Violin plots (solid lines indicate medians; red dots indicate means; dashed lines indicate 25th and 75th percentiles) indicate that depletion of NSD3 significantly reduces the total fluorescence intensity of BRD4-NUT condensates. 5,710 (sgNT, purple) and 4,318 (sgNSD3, pink) condensates were analyzed from 10-15 cell nuclei; 5,594 (sgNT, purple) and 7,511 (sgNSD3, pink) condensates were analyzed from TC-797 cell nuclei. p values determined using one-sided Mann-Whitney *U* tests. (**D**) Violin plots (solid lines indicate medians; red dots indicate means; dashed lines indicate 25th and 75th percentiles) of the total nuclear fluorescence intensity of histone H3K27ac. 377 (sgNT, dark green) and 247 (sgNSD3, light green) 10-15 cell nuclei were analyzed; 314 (sgNT, dark green) and 309 (sgNSD3, light green) TC-797 cell nuclei were analyzed. NSD3 depletion slightly increases the total nuclear fluorescence intensity of histone H3K27ac in 10-15 cells but does not affect the intensity of histone H3K27ac in TC-797 cells. p values determined using one-sided Mann-Whitney *U* tests. (**E**) Violin plots (solid lines indicate medians; red dots indicate means; dashed lines indicate 25th and 75th percentiles) indicate that depletion of NSD3 significantly reduces the total fluorescence intensity of histone H3K27ac in condensates. 10,064 (sgNT, dark green) and 9,151 (sgNSD3, light green) condensates were analyzed from 10-15 cell nuclei; 10,691 (sgNT, dark green) and 13,408 (sgNSD3, light green) condensates were analyzed from TC-797 cell nuclei. p values determined using one-sided Mann-Whitney *U* tests. (**F**) Violin plots (solid lines indicate medians; red dots indicate means; dashed lines indicate 25th and 75th percentiles) of nuclear volume identified by DAPI staining. 455 (sgNT, dark gray) and 273 (sgNSD3, light gray) 10-15 cell nuclei were analyzed; 268 (sgNT, dark gray) and 303 (sgNSD3, light gray) TC-797 cell nuclei were analyzed. NSD3 depletion significantly increases the nuclear volume of 10-15 cells but does not affect the nuclear volume of TC-797 cells. p values determined using one-sided Mann-Whitney *U* tests.

**Figure S7.**
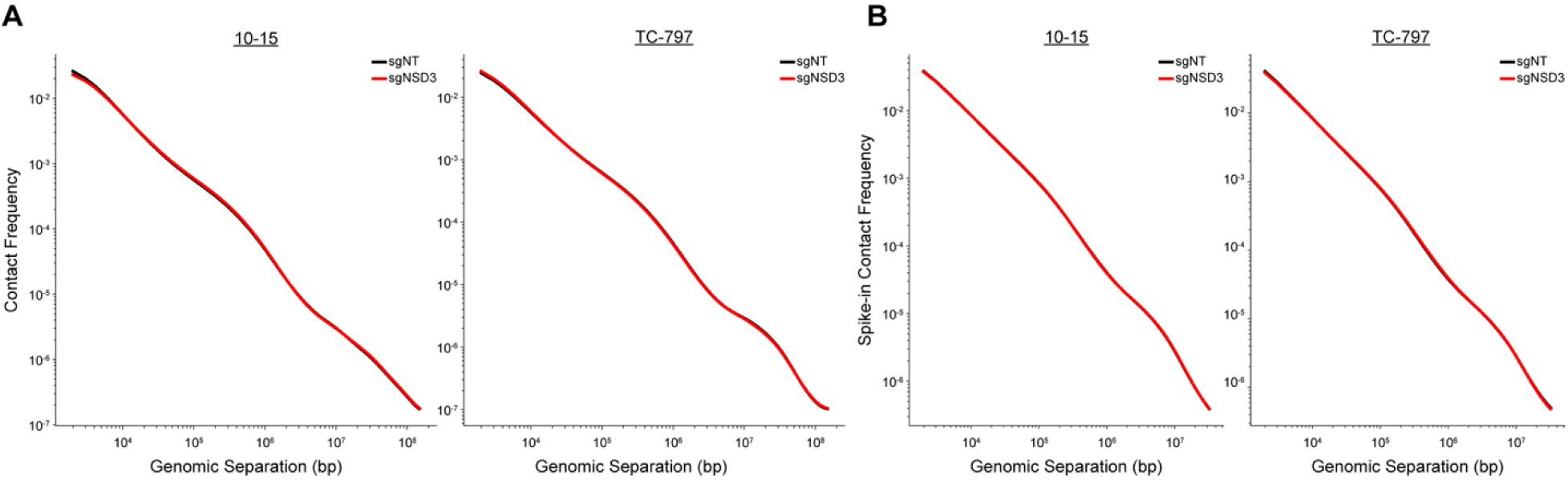
Further analysis of Hi-C data from NUT carcinoma cells depleted of NSD3. (**A**) *P*(*s*) curves of the average intrachromosomal contact frequency separated by the genomic distance on the x-axis in control (sgNT, black) and NSD3-depleted (sgNSD3, red) 10-15 and TC-797 cells. (**B**) *P*(*s*) curves of the average intrachromosomal contact frequency for loci from identical amounts of Kc167 *Drosophila melanogaster* cells spiked-in to identical amounts of control (sgNT, black) and NSD3-depleted (sgNSD3, red) 10-15 and TC-797 cells separated by the genomic distance on the x-axis. Overlap of the *Drosophila P*(*s*) curves in both conditions at all genomic separations indicates no technical differences in the Hi-C analysis between control and NSD3-depleted cells.

**Figure S8.**
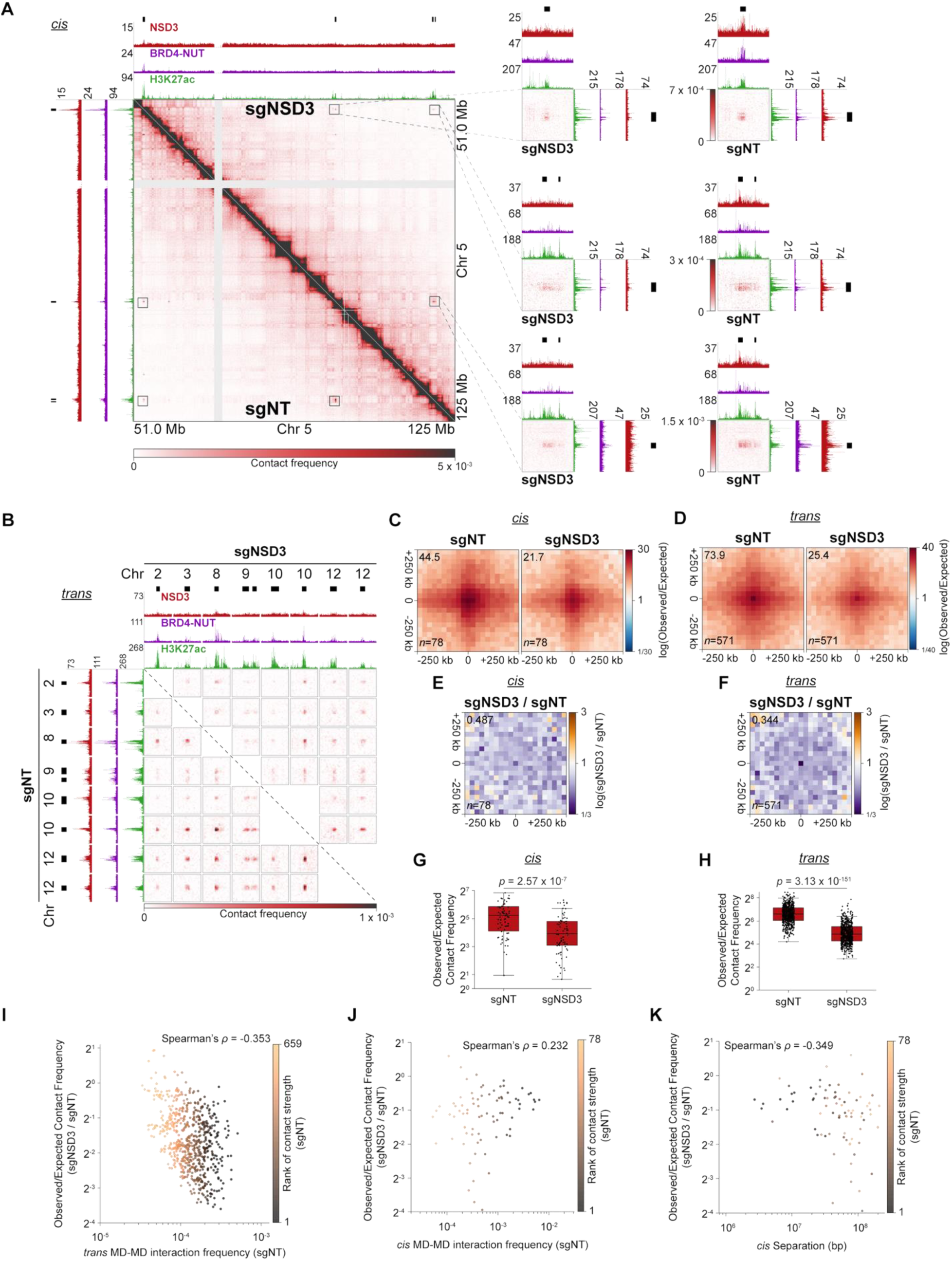
NSD3 stabilizes chromatin 3D contacts between BRD4-NUT megadomains in TC-797 cells. (**A**) Intrachromosomal (*cis*) Hi-C contact maps at 100 kb resolution (left bottom) from control (sgNT; below diagonal) and NSD3-depleted (sgNSD3; above diagonal) TC-797 cells show that the depletion of NSD3 reduces *cis* megadomain-megadomain (MD-MD) contacts. Boxed regions displaying individual MD-MD contacts are enlarged to the right and the top of the contact map at 10 kb resolution. CUT&RUN profiles for NSD3, BRD4-NUT, and histone H3K27ac are aligned with the contact maps (sgNT: vertically; sgNSD3: horizontally for the 100 kb resolution map). Black bars indicate megadomains. (**B**) Interchromosomal (*trans*) Hi-C contact maps at 100 kb resolution from control (sgNT; below diagonal) and NSD3-depleted (sgNSD3; above diagonal) TC-797 cells show that the depletion of NSD3 reduces *trans* MD-MD contacts. CUT&RUN profiles for NSD3, BRD4-NUT, and histone H3K27ac are aligned with the contact maps (sgNT: vertically; sgNSD3: horizontally for the 100 kb resolution map). Black bars indicate megadomains. (**C**) Genome-wide pileups of mean observed/expected *cis* Hi-C contact frequencies at MD-MD contacts for control (sgNT) and NSD3-depleted (sgNSD3) TC-797 cells. Values of the central pixel are given at top left. Windows extending beyond chromosome ends and contacts with zero or undefined observed/expected values in any dataset were excluded from pileups. (**D**) Genome-wide pileups of mean observed/expected *trans* Hi-C contact frequencies at MD-MD contacts for control (sgNT) and NSD3-depleted (sgNSD3) TC-797 cells. Values of the central pixel are given at top left. Windows extending beyond chromosome ends and contacts with zero or undefined observed/expected values in any dataset were excluded from pileups. (**E**) Genome-wide pileups of the fold-change in mean observed/expected *cis* Hi-C contact frequencies at MD-MD contacts for NSD3-depleted (sgNSD3) versus control (sgNT) TC-797 cells. The value of the central pixel is given at the top left. Windows extending beyond chromosome ends and contacts with zero or undefined observed/expected values in any dataset were excluded from pileups. (**F**) Genome-wide pileups of the fold-change in mean observed/expected *trans* Hi-C contact frequencies at MD-MD contacts for NSD3-depleted (sgNSD3) versus control (sgNT) TC-797 cells. The value of the central pixel is given at the top left. Windows extending beyond chromosome ends and contacts with zero or undefined observed/expected values in any dataset were excluded from pileups. (**G**) Box plots (horizontal lines show medians; boxes indicate interquartile range; whiskers extend 1.5 times the interquartile range) of mean observed/expected *cis* Hi-C contact frequencies for MD-MD contacts for control (sgNT) and NSD3-depleted (sgNSD3) TC-797 cells show a significant reduction of *cis* MD-MD contacts upon NSD3 depletion. Contacts with zero or undefined observed/expected values in any dataset were removed for a total of 78 *cis* MD-MD contacts. *p* values determined using a one-sided Mann-Whitney *U* test. (**H**) Box plots (horizontal lines show medians; boxes indicate interquartile range; whiskers extend 1.5 times the interquartile range) of mean observed/expected *trans* Hi-C contact frequencies for MD-MD contacts for control (sgNT) and NSD3-depleted (sgNSD3) TC-797 cells show a significant reduction of *trans* MD-MD contacts upon NSD3 depletion. Contacts with zero or undefined observed/expected values in any dataset were removed for a total of 659 *trans* MD-MD contacts. *p* values determined using a one-sided Mann-Whitney *U* test. (**I**) For each *trans* MD-MD contact (n = 659), the fold-change in mean observed/expected Hi-C contact frequency versus the observed MD-MD contact frequency in control (sgNT) TC-797 cells. The color of data points reflects the rank of the observed *trans* MD-MD contact frequency from least to greatest in control (sgNT) TC-797 cells. (**J**) For each *cis* MD-MD contact (*n* = 78), the fold-change in mean observed/expected Hi-C contact frequency versus the observed MD-MD contact frequency in control (sgNT) TC-797 cells. The color of data points reflects the rank of the observed *cis* MD-MD contact frequency from least to greatest in control (sgNT) TC-797 cells. (**K**) For each *cis* MD-MD contact (*n* = 78), the fold-change in mean observed/expected Hi-C contact frequency versus genomic distance separating the interacting megadomains. The color of data points reflects the rank of the observed *cis* MD-MD contact frequency from least to greatest in control (sgNT) TC-797 cells.

**Figure S9.**
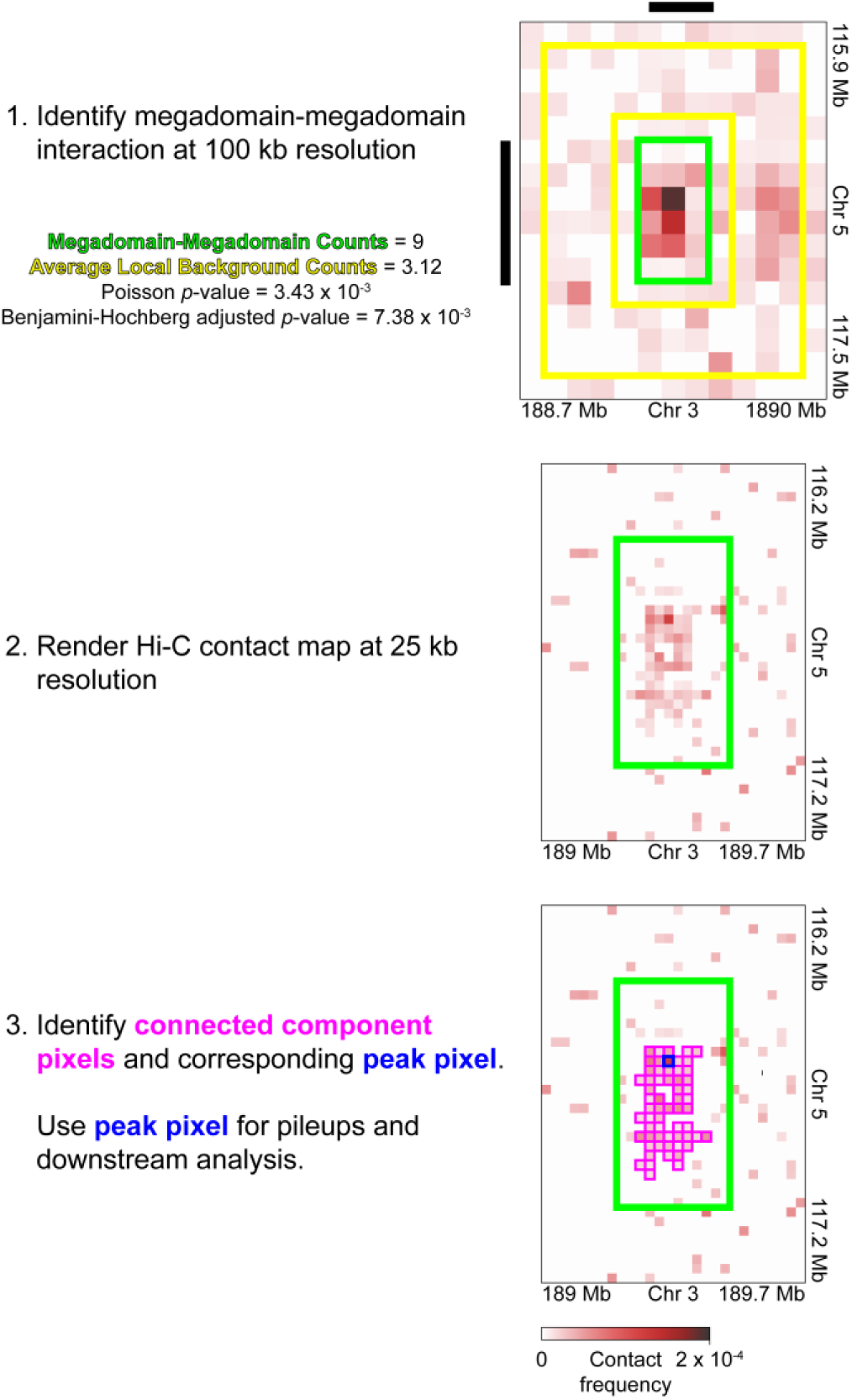
Peak pixel extraction from megadomain-megadomain interactions. (1) Megadomain-megadomain interactions were annotated at 100 kb resolution as described in the Methods section and previously^34^. (2) To identify the peak pixel of each megadomain-megadomain interaction, the Hi-C contact map was rendered at 25 kb resolution. (3) First, the edge-adjacent-only connected pixels were identified. If multiple clusters of connected pixels were identified, the connected pixels contributing to the cluster with the greatest number of pixels were identified as “connected component pixels”. Lastly, the pixel with the highest contact frequency among the connected component pixels was extracted as the peak pixel and was used for pileup and downstream analysis.

**Figure S10.**
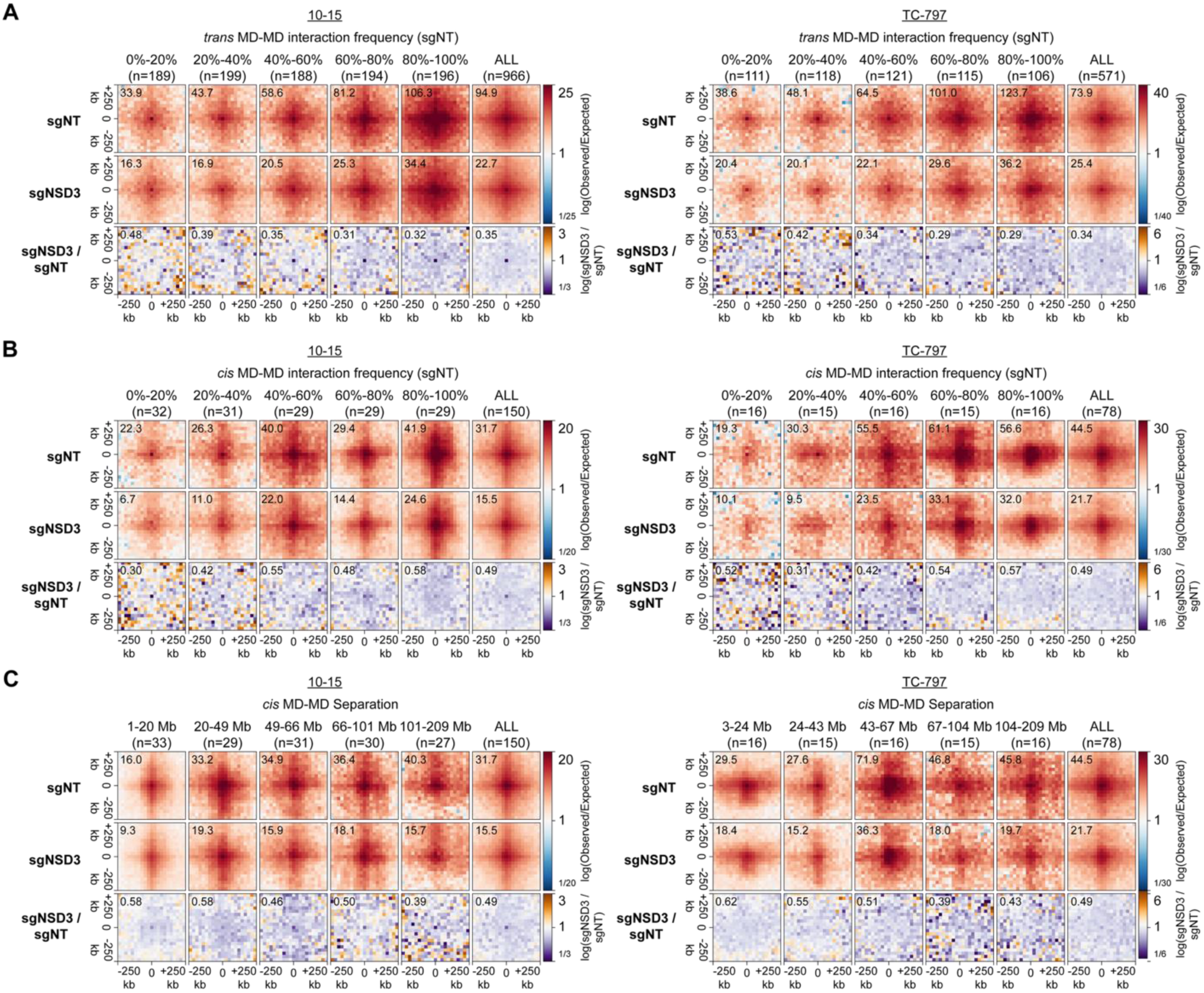
Additional pileups of *cis* and *trans* megadomain-megadomain (MD-MD) interactions. (**A**) Genome-wide pileups of mean observed/expected *trans* Hi-C contact frequencies at MD-MD contacts categorized by observed *trans* MD-MD contact frequencies in control (sgNT) cells for control (sgNT) and NSD3-depleted (sgNSD3) 10-15 and TC-797 cells. Below the observed/expected pileups are pileups of the fold-change for NSD3-depleted (sgNSD3) versus control (sgNT) cells. The value of the central pixel is given at the top left. Windows extending beyond chromosome ends and contacts with zero or undefined observed/expected values in any dataset were excluded from pileups. (**B**) Genome-wide pileups of mean observed/expected *cis* Hi-C contact frequencies at MD-MD contacts categorized by observed *cis* MD-MD contact frequencies in control (sgNT) cells for control (sgNT) and NSD3-depleted (sgNSD3) 10-15 and TC-797 cells. Below the observed/expected pileups are pileups of the fold-change for NSD3-depleted (sgNSD3) versus control (sgNT) cells. The value of the central pixel is given at the top left. Windows extending beyond chromosome ends and contacts with zero or undefined observed/expected values in any dataset were excluded from pileups. (**C**) Genome-wide pileups of mean observed/expected *cis* Hi-C contact frequencies at MD-MD contacts categorized by genomic distance separating the interacting megadomains for control (sgNT) and NSD3-depleted (sgNSD3) 10-15 and TC-797 cells. Below the observed/expected pileups are pileups of the fold-change for NSD3-depleted (sgNSD3) versus control (sgNT) cells. The value of the central pixel is given at the top left. Windows extending beyond chromosome ends and contacts with zero or undefined observed/expected values in any dataset were excluded from pileups.

**Figure S11.**
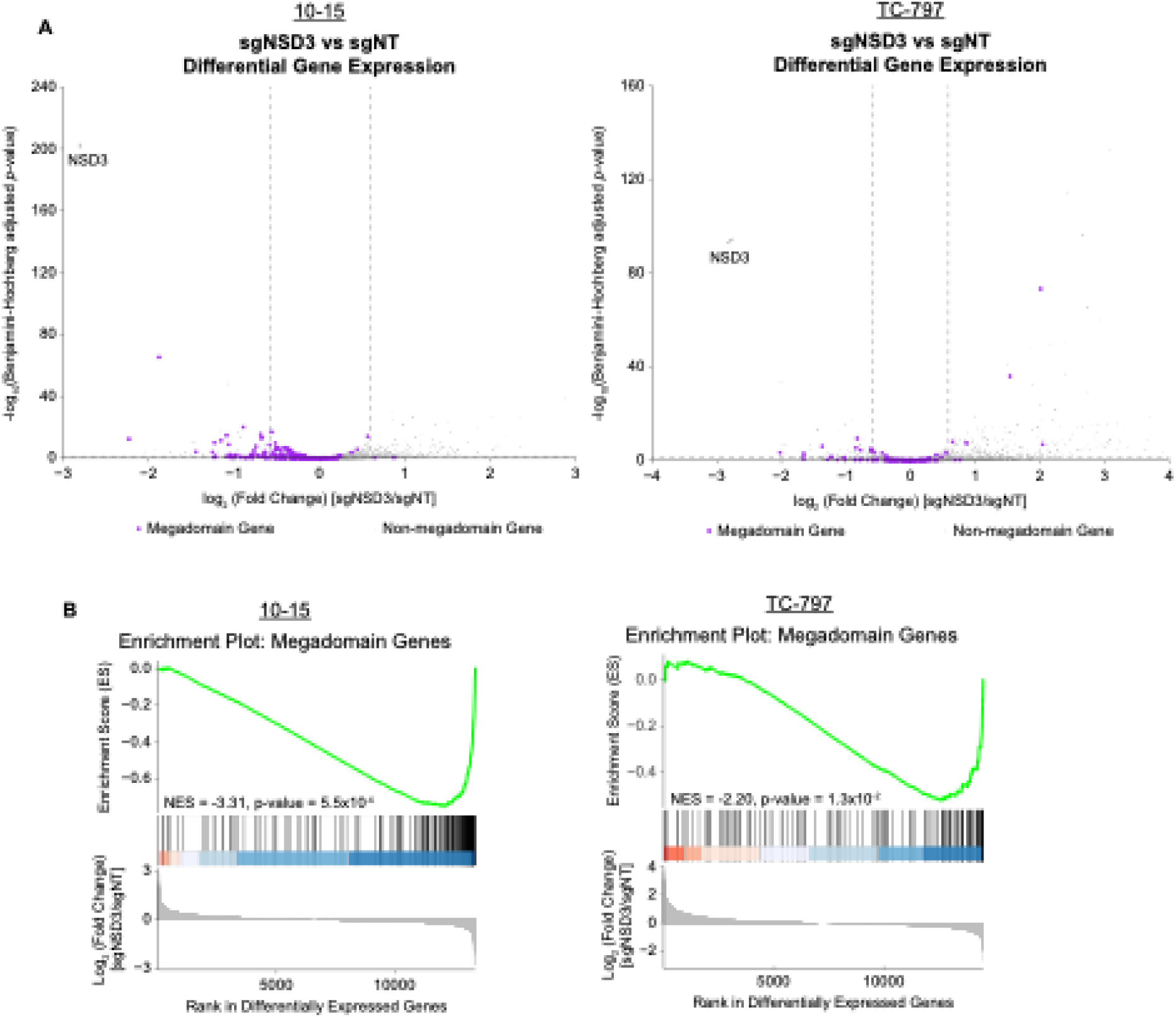
Depletion of NSD3 downregulates genes within megadomains. (**A**) Volcano plots (significance versus fold change) of differentially expressed genes show that after depletion of NSD3 the majority of genes within megadomains (purple dots) are downregulated in 10-15 cells and TC-797 cells. Horizontal dashed lines indicate a Benjamini-Hochberg adjusted p value of 0.05; dashed vertical lines indicate a 1.5-fold change in expression. (**B**) Gene Set Enrichment Analysis (GSEA) indicates that genes within megadomains are downregulated upon depletion of NSD3 (sgNSD3) compared to control (sgNT) in 10-15 and TC-797 cells. NES, normalized enrichment score.

**Figure S12.**
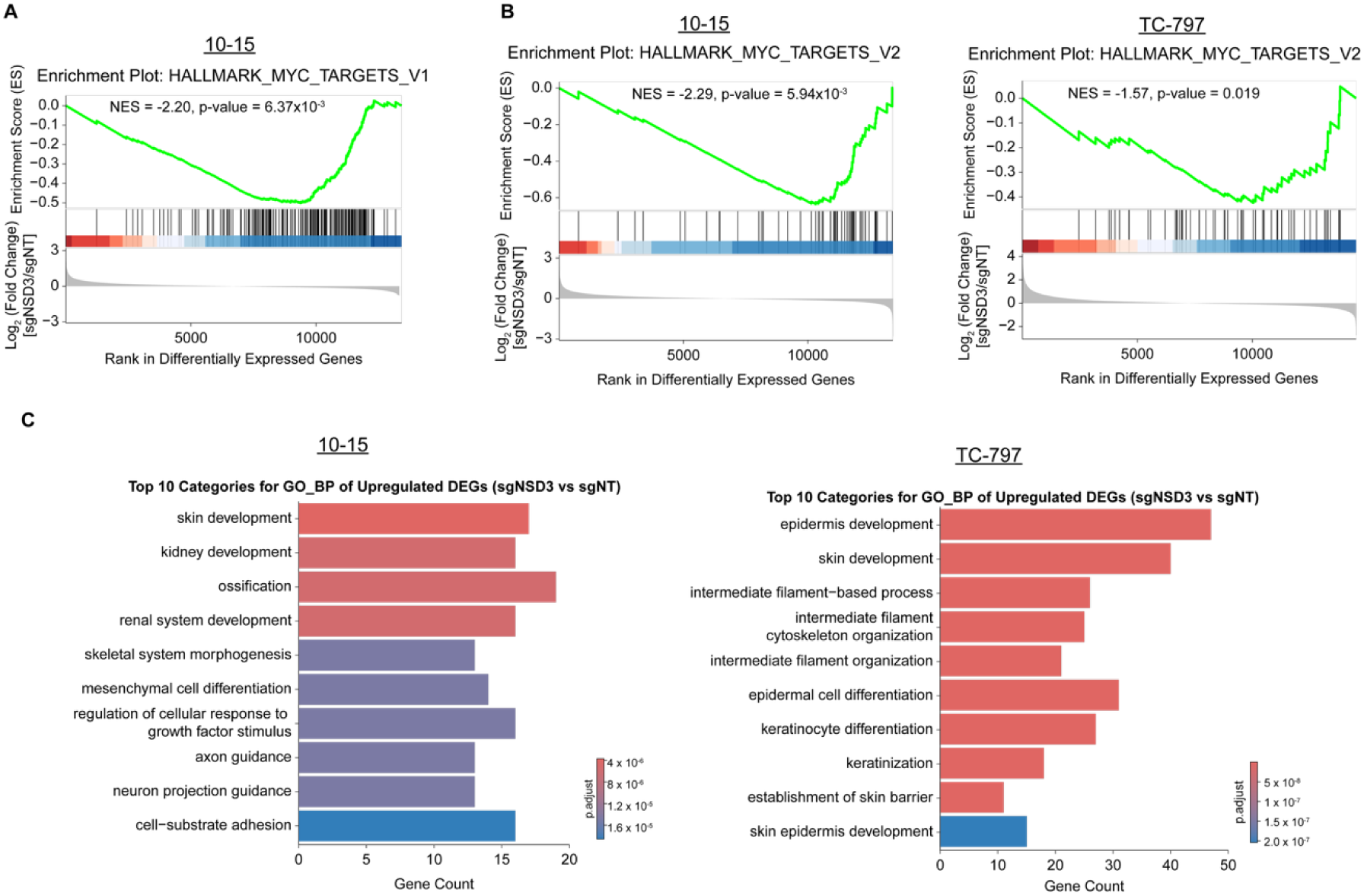
Depletion of NSD3 downregulates MYC target genes and upregulates genes linked to biological processes controlling cellular differentiation. (**A**) Gene Set Enrichment Analysis (GSEA) indicates that genes within the HALLMARK_MYC_TARGETS_V1 gene set are downregulated upon depletion of NSD3 (sgNSD3) compared to control (sgNT) in 10-15 cells. NES, normalized enrichment score. (**B**) Gene Set Enrichment Analysis (GSEA) indicates that genes within the HALLMARK_MYC_TARGETS_V2 gene set are downregulated upon depletion of NSD3 (sgNSD3) compared to control (sgNT) in 10-15 cells and TC-797 cells. NES, normalized enrichment score. (**C**) Bar chart of the top ten enriched Gene Ontology Biological Processes (GO BP) within upregulated differentially expressed genes upon depletion of NSD3 from 10-15 cells and TC-797 cells. The length of each bar represents the number of enriched genes in each term. The color scale represents the Benjamini-Hochberg adjusted p value. Terms are ranked by statistical significance of enrichment.

**Figure S13.**
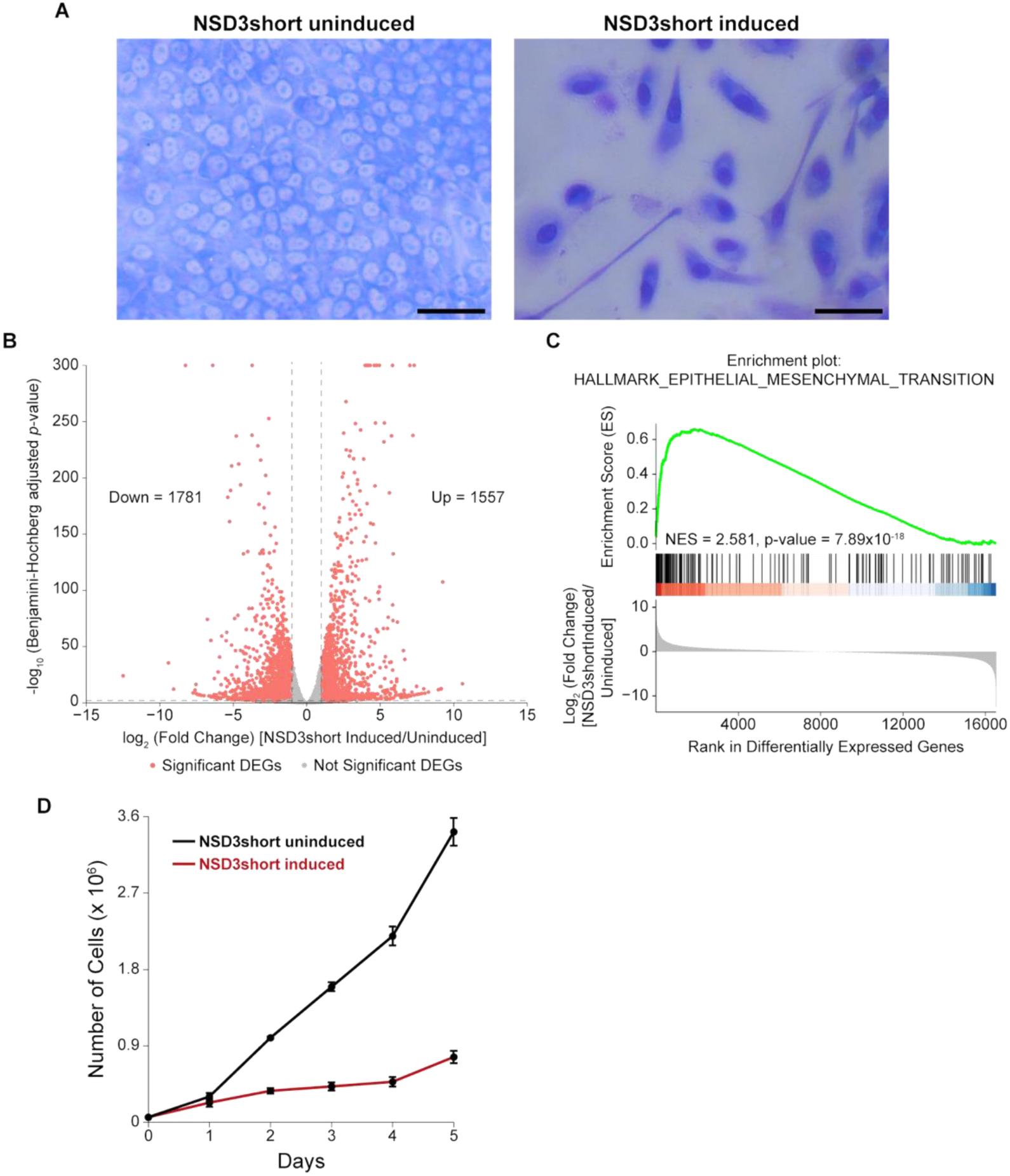
Cellular and molecular phenotypic analyses of NSD3short gain-of-function system in MCF10A cells. (**A**) Photomicrographs of uninduced MCF10A cells and cells induced to express NSD3short show that NSD3short induces cells to transition from an epithelial-like to a mesenchymal-like morphology. Cells were treated with 30 ng/mL doxycycline or water for 5 days followed by Wright-Giemsa staining. Scale bars, 50 µm. (**B**) Volcano plots (significance versus fold change) of differentially expressed genes upon induced expression of NSD3short in MCF10A cells. Salmon dots are significant differentially expressed genes (DEGs); gray dots are genes whose expression does not significantly change. Horizontal dashed lines indicate a Benjamini-Hochberg adjusted p value of 0.01; dashed vertical lines indicate a 2-fold change in expression. (**C**) Gene Set Enrichment Analysis (GSEA) indicates that induced expression of NSD3short in MCF10A cells upregulates genes within the HALLMARK_EPITHELIAL_MESENCHYMAL_TRANSITION gene set. NES, normalized enrichment score. (**D**) Count of uninduced MCF10A cells and cells induced to express NSD3short on each day given on the x-axis. Data points and error bars represent the mean and standard deviation, respectively, of three biological replicates.

**Figure S14.**
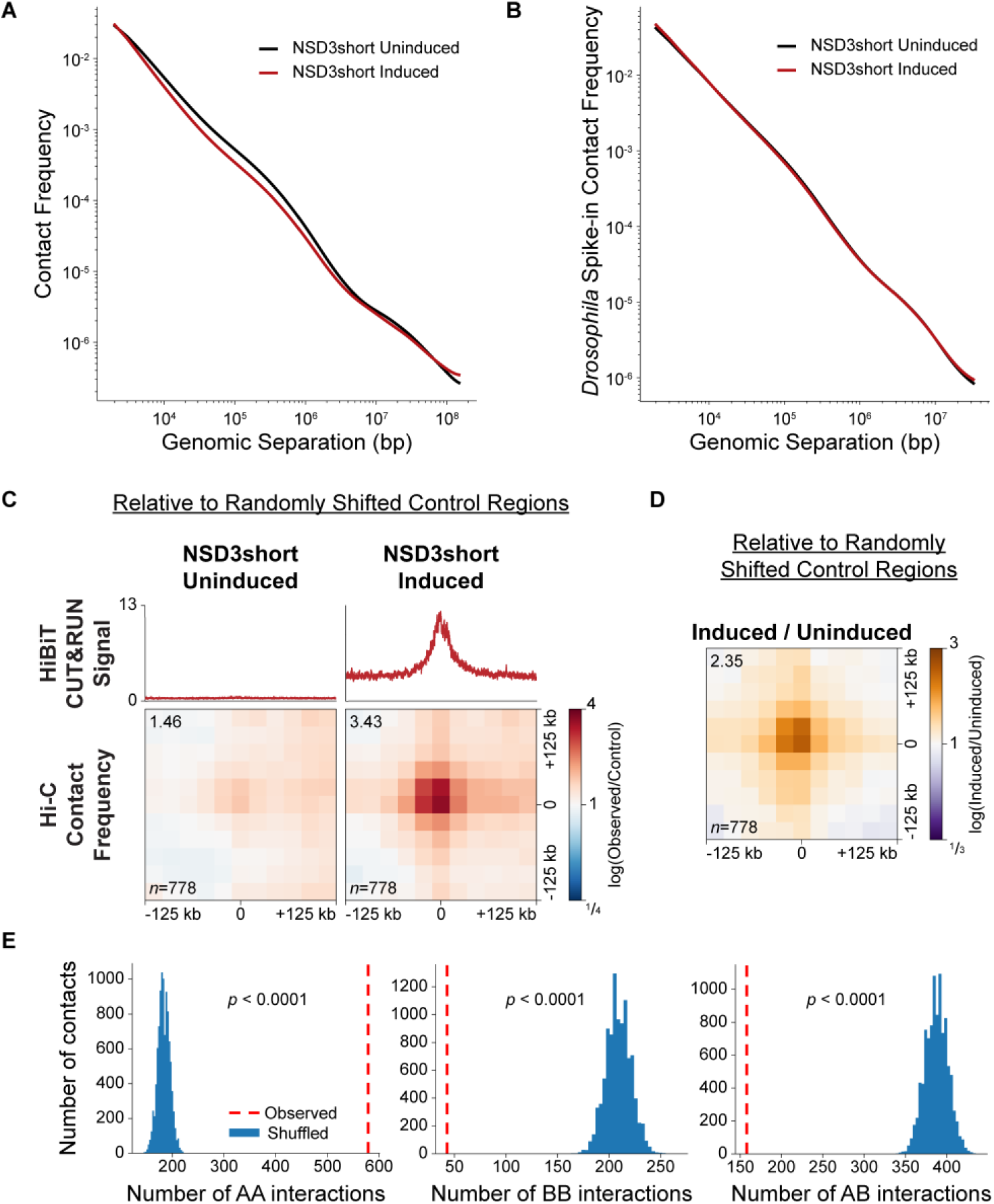
Further analysis of Hi-C data from MCF10A cells expressing NSD3short. (**A**) *P*(*s*) curves of the average intrachromosomal contact frequency separated by the genomic distance on the x-axis in uninduced MCF10A cells (black) and cells induced to express NSD3short (red). (**B**) *P*(*s*) curves of the average intrachromosomal contact frequency for loci from identical amounts of Kc167 *Drosophila melanogaster* cells spiked-in to identical amounts of uninduced (black) and induced (red) MCF10A cells expressing NSD3short separated by the genomic distance on the x-axis. Overlap of the *Drosophila P*(*s*) curves in both conditions at all genomic separations indicates no technical differences in the Hi-C analysis between uninduced and induced cells. (**C**) Genome-wide pileups of mean observed/control Hi-C contact frequencies at NSD3short-dependent chromatin interactions in uninduced MCF10A cells and cells induced to express NSD3short. 100 randomly shifted regions were used as a control. The mean HiBiT CUT&RUN signal at NSD3short contact sites is shown above the pileups. Values of the central pixel are given at top left. (**D**) Genome-wide pileups of the fold-change in mean observed/control Hi-C contact frequencies at NSD3short-dependent chromatin interactions in MCF10A cells induced to express NSD3short versus uninduced cells. 100 randomly shifted regions were used as a control. The value of the central pixel is given at top left. (**E**) Histograms of the overlap of 10,000 random shuffle NSD3short-dependent chromatin contacts with AA, BB, or AB Hi-C compartment interactions. p values were then determined by comparing the observed overlap to the distribution of overlaps using a random permutation test, indicating that the chance of observed NSD3short-dependent chromatin contacts with AA, BB, or AB Hi-C compartment interactions is less than 0.0001 in each case.

**Figure S15.**
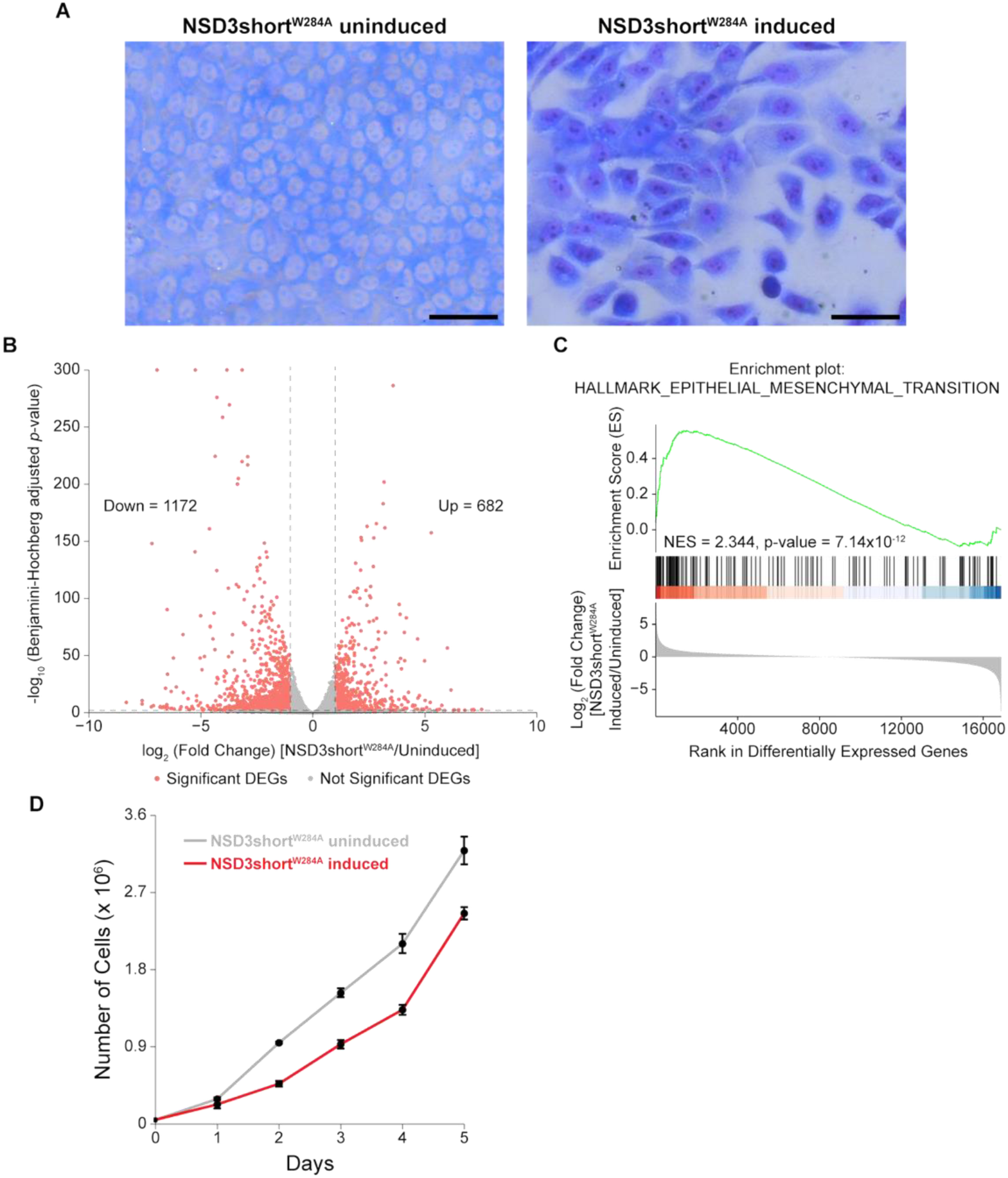
Cellular and molecular phenotypic analyses of MCF10A cells expressing NSD3short^W284A^. (**A**) Photomicrographs of uninduced MCF10A cells and cells induced to express NSD3short^W284A^ show that NSD3short^W284A^ induces a transition from a tightly packed monolayer of epithelial cells to a more loosely packed monolayer. Cells were treated with 25 ng/mL doxycycline or water for 5 days, followed by Wright-Giemsa staining. Scale bars, 50 µm. (**B**) Volcano plots (significance versus fold change) of differentially expressed genes upon induced expression of NSD3short^W284A^ in MCF10A cells. Salmon dots are significant differentially expressed genes (DEGs); gray dots are genes whose expression does not significantly change. Horizontal dashed lines indicate a Benjamini-Hochberg adjusted p value of 0.01; dashed vertical lines indicate a 2-fold change in expression. (**C**) Gene Set Enrichment Analysis (GSEA) indicates that induced expression of NSD3short^W284A^ in MCF10A cells upregulates genes within the HALLMARK_EPITHELIAL_MESENCHYMAL_TRANSITION gene set. NES, normalized enrichment score. (**D**) Count of uninduced MCF10A cells and cells induced to express NSD3short^W284A^ on each day given on the x-axis. Data points and error bars represent the mean and standard deviation, respectively, of three biological replicates.

**Figure S16.**
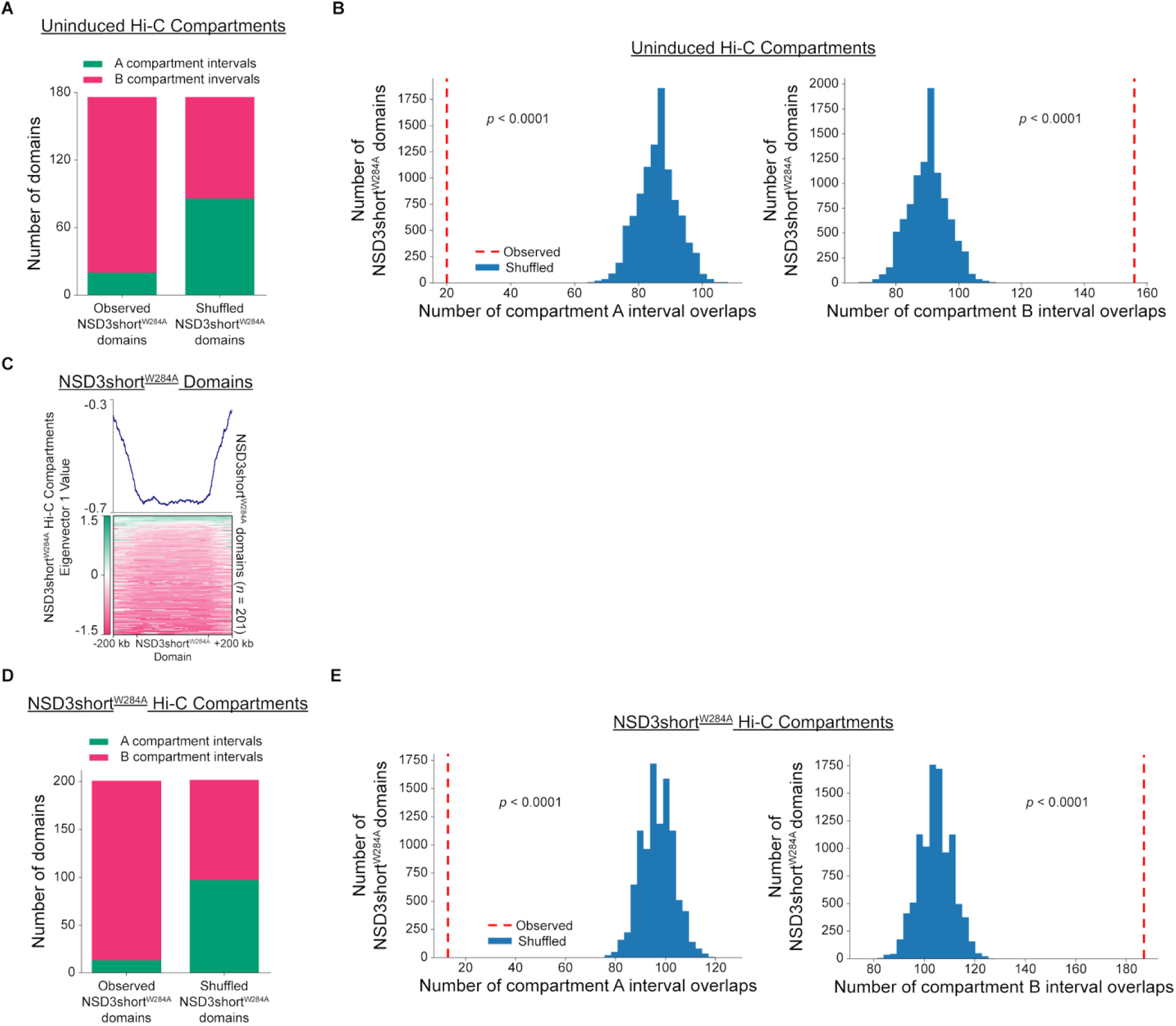
Further analysis of NSD3short^W284A^ domains. (**A**) Number of NSD3short^W284A^ domains overlapping with A or B Hi-C compartment intervals from uninduced MCF10A cells, compared to a random shuffled control set of domains, shows that NSD3short^W284A^ domains arise from the B compartment. (**B**) Histograms of the overlap of 10,000 random shuffle NSD3short^W284A^ domains with A or B Hi-C compartment intervals from uninduced MCF10A cells. p values were then determined by comparing the observed overlap to the distribution of overlaps using a random permutation test, indicating that the chance of observed NSD3short^W284A^ domains overlapping with Hi-C B compartment intervals and not overlapping with Hi-C A compartment intervals is less than 0.0001 in each case. (**C**) Profile (top) and heatmap (bottom) of the Hi-C compartment eigenvector 1 values from MCF10A cells expressing NSD3short^W284A^ within NSD3short^W284A^ domains show that NSD3short^W284A^ domains occupy regions of negative eigenvector 1 values corresponding to B compartment intervals. Profile represents genome-wide mean of the Hi-C compartment eigenvector 1 value. NSD3short^W284A^ domains were normalized to the same length, and 200 kb of flanking DNA is shown next to each normalized NSD3short^W284A^ domain. (**D**) Number of NSD3short^W284A^ domains overlapping with A or B Hi-C compartment intervals from MCF10A cells expressing NSD3short^W284A^, compared to a random shuffled control set of domains, shows that NSD3short^W284A^ domains are found within the B compartment. (**E**) Histograms of the overlap of 10,000 random shuffle NSD3short^W284A^ domains with A or B Hi-C compartment intervals from MCF10A cells expressing NSD3short^W284A^. p values were then determined by comparing the observed overlap to the distribution of overlaps using a random permutation test, indicating that the chance of observed NSD3short^W284A^ domains overlapping with Hi-C B compartment intervals and not overlapping with Hi-C A compartment intervals is less than 0.0001 in each case.

**Figure S17.**
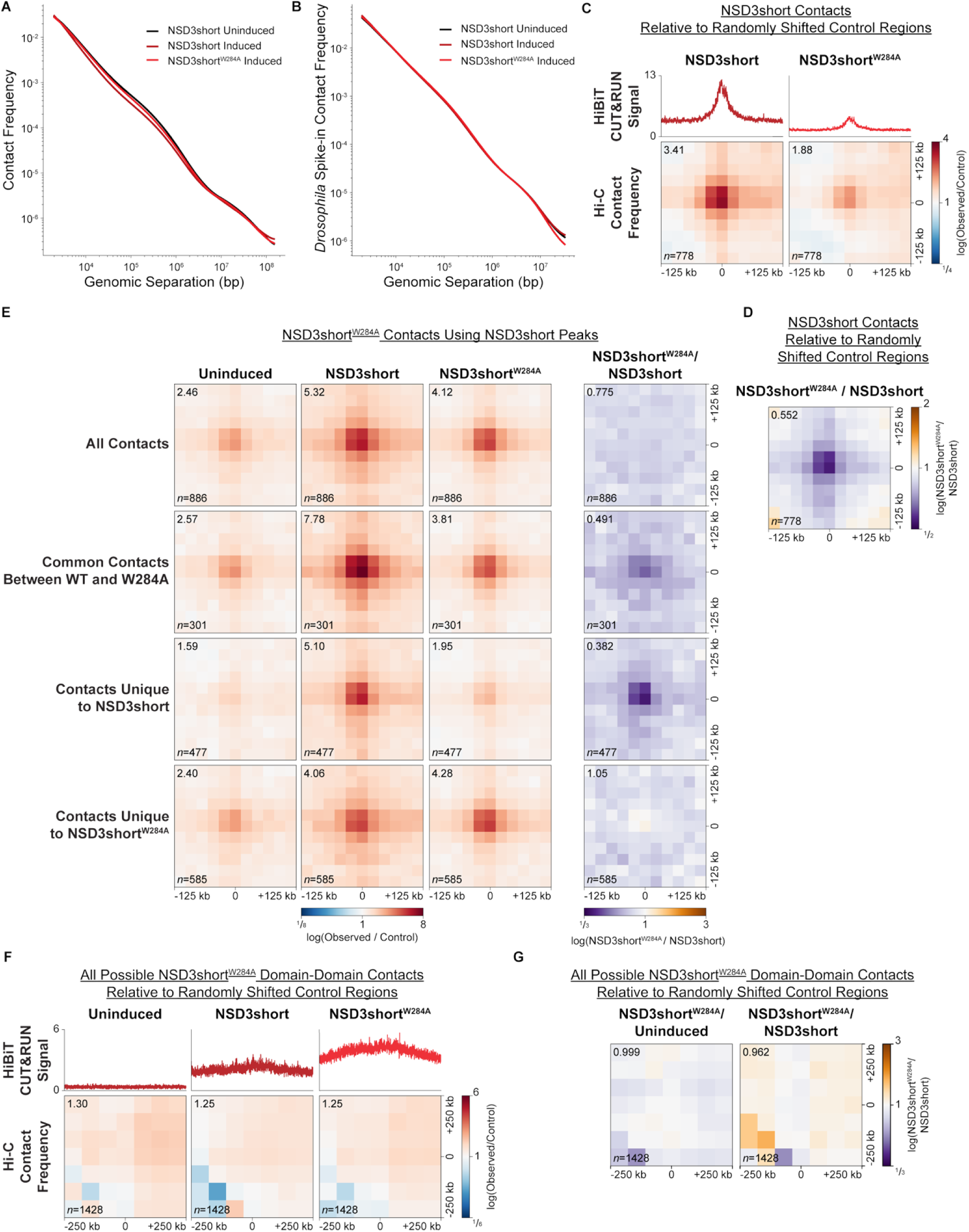
Further analysis of Hi-C data from MCF10A cells expressing NSD3short^W284A^. (**A**) *P*(*s*) curves of the average intrachromosomal contact frequency separated by the genomic distance on the x-axis in uninduced MCF10A cells (black), cells expressing NSD3short (dark red), and cells expressing NSD3short^W284A^ (light red). Data for uninduced MCF10A cells and cells expressing NSD3short are reproduced from Figure S14A. (**B**) *P*(*s*) curves of the average intrachromosomal contact frequency for loci from identical amounts of Kc167 *Drosophila melanogaster* cells spiked-in to identical amounts of uninduced MCF10A cells (black), cells expressing NSD3short (dark red), and cells expressing NSD3short^W284A^ (light red) separated by the genomic distance on the x-axis. Overlap of the *Drosophila P*(*s*) curves in all conditions at all genomic separations indicates no technical differences in the Hi-C analysis between conditions. Data for uninduced MCF10A cells and cells expressing NSD3short are reproduced from Figure S14B. (**C**) Genome-wide pileups of mean observed/control Hi-C contact frequencies at NSD3short-dependent chromatin interactions in MCF10A cells expressing NSD3short ad cells expressing NSD3short^W284A^. 100 randomly shifted regions were used as a control. The mean HiBiT CUT&RUN signal at NSD3short contact sites is shown above the pileups. Values of the central pixel are given at top left. (**D**) Genome-wide pileups of the fold-change in mean observed/control Hi-C contact frequencies at NSD3short-dependent chromatin interactions in MCF10A cells expressing NSD3short and cells expressing NSD3short^W284A^. 100 randomly shifted regions were used as a control. The value of the central pixel is given at top left. (**E**) Genome-wide pileups of mean observed/control Hi-C contact frequencies at NSD3short^W284A^ chromatin contacts in uninduced MCF10A cells, cells expressing NSD3short, and cells expressing NSD3short^W284A^. NSD3short^W284A^ chromatin contacts were identified from Hi-C data from MCF10A cells expressing NSD3short^W284A^ using the locations of the wild-type NSD3short peaks. Values of the central pixel are given at top left. (**F**) Genome-wide pileups of mean observed/control Hi-C contact frequencies at all possible NSD3short^W284A^ domain-domain contacts in uninduced MCF10A cells, cells expressing NSD3short, and cells expressing NSD3short^W284A^ show that NSD3short^W284A^ domains do not interact. 100 randomly shifted regions were used as a control. The mean HiBiT CUT&RUN signal at NSD3short^W284A^ domains is shown above the pileups. Values of the central pixel are given at top left. (**G**) Genome-wide pileups of the fold-change in mean observed/control Hi-C contact frequencies at all possible NSD3shortW284A domain-domain contacts in uninduced MCF10A cells, cells expressing NSD3short, and cells expressing NSD3short^W284A^. 100 randomly shifted regions were used as a control. The value of the central pixel is given at top left.

## SUPPLEMENTARY TABLE LEGENDS

**Table S1. Immunofluorescence Microscopy Mander’s Coefficient Analysis.**

For definitions of each metric see Stirling et al., 2021^63^.

**Table S2. Immunofluorescence Microscopy BRD4-NUT Condensate Analysis in 10-15 Cells.**

For definitions of each metric see Stirling et al., 2021^63^.

**Table S3. Immunofluorescence Microscopy BRD4-NUT Condensate Analysis in TC-797 Cells.**

For definitions of each metric see Stirling et al., 2021^63^.

**Table S4. Hi-C Sequencing Statistics and Quality Metrics.**

For definitions of each metric see Rao et al., 2014^64^.

**Table S5. RNA-seq Differentially Expressed Genes (DEGs) Analysis.**

## Notes

### Competing Interest Statement

The authors have declared no competing interest.

